# Mechanistic insights into redox activity and catalytic determinants of the haloarchaeal flavin-dependent oxidoreductase *Hv*FdR

**DOI:** 10.64898/2026.08.10.743915

**Authors:** Katherine R. Weber, Peter Huynh, Brianna Novillo, Semaj Bulter-Drinks, Christian Heryakusuma, Biswarup Mukhopadhyay, Endang Purwantini, Julie A. Maupin-Furlow

## Abstract

Members of the FAD-dependent oxidoreductase family (IPR050260) play diverse and key roles in maintaining cellular redox balance, yet the functions of many distinct subgroups within this family remain unknown. Here, we define the biochemical and physiological functions of the *Haloferax volcanii* flavin-dependent oxidoreductase *Hv*FdR (HVO_2345; *fdr*), a haloarchaeal member of a previously uncharacterized IPR050260 subgroup. *Hv*FdR binds FAD and catalyzes NAD(P)H oxidase, diaphorase and ferredoxin reductase activities, with a kinetic preference for NADPH over NADH and catalytic properties that are strongly influenced by oxygen availability. Under stoichiometric conditions, *Hv*FdR mediates reverse electron transfer to NADP⁺, suggesting that intracellular nicotinamide nucleotide pools regulate electron flow bidirectionally. Consistent with this reversibility, *Hv*FdR bound-FAD exhibits a low midpoint redox potential (−413 mV), supporting its capacity to function as an electron donor. Deletion of *fdr* impairs growth and elevates intracellular NADPH levels, consistent with a role for *Hv*FdR in maintaining NADP(H) homeostasis. Conserved residues K47 and Y323 are identified as determinants of *Hv*FdR electron transfer activity and may function as a regulatory gate that modulates electron flow while limiting excessive H_2_O_2_ production under aerobic conditions. Together, these findings establish *Hv*FdR as an oxygen-responsive flavin-dependent oxidoreductase that contributes to cellular redox homeostasis and provides functional insight into a previously uncharacterized subgroup of the IPR050260 family.

## Introduction

Flavins, including flavin mononucleotide (FMN) and flavin adenine dinucleotide (FAD), are essential cofactors synthesized from riboflavin (vitamin B2) through evolutionarily ancient biosynthetic pathways [1–4]. Flavoenzymes utilize FMN and FAD cofactors to mediate biological electron transfer [5], with approximately 75% of flavoproteins binding FAD and 25% binding FMN [6], underscoring the predominant role of FAD-dependent enzymes in cellular bioenergetics. The functional versatility of flavins arises from the isoalloxazine ring, which undergoes reversible one- and two-electron redox reactions by cycling among three distinct redox states: the oxidized quinone, one-electron reduced semiquinone, and two-electron reduced hydroquinone, each with characteristic spectroscopic properties [7–10]. This redox flexibility enables flavoenzymes to catalyze a wide range of reactions fundamental to cellular metabolism, including oxidase, monooxygenase, dehydrogenase, ferredoxin-dependent, and electron bifurcation processes [11–15].

Oxygenase-coupled NAD(P)H-dependent ferredoxin reductases (ONFRs) and FAD-dependent NAD(P)H oxidases cluster to the FAD-dependent oxidoreductase families IPR050446 and IPR050260, respectively. Both flavoenzymes oxidize NAD(P)H, but differ in their primary physiological roles. ONFRs transfer electrons from NAD(P)H to ferredoxins, thereby supporting cytochrome P450-catalyzed reactions [16–20]. In contrast, FAD-dependent NAD(P)H oxidases oxidize nicotinamide cofactors and reduce molecular oxygen to H_2_O_2_ or H_2_O, contributing to cellular redox homeostasis under aerobic conditions [21–26]. Notably, certain NAD(P)H oxidases can also transfer electrons to ferredoxins, suggesting that these enzymes may overlap ONFR activities and serve multifunctional roles in regulating intracellular electron flow and redox metabolism [27–30].

Comparative structural studies provide insight into conserved residues important for ONFR and NAD(P)H oxidase catalytic activity, as well as cofactor binding [17, 18, 31]. Similarly to well-characterized flavoproteins of the ferredoxin-NADP^+^ reductase family (IPR033892), ONFRs and NAD(P)H oxidases have an N-terminal domain that binds FAD and a Rossmann-like fold C-terminal domain that binds NAD(P)H and includes conserved glycine-rich motifs that stabilize nucleotide cofactors and facilitate hydride transfer [32–35]. Additionally conserved are residues at the flavin and nicotinamide electron transfer interfaces, including a lysine residue near the N-terminus that interacts with the N5 and O4 atom of FAD, and an aromatic residue near the C-terminus that may stabilize the isoalloxazine ring [17, 18, 31]. Analogous residues in related reductases are suggested to be utilized for hydride transfer between NAD(P)H, flavin, and other electron transfer cofactors [20, 36, 37]. Despite the availability of crystal structures for ONFRs, NAD(P)H oxidases, and related homologs [16–18, 36, 38], direct experimental validation linking conserved residues to electron transfer kinetics remains limited.

Haloarchaea provide an ideal model for studying flavin-dependent electron transfer due to their adaptation to hypersaline environments, where fluctuating oxygen availability and the energetic constraints of the “salt-in” osmoadaptation strategy challenge redox metabolism and protein stability [39–41]. Despite the importance of flavin-dependent oxidoreductases in cellular electron transfer, their functions remain poorly characterized in haloarchaea. Here, we characterized *Hv*FdR (HVO_2345), a flavin-dependent oxidoreductase from the haloarchaeon *Haloferax volcanii* that belongs to a previously uncharacterized archaeal and bacterial subgroup within the IPR050260 family. Biochemical, spectroscopic, structural modeling, and site-directed mutagenesis approaches reveal that *Hv*FdR is an FAD-binding multifunctional redox enzyme with NAD(P)H oxidase, diaphorase, and ferredoxin reductase activities. *Hv*FdR activity and electron transfer properties are strongly influenced by oxygen availability, and conserved residues (K47 and Y323) involved in flavin coordination, electron transfer efficiency, and reactive oxygen species production were identified. Together, these findings establish *Hv*FdR as a versatile flavin-dependent enzyme that integrates oxidase and ferredoxin reductase activities, providing mechanistic insight into the regulation of electron flow and redox homeostasis in haloarchaea under extreme environmental conditions.

## Results

### Flavin-dependent oxidoreductase homolog *Hv*FdR and its associated operon organization

*H. volcanii Hv*FdR (HVO_2345) is a 433 aa protein of the flavin-dependent pyridine nucleotide oxidoreductase (IPR050260) family. *Hv*FdR is composed of an N-terminal FAD/NAD(P)-binding domain (residues 4–279) and a C-terminal NAD^+^–rubredoxin oxidoreductase-like domain (residues 333–386), the latter sharing similarity with enzymes involved in oxygen detoxification and redox balance under low-oxygen conditions [42, 43]. The *fdr* gene encoding *Hv*FdR is transcribed as a leaderless mRNA from genomic position 2,211,216 [44] and is positioned downstream of *hvo_2344* (**Supplemental Fig. S1**), encoding an ArsR-like helix-turn-helix (HTH) domain protein that may act as a redox-sensitive transcription factor via cysteine-mediated metal coordination and disulfide bond formation (**Supplemental Fig. S2**). Small non-coding RNAs are also prevalent in the *fdr* region including those transcribed under oxidative and non-oxidative stress conditions [45] (**Supplemental Fig. S1**). In some *Haloferax* species, an additional ORF encoding a putative transmembrane protein (*e.g*., *Haloferax gibbonsii* LR2-5 HfgLR_11780) is predicted between the *fdr* and HTH-encoding genes (**Supplemental Fig. S1**) [46], suggesting potential species-specific variation in electron transfer or redox-associated functions.

### Sequence network analysis of *Hv*FdR

To investigate the evolutionary relationships and potential functions of *Hv*FdR and its homologs, a sequence similarity network (SSN) and a genome neighborhood network (GNN) were constructed using 898 unique protein sequences identified by homology to HVO_2345 through the BLAST retrieval function of the Enzyme Function Initiative (EFI) tools [47]. Among these proteins, 517 were from Archaea (*Methanobacteriati* including haloarchaea and methanogens), and 477 were from Bacteria (including *Bacillati* and *Pseudomonadati*) (**Supplemental Fig. S3A**). However, neither *Hv*FdR nor its homologs have assigned EC numbers or experimentally characterized functions. Clustering of the SSN generated using EFI-EST alignment score thresholds corresponding to 35–40% pairwise amino acid sequence identity revealed *Hv*FdR to be most closely related to archaeal proteins (**Supplemental Fig. S3B**). Further analysis of the GNN corresponding to the 35% identity cluster, composed of 483 proteins including *Hv*FdR, identified conserved genomic neighborhoods containing genes encoding cysteine-rich HTH domain proteins, as well as proteins annotated as members of DUF6149, DUF7124, DUF5815, DUF7979 and the flavoprotein amino oxidase PF01593 families (**Supplemental Fig. S3C**). Although conservation of genomic synteny alone does not establish a direct functional relationship, the repeated co-localization of these genes across diverse archaeal genomes suggests that they may participate in related biological processes or be subject to shared regulatory constraints. Together, these results indicate that *Hv*FdR belongs to a subgroup of uncharacterized archaeal flavoprotein homologs encoded in genomic synteny with other uncharacterized proteins, many of which contain domains of unknown function (DUFs).

### Comparative protein domain architecture of *Hv*FdR

The protein domain architecture of *Hv*FdR was compared to proteins of known catalytic function using Pfam-based classification. *Hv*FdR contains a N-terminal pyridine nucleotide-disulfide oxidoreductase domain (PF07992) fused to a C-terminal rubredoxin NAD^+^ reductase domain (PF18267) (**Supplemental Fig. S4A**). The PF18267 domain is commonly associated with assimilatory nitrite reductase complexes and reactive oxygen species (ROS) scavenging enzymes, whereas PF07992 is found in NAD(P)H oxidases, ONFRs, and related enzymes. Analysis of other PF07992-containing proteins showed that the NAD(P)H oxidases possess an additional dimerization domain (PF02852), while ONFRs contain a C-terminal reductase domain (PF14759); both of which are absent in *Hv*FdR. Additional analysis identified NAD(P)H oxidizing flavoproteins that share the tandem PF07992-PF18267 architecture of *Hv*FdR, including the tRNA ligase complex-associated NAD(P)H dehydrogenase PYROXD1 from *Homo sapiens* [48] and the NADH-rubredoxin oxidoreductase NROR of *Clostridium acetobutylicum* [49] (**Supplemental Fig. S4A**). However, *Hv*FdR exhibits only 20-22% amino acid sequence identity with these characterized flavoproteins (**Supplemental Fig. S4B**). Moreover, *Hv*FdR lacks cysteine residues entirely, suggesting that its classification as a disulfide-based oxidoreductase is overly broad.

### 3D-structural modeling and amino acid sequence alignment of *Hv*FdR to proteins of known catalytic function

To further predict the structure and function of *Hv*FdR, 3D-modeling and amino acid sequence alignments were performed with proteins of known catalytic function. *Hv*FdR was found to exhibit 3D-structrual homology (**Supplemental Table S1**) and to share conserved primary amino acid sequence motifs (**Supplemental Fig. S5**) with NAD(P)H oxidases and ONFRs of PF07992. Notably, while *Hv*FdR lacked the conserved redox-active cysteine residue typically required for NAD(P)H oxidase four-electron reduction of O_2_ to H_2_O [30, 50, 51], two glycine-rich motifs (G-X_1–2_-G-X_2_-G-X_3_-G/A) used by flavin-dependent oxidoreductases to bind NAD(P)H and FAD [33, 35, 52] were conserved. Furthermore, *Hv*FdR contains the lysine residue proposed to be key for flavin binding in ONFRs [17, 18, 31]. Collectively, these findings indicate that *Hv*FdR shares structural features with NAD(P)H oxidases, ONFRs, and other related oxidoreductases, while also possessing distinct features in its domain architecture.

### 3D-structural modeling of the putative *Hv*FdR active site

To initiate the investigation of catalytic mechanism, the 3D-structural models of *Hv*FdR were evaluated for cofactor (*i.e*., NAD^+^, NADP^+^, NADPH, and FAD) binding sites and the feasibility of electron transfer between these cofactors. Based on AlphaFold modeling, *Hv*FdR is predicted to bind NADPH and FAD with an approximate spatial separation of 10.5 Å between the redox centers (**Fig. 1A**), positioning the NADPH within a neutrally charged binding pocket and in proximity to the isoalloxazine ring, consistent with canonical hydride transfer mechanisms observed in related flavoproteins [53–56]. In the *Hv*FdR 3D-models, the conserved residues K47 and Y323 were found positioned on opposite sides of the flavin interface and were predicted to contribute to FAD binding and stabilization, as well as electron transfer (**Fig. 1B, left and Fig. 1C**). Electrostatic surface analysis revealed a predominantly negative surface potential, with neutrally charged pockets surrounding the FAD-binding region (**Fig. 1B, right**). Compared to the *Pseudomonas putida* Pdr [16] (**Fig. 1C),** the conserved residues identified in *Hv*FdR clustered around the FAD-binding pocket (**Fig. 1D)**, demonstrating their positioning relative to the isoalloxazine ring. Together, these analyses support classification of *Hv*FdR as a structurally conserved flavin-dependent oxidoreductase capable of mediating electron transfer from NAD(P)H to ferredoxin, oxygen, or alternative electron acceptors.

**Figure 1.**
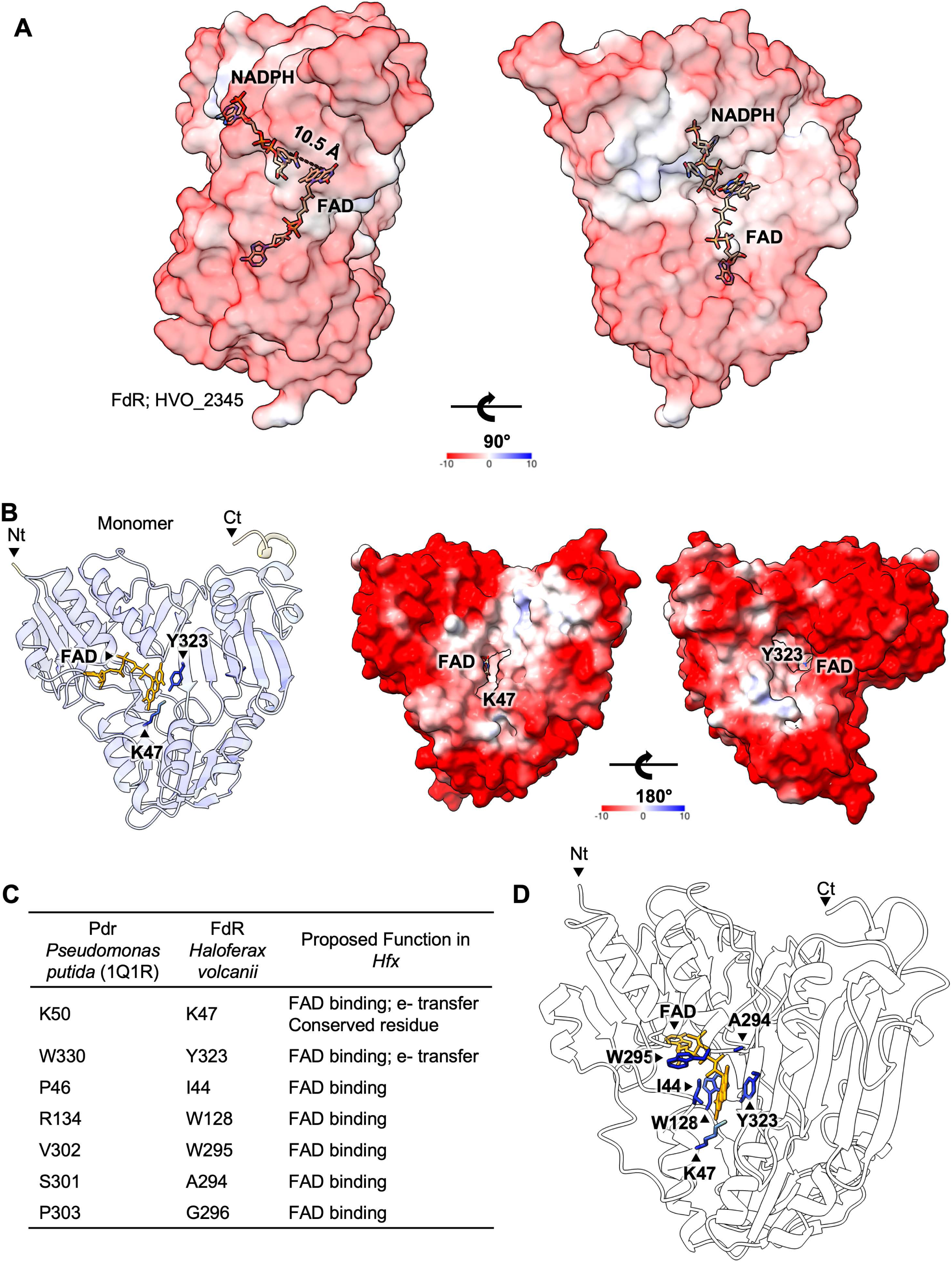
Proposed mechanisms of a ferredoxin reductase (flavin-based oxidoreductase; FdR). A. The schematic depicts the proposed NADPH oxidase two-electron transfer mechanism, where NADPH donates a hydride equivalent (2e^-^) to FAD, which subsequently transfers electrons to downstream acceptors, including molecular oxygen. AlphaFold 3 server model (ipTM = 0.96 and pTM = 0.95) of *Hv*FdR, FAD, and NADPH. Electrostatic surface representation of the ferredoxin reductase colored by surface potential (red, negative; blue, positive; scale shown). The edge-to-edge distance between NADPH and FAD is approximately 10.5 Å, consistent with efficient hydride transfer. Midpoint redox potentials: NADPH, -320 mV; oxygen, +695 mV. A 90° rotated view (right) highlights the spatial alignment of the cofactors within the protein matrix. Electrons are transferred from the C4 carbon of NADPH’s nicotinamide ring to the N5 nitrogen of the FAD isoalloxazine ring via a hydride transfer. From there, electrons are passed one at a time to electron acceptors, such as molecular oxygen or ferredoxins. AlphaFold 3 server model of *Hv*FdR, FAD, and NAD^+^ (ipTM = 0.94 and pTM = 0.95), as well as *Hv*FdR, FAD, and NADP^+^ (ipTM = 0.96 and pTM = 0.95). B. **Left:** FdR (HVO_2345) and FAD AlphaFold 3 server model (ipTM = 0.98 pTM = 0.94) with predicted FAD (yellow), as well as binding residues K47 and Y323 side chains (blue). **Right**: FdR surface charge model with predicted Fdx binding pocket (middle). Positive (blue) and neutral (white) charges on acetylation sites suggest protein-protein interaction sites. C. Comparing Pdr from *P. putida* to FdR from *Hv*. The predicted non-covalent FAD binding site is the conserved residue K47. The aromatic ring of Y323 is predicted to shield the FAD cofactor from solvent and support flavin binding. Both K47 and Y323 are suggested to be important residues for efficient electron transfer between protein partners. D. Side profile of FdR homologues residues predicted to bind FAD (orange) highlighted.

### Purification and oligomeric state of *Hv*FdR

To characterize biochemical properties, *Hv*FdR was expressed with an N-terminal His_6_-tag in *H. volcanii* KT05 (H1207 Δ*fdr*) and purified by Ni-NTA affinity chromatography, followed by size exclusion chromatography. SDS-PAGE analysis of the purified His_6_-*Hv*FdR revealed a single predominant protein band migrating at approximately 60 kDa (**Fig. 2A, Supplemental Fig. S6**). Although His_6_-*Hv*FdR migrated more slowly by SDS-PAGE than expected based on its theoretical molecular mass of 47.9 kDa, the protein is predicted to be highly acidic (pI 4.44) [57]. Acidic proteins can exhibit altered SDS-binding properties, resulting in anomalous electrophoretic mobility [58]. Purified His_6_-*Hv*FdR had a distinct yellow coloration (**Fig. 2B**), consistent with incorporation of a flavin cofactor. SEC analysis revealed His_6_- *Hv*FdR to elute as a single, symmetrical peak (**Fig. 2C, left**) corresponding to an observed molecular mass of 51.5 kDa (**Fig. 2C, right**). Thus, His_6_-*Hv*FdR adopts a monomeric conformation consistent with characterized flavin-dependent oxidoreductases [27, 48, 59–62]. Together, these data indicate His_6_-*Hv*FdR purifies as a soluble, monomeric flavoprotein that retains a yellow color, indicative of a bound flavin cofactor.

**Figure 2.**
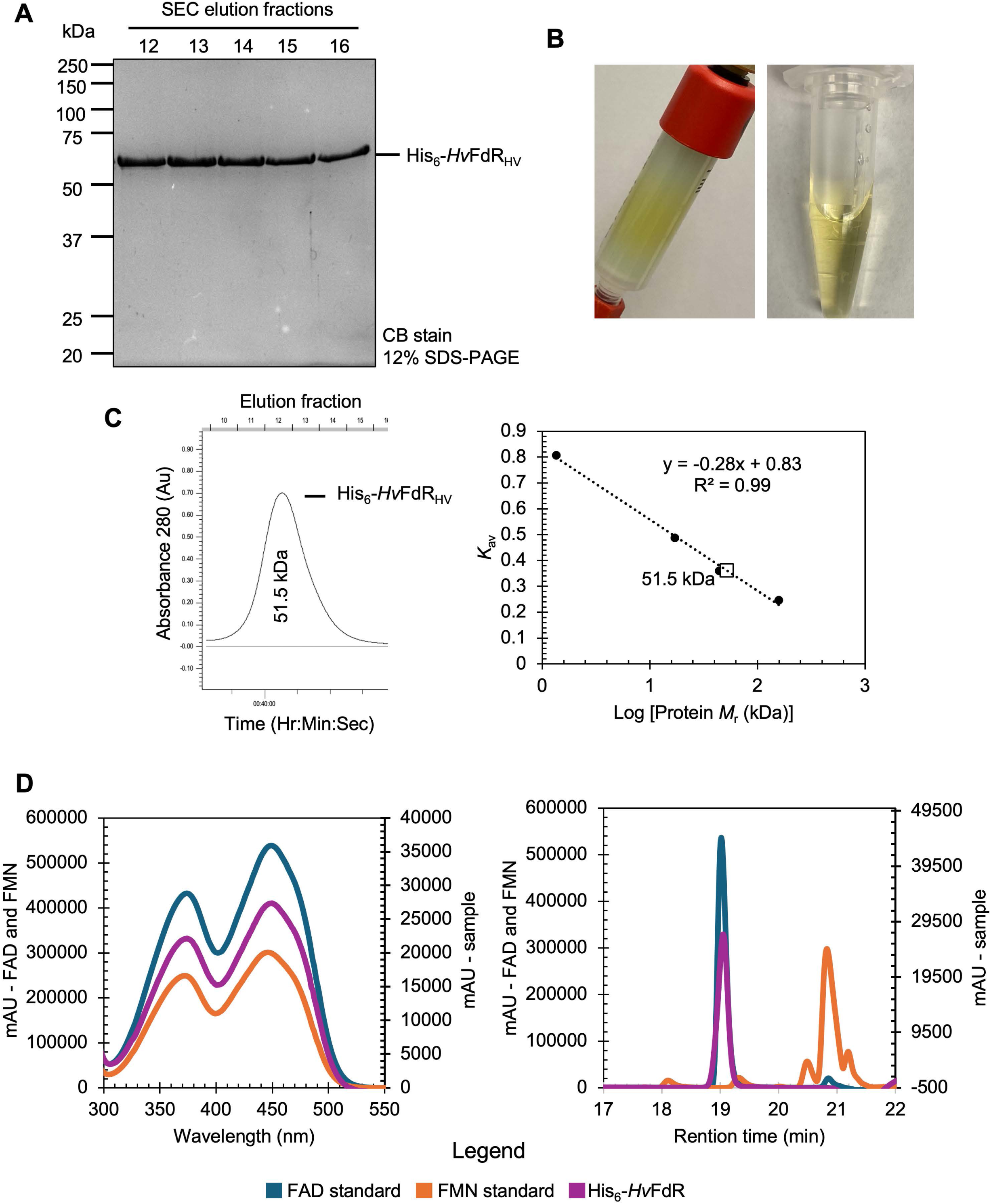
Purity, monomeric state, and flavin-content of His_6_-*Hv*FdR isolated from *H. volcanii*. Strain KT05/pJAM3944 was used to express *Hv*FdR with an N-terminal His_6_- tag using a P2*_rrn_* promoter. Cells were grown to stationary phase (OD_600_ 1.2) on ATCC974 rich medium supplemented with 0.2 µg·mL^-1^ novobiocin. His_6_-*Hv*FdR was purified by Ni-NTA affinity (HisTrap) and Superdex 75 10/300 GL (SEC) chromatography. His_6_-*Hv*FdR fractions were analyzed by SDS-PAGE for purity, by SEC for oligomeric state and by reversed phase high-performance liquid chromatography (RP-HPLC) after methanol-methylene chloride mixture (9:10, v/v) extraction for flavin content (see Methods for details). A. His_6_-*Hv*FdR purifies to apparent homogeneity based on analysis of SEC fractions 12 to 16 (1 µg per lane) by reducing 12% SDS-PAGE and Coomassie blue staining (CB stain). B. His_6_-*Hv*FdR likely binds a flavin cofactor based on the yellow coloration observed during HisTrap chromatography (left) and after elution from the column (right). C. His_6_-*Hv*FdR purifies as a monomer based on SEC. **Left**: SEC chromatogram of His_6_-*Hv*FdR. Fraction 12.39 correlates to an observed molecular weight of 51.5 kDa. **Right**: SEC standard curve. His_6_-*Hv*FdR (open square) displays an observed molecular weight of 51.5 kDa, with a theoretical molecular weight of 47.9 kDa, compared to the standards (closed circles). D. His_6_-*Hv*FdR carries bound FAD, as evidenced by the UV-Vis spectrum of purified protein (**left**) and analysis of extracted cofactor (**right**). **Left**: UV-Visible spectrum of His_6_-*Hv*FdR compared to FAD and FMN standards. Peaks were observed at 374 and 448 nm for the FAD standard (blue), at 371 and 446 nm for the FMN standard (orange), and 374 and 449 nm for His_6_-*Hv*FdR (purple). **Right**: RP-HPLC chromatogram of flavins with X-axis displaying retention time (min) and y-axis in milli-arbitrary unit (mAU) at A_450_. Profiles and peak retention times are displayed for: FAD standard (blue, 19.0 min), FMN standard (orange, 20.8 min), and cofactor extracted from His_6_-*Hv*FdR (purple, 19.0 min) (see Methods for details).

### *Hv*FdR contains FAD as its flavin cofactor

To characterize the putative flavin cofactor bound to His_6_-*Hv*FdR, UV-visible spectral analysis of the purified protein was performed under aerobic conditions. From this analysis, absorbance maxima characteristic of oxidized FAD bound to His_6_-*Hv*FdR were observed. Peaks at 374 nm and 449 nm were detected for His_6_-*Hv*FdR that closely matched those of the FAD standard (374 nm and 448 nm) compared to FMN (371 nm and 446 nm), suggesting the presence of bound FAD (**Fig. 2D, left**).

To further determine the bound cofactor, flavins were extracted from the purified His_6_-*Hv*FdR using a methanol: methylene chloride mixture (9:10, v/v) and analyzed by reversed phase high-performance liquid chromatography (RP-HPLC). Under identical chromatographic conditions, the FAD and FMN standards exhibited distinct retention times, with FAD eluting at 19.0 min and FMN at 20.8 min (**Fig. 2D, right**). The flavin extracted from His_6_-*Hv*FdR exhibited a single dominant peak at 19.0 min, closely matching the FAD standard and distinct from FMN (**Fig. 2D, right**). No detectable peak corresponding to FMN was observed, indicating that His_6_-*Hv*FdR specifically bound FAD.

To determine flavin occupancy, the corresponding FAD peak detected by RP-HPLC analysis was quantified based on the area-under-the-curve (AUC) values. Using this approach, His_6_-*Hv*FdR (6.26 µM protein) contained 4.3 µM bound FAD, representing approximately 68% flavin occupancy when compared to a theoretical 1:1 molar binding. This level of flavin occupancy is consistent with other flavoprotein purifications and reconstitutions [63–66]. Together, UV–visible spectral signatures, RP-HPLC analysis of flavin extracts, and quantitative analysis of flavin occupancy reveal His_6_-*Hv*FdR to bind FAD as its physiological flavin cofactor. Furthermore, successful release of the flavin by organic extraction reveals FAD to be non-covalently associated with *Hv*FdR.

### Reduced growth phenotype of Δ*fdr* mutant strain

A *H. volcanii* Δ*fdr* mutant strain (KT05) was generated by homologous recombination methods [67] and compared to the parent strain (H1207) for growth in ATCC974 nutrient-rich medium. Growth was monitored over 42 h by measuring optical density at 600 nm (OD_600_) and key growth metrics were calculated including AUC, growth rate (h^-1^), doubling time (h) and maximum OD_600_ (**Fig. 3A-B**; **Table 1**). The Δ*fdr* mutant exhibited a growth defect, characterized by an increased doubling time, reduced growth rate, and lower maximum OD_600_ compared to the parent strain (H1207). To assess whether the Δ*fdr* mutation could be complemented, the plasmid expressing His_6_-*Hv*FdR (pJAM3944) was compared to the empty vector control (EV). The parent and Δ*fdr* mutant carrying the empty vector were found to exhibit reduced growth, attributed to plasmid burden, relative to the plasmid-free strains. Importantly, homologous expression of His_6_-*Hv*FdR restored growth of the *Δfdr* mutant to levels exceeding those of both the parent and *Δfdr* mutant strain carrying the empty vector. Collectively, these findings indicate that *Hv*FdR is important for optimal growth under nutrient-rich conditions and that its expression can enhance growth. One possible explanation is that deletion of *fdr* disrupts cellular redox balance, potentially by altering NAD(P)H/NAD(P)^+^ homeostasis.

**Figure 3.**
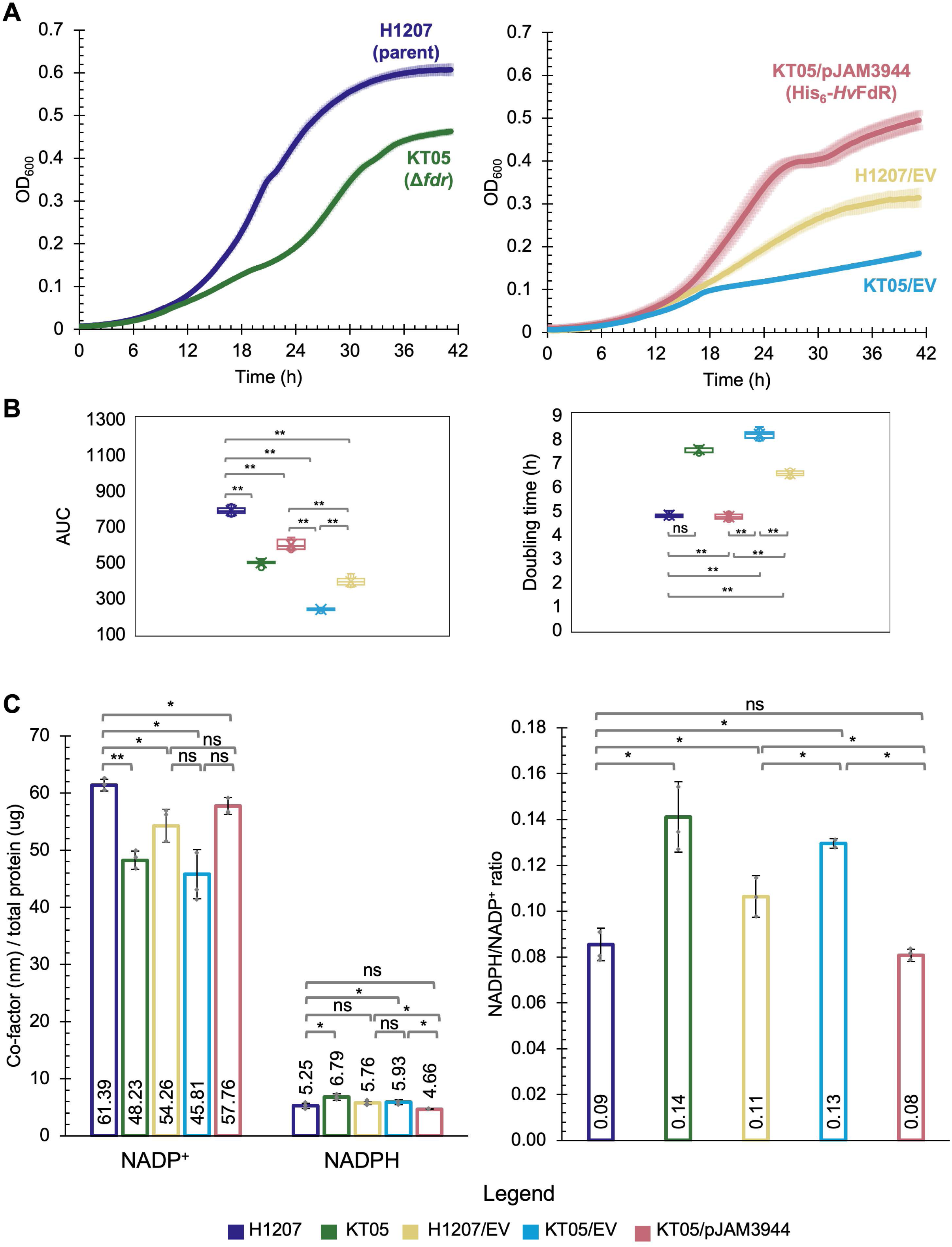
*H. volcanii* Δ*fdr* mutation impairs growth and reduces NADP^+^ levels. *H. volcanii* strains included: H1207 (parent, dark purple), KT05 (H1207 *Δfdr,* green), KT05/pJAM3944 (His_6_-*Hv*FdR, pink), KT05/EV (empty vector, light blue), and H1207/EV (empty vector, yellow). A. Growth curve at 42 °C in ATCC974 rich medium. Growth was then monitored by OD_600_ using EPOCH2 microplate reader and Gen5 software, every 15 min for 99 h, shaking at double orbital (continuously) (see Methods for details). No-inoculum control was used as a blank. Values represent three biological and three technical replicates and are reproducible across experiments. **Left**, OD_600_ measurements of parent vs mutant strains. **Right**, OD_600_ measurements of plasmid complementation strains. B. A student’s *t*-test was used to determine the statistical significance (*p*-value <0.005, **; ns, not significant) of the area under the curve (AUC; **left**) and doubling time (*t_d_*; **right**) calculated over the 42 h time course for H1207 compared to KT05 (*p* = 4.33 x 10^-15^ ** for AUC and 2.64 x 10^-19^** for *t_d_*) H1207/EV (*p* = 1.03 x 10^-16^ ** for AUC and 1.24 x 10^-16^ ** for *t_d_*), KT05/EV (*p* = 6.30 x 10^-15^ ** for AUC and 2.23 x 10^-15^ ** for *t_d_*), and KT05/pJAM3944 (*p* = 1.13 x 10^-10^ ** for AUC and 0.22 for *t_d_*). KT05/pJAM3944 verses KT05/EV (*p* = 5.49 x 10^-11^ ** for AUC and 1.24 x 10^-16^ ** for *t_d_*).H1207/EV verses KT05/EV (*p* = 1.41 x 10^-8^ ** for AUC and 1.90 x 10^-12^ ** for *t_d_*). H1207/EV verses KT05/pJAM3944 (*p* = 2.58 x 10^-11^ ** for AUC and 8.20 x 10^-17^ ** for *t_d_*). C. NADP^+^ and NADPH levels (**left**) and NADPH/NADP^+^ ratio (**right**) quantified in whole cell lysate (see Methods for details). Values represent normalized metabolite levels measured from three biological and three technical replicates and are reproducible across experiments around the same OD_600_. A student’s *t*- test was used to determine the statistical significance (*p*-value <0.005, **; ns, not significant) of H1207 compared to KT05 (*p* = 0.0006** for NADP^+^, 0.02 for NADPH and 0.01 for ratio), H1207/EV (*p* = 0.03 for NADP^+^, 0.063 for NADPH and 0.03 for ratio), KT05/EV (*p* = 0.02 for NADP^+^, 0.04 for NADPH and 0.005 for ratio), and KT05/pJAM3944 (*p* = 0.02 for NADP^+^, 0.37 for NADPH and 0.37 for ratio). KT05 compared to KT05/pJAM3944 (*p* = 0.001** for NADP^+^, 0.02 for NADPH and 0.01 for ratio). H1207/EV verses KT05/EV (*p* = 0.05 for NADP^+^, 0.63 for NADPH and 0.04 for ratio). H1207/EV verses KT05/pJAM3944 (*p* = 0.16 for NADP^+^, 0.02 for NADPH and 0.03 for ratio). KT05/EV verses KT05/pJAM3944 (*p* = 0.03 for NADP^+^, 0.04 for NADPH and 2.51 x 10^-5^ for ratio).

**Table 1.**
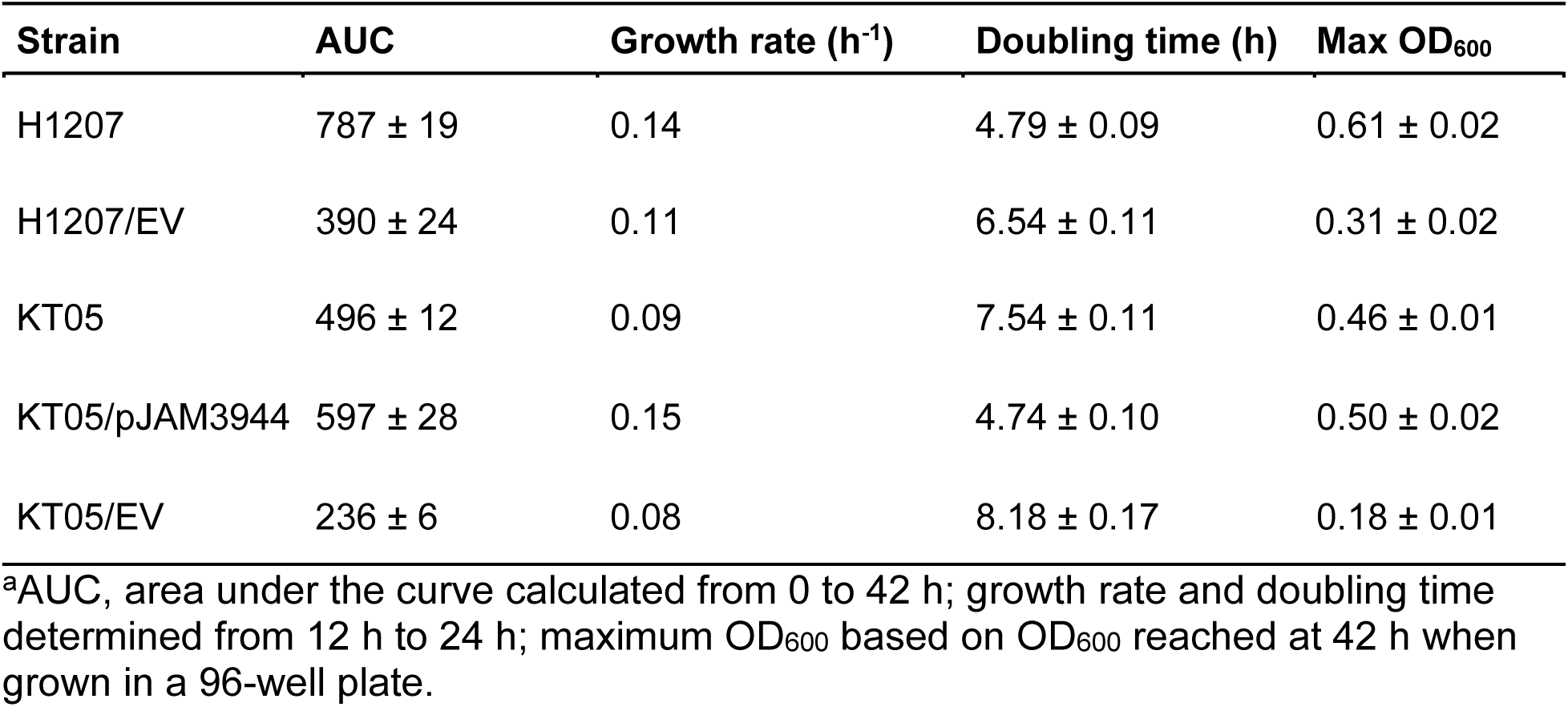
Area under the curve, growth rates, and doubling times of parent (H1207), *Δfdr* mutant (KT05), and complementation (KT05/pJAM3944) strains cultivated in ATCC974 rich medium.

| Strain | AUC | Growth rate (h<sup>-1</sup>) | Doubling time (h) | Max OD<sub>600</sub> |
| --- | --- | --- | --- | --- |
| H1207 | 787 ± 19 | 0.14 | 4.79 ± 0.09 | 0.61 ± 0.02 |
| H1207/EV | 390 ± 24 | 0.11 | 6.54 ± 0.11 | 0.31 ± 0.02 |
| KT05 | 496 ± 12 | 0.09 | 7.54 ± 0.11 | 0.46 ± 0.01 |
| KT05/pJAM3944 | 597 ± 28 | 0.15 | 4.74 ± 0.10 | 0.50 ± 0.02 |
| KT05/EV | 236 ± 6 | 0.08 | 8.18 ± 0.17 | 0.18 ± 0.01 |

<sup>a</sup>AUC, area under the curve calculated from 0 to 42 h; growth rate and doubling time determined from 12 h to 24 h; maximum OD<sub>600</sub> based on OD<sub>600</sub> reached at 42 h when grown in a 96-well plate.

### Deletion of *fdr* disrupts NADPH/NADP^+^ redox balance

To determine whether loss of *fdr* affects cellular redox homeostasis, intracellular NADP^+^ and NADPH levels were quantified under aerobic conditions. Although NADP(H) levels were generally comparable among the strains, the Δ*fdr* mutant exhibited a shift in NADP(H) relative to the parent strain. When NADPH and NADP⁺ pools were analyzed separately, the *Δfdr* mutant resulted in 1.3-fold higher NADPH levels, with a 0.78-fold lower NADP⁺ levels relative to the parent strain (**Fig. 3C left**). Consequently, the NADPH: NADP^+^ ratio was elevated in the *Δfdr* mutant, indicating a more reduced intracellular redox state compared with the parent strain (**Fig. 3C right**). Using a plasmid-based complementation system, introduction of the empty vector control increased the NADPH:NADP⁺ ratio in both the parent and *Δfdr* strains, likely reflecting the metabolic burden associated with plasmid maintenance and expression (**Fig. 3C**). In contrast, plasmid-based expression of His_6_-HvFdR in the Δ*fdr* mutant (KT05/pJAM3944) restored the NADPH:NADP⁺ ratios to levels comparable to the parent strain (H1207) (**Fig. 3C**). Together these results support a role for *Hv*FdR in maintaining cellular redox homeostasis by facilitating NAD(P)H oxidation and demonstrate that reintroduction of His_6_-*Hv*FdR can restore NAD(P)H redox balance in the Δ*fdr* mutant.

### Influence of buffer, pH, salinity, and temperature on *Hv*FdR activity

Given our evidence suggesting that His_6_-*Hv*FdR is active and can mediate electron transfer with NADP(H) *in vivo*, its capacity to function as an NAD(P)H oxidase was further evaluated *in vitro*. This assay evaluated the ability of His_6_-*Hv*FdR to utilize NADPH or NADH as electron donors for oxygen reduction and H_2_O_2_ production under aerobic conditions To determine optimal buffer and pH conditions, the specific activity of His_6_- *Hv*FdR was evaluated over a broad pH range (6–11) using MES, HEPES, Tris, CAPSO, and CAPS buffers (**Fig. 4A**). Both buffer composition and pH differentially affected enzymatic activity depending on the electron donor. NADPH-dependent activity remained relatively stable between pH 6 to 8, whereas NADH-dependent activity progressively decreased from pH 7 to 11. Overall, NADPH-dependent activity was optimal under mildly basic conditions (HEPES buffer at pH 8) (**Fig. 4A, left**), while NADH-dependent activity was favored under acidic conditions (MES buffer at pH 6) (**Fig. 4A, right**).

**Figure 4.**
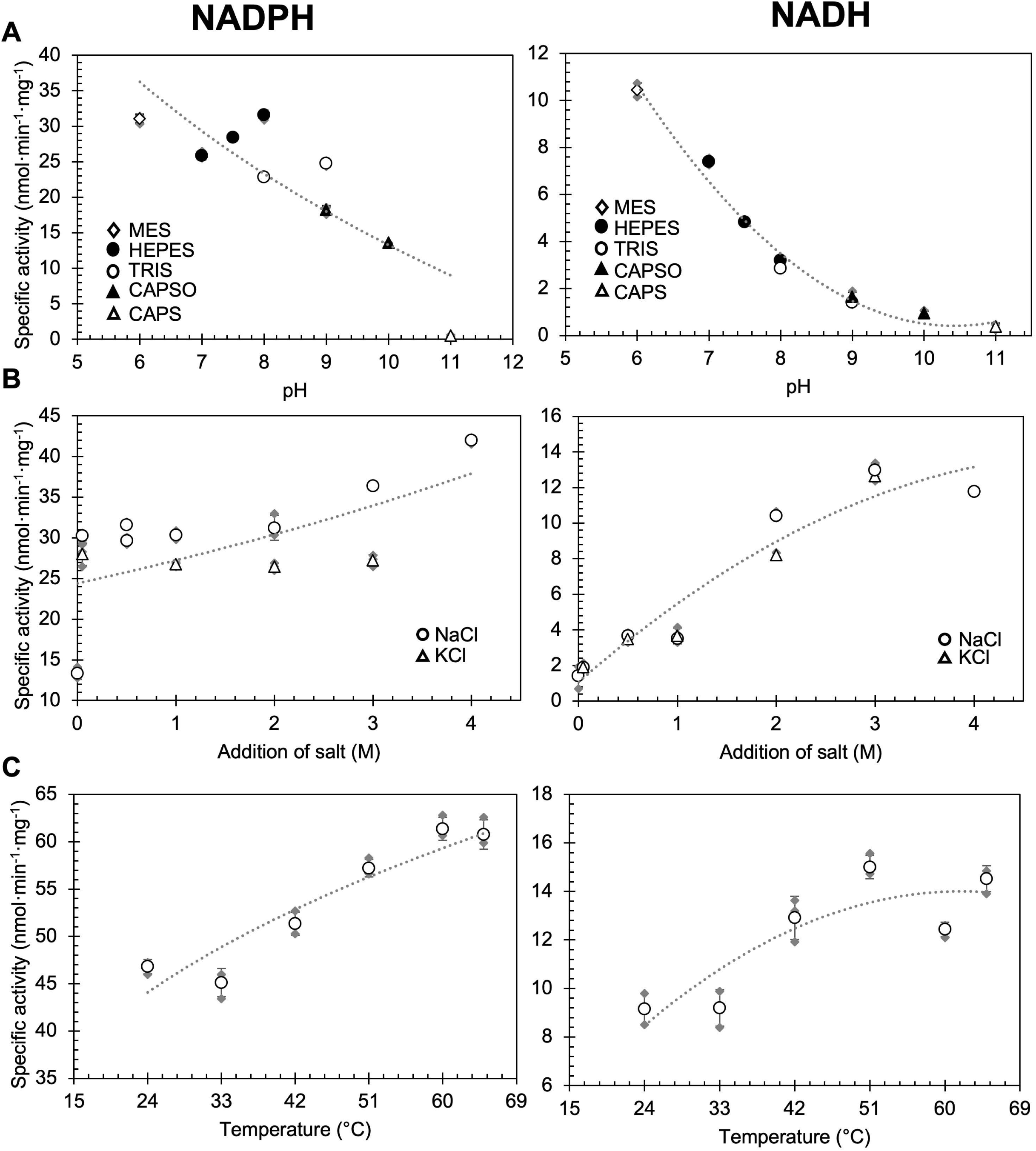
Optimization buffer conditions for FdR oxidase activity. NADPH oxidation measurements were taken at A_340_ over 30 minutes. Specific activity (nmols·min^-1^·mg^-1^) is defined as nmol of NAD(P)H oxidized per minute per milligram of *Hv*FdR. Open circles represent the average of the three replicates and closed grey diamonds represent the individual replicates. In a total reaction volume of 150 µL, 1.5 µM of FdR and 150 µM NADPH (**left**) and 150 µM NADH (**right**) as the electron donor were added under aerobic conditions. Dashed lines represent the best-fit curves generated from the mean specific activity values. Dashed lines represent the best-fit curves generated from the mean specific activity values. A. Optimal pH was tested at 42 °C with 2 M NaCl and 20 mM MES pH 6, HEPES pH 7, HEPES pH 7.5, Tris pH 7.5, HEPES pH 8, Tris pH 8, CAPSO pH 9, Tris pH 9, CAPSO pH 10, or CAPS pH 11. MES, open diamond; HEPES, closed circle; TRIS, open circle; CAPSO, closed triangle; CAPS, open triangle. B. Optimal salinity was tested at 42 °C with 20 mM HEPES [pH 8] for NADPH (**left**) and 20 mM MES [pH 6] for NADH **(right)** with the following addition of NaCl: 0 M, 0.05 M, 0.5 M, 1 M, 2 M, 3 M, and 4 M. Salinity was also tested with the following addition of KCl: 0.05 M, 0.5 M, 1 M, 2 M and 3 M. NaCl, open circles; KCl, open triangles. C. Optimal temperature was tested at 24, 33, 42, 51, 60, and 65 °C with 20 mM HEPES [pH 8] and 4 M NaCl for NADPH (**left**) and 20 mM MES [pH 6] and 3 M NaCl for NADH (**right**).

Optimal salinity for His_6_-*Hv*FdR activity was assessed using the optimal pH and buffer identified for each electron donor (NADPH and NADH). Reactions were supplemented with varying concentrations of NaCl (0.05–4 M) or KCl (0.05–3 M) (**Fig. 4B**). Irrespective of the type of electron donor, His_6_-*Hv*FdR activity was higher in the presence of NaCl compared to KCl and increased with increasing salt concentration. NADPH-dependent activity was maximal at 4 M NaCl (**Fig. 4B, left**), whereas NADH-dependent activity peaked at 3 M NaCl (**Fig. 4B, right**). At lower salt concentrations, enzyme activity was reduced. At 0.05 M NaCl, specific activity was reduced 1.4-fold with NADPH and 6.2-fold with NADH relative to 4 M NaCl. This preference of His_6_-*Hv*FdR for activity in high-salt conditions is consistent with the adaptation of haloarchaeal proteins to the “salt-in” strategy used by haloarchaea in hypersaline environments [68, 69].

Temperature optimization of the reaction was performed under the optimized buffer and salinity conditions (**Fig. 4C**). His_6_-*Hv*FdR exhibited high activity at 51 °C for NADH and 60-65 °C NADPH. Lower activities were observed at 24 °C and 33 °C for both cofactors. The temperatures where His_6_-*Hv*FdR activity was high (51 to 65 °C) were consistent with the optimal growth temperature of *H. volcanii* at 42 to 45 °C [70–72]. Together, these results indicate that the choice of electron donor (NADH vs. NADPH) dictates the optimal biochemical conditions for His_6_-*Hv*FdR activity, with NADPH supporting maximal activity under moderately alkaline, high-salt conditions and high temperature, whereas NADH favors more acidic conditions, slightly reduced salinity, and lower temperature.

### *Hv*FdR concentration optimized to monitor NAD(P)H oxidation activity

To further optimize the assay conditions for NAD(P)H oxidation, different concentrations of His_6_-*Hv*FdR were tested. Across all enzyme concentrations examined, NADPH consistently supported higher specific activity compared to NADH as the electron donor. When NADH was used, lower concentrations of His_6_-*Hv*FdR resulted in increased specific activity. For example, at 250 μM NADH, increasing the enzyme from 1.5 μM to 6 μM resulted in a 1.9-fold reduction in specific activity (**Fig. 5A**), also displaying that excessive enzyme concentrations reduce catalytic efficiency. Additionally, when NADPH was used, the highest specific activity for NADPH oxidation was observed at 1.5 μM His_6_-*Hv*FdR, with lower specific activity detected when 0.5 μM and 2.5 μM of enzyme was used in the assay (**Fig. 5A**). Steady-state kinetic analysis indicated that increasing His_6_-*Hv*FdR concentration also decreased catalytic efficiency (*k*_cat/_*K*_m_) (**Fig. 5B)**. Although increasing the His_6_-*Hv*FdR concentration from 1.5 to 2.5 µM resulted in a modest increase in *V*_max_, the apparent *K*_m_ and *k*_cat_ values were similar within experimental error. The reaction additionally followed a classical hyperbolic profile consistent with Michaelis–Menten kinetics. Therefore, 1.5 µM His_6_-*Hv*FdR was selected for subsequent assays as it provided robust catalytic activity while minimizing substrate consumption.

**Figure 5.**
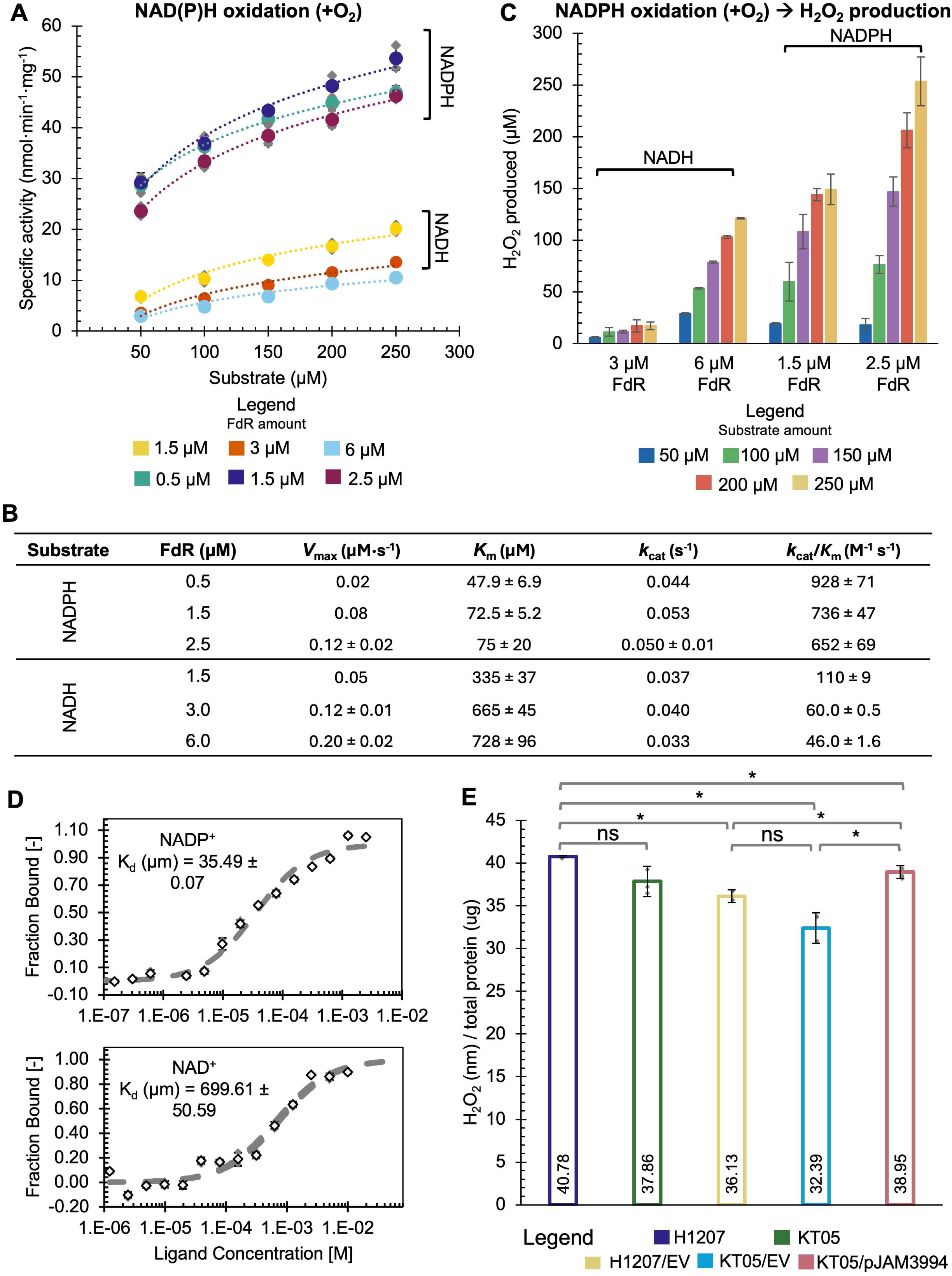
NADPH is the preferred electron donor of His_6_-*Hv*FdR under saturating oxidase activity conditions. Kinetic analysis of varying concentrations of FdR and NA(D)H under aerobic conditions in 150 µL total reaction. Kinetic analysis with NADPH was conducted in a reaction buffer containing 20 mM HEPES [pH 8] and 4 M NaCl at 51 °C, while analysis with NADH was conducted in a reaction buffer containing 20 mM MES [pH 6] and 3 M NaCl at 51 °C. Measurements were taken at A_340_ over 30 minutes. Specific activity (nmols·min^-1^·mg^-1^) is defined as nanomoles of NAD(P)H oxidized per minute per milligram of *Hv*FdR. A. Specific activity of varying concentrations of His_6_-*Hv*FdR and NAD(P)H. The circles represent the average of the three replicates and solid grey diamonds represent the individual replicates. For NADH, 1.5 µM (yellow), 3 µM (orange), and 6 µM (light blue) of His_6_-*Hv*FdR were added. For NADPH, 0.5 µM (teal), 1.5 µM (purple), and 2.5 µM (burgundy) of His_6_-*Hv*FdR were added. Dashed lines represent the best-fit curves generated from the mean specific activity values. B. Steady-state kinetic parameters of varying concentrations of His_6_-*Hv*FdR with NADPH and NADH as the electron donor (substrate) under aerobic conditions. *V*_max_ represents the maximal reaction velocity (µM·s^-1^). *K*_m_ denotes the Michaelis constant (µM), *k*_cat_ (s^-1^) is the catalytic turnover number. *k*_cat_/*K*_m_ (M^-1^ s^-1^) represents catalytic efficiency. Values are reported as mean ± standard deviation. Steady-state kinetic parameters were determined from the initial linear portion of each reaction, where less than 20% of the NAD(P)H substrate was consumed. C. H_2_O_2_ production of varying concentrations of His_6_-*Hv*FdR and NAD(P)H. ROS production was determined via ABTS oxidation (see Methods for details) after 30 minutes of NADPH oxidation. ABTS oxidation was monitored by measuring A_412_ at 51 °C. The bar graph represents the average of the three replicates and the grey diamonds represent the individual replicates. H_2_O_2_ production of 1.5 µM His_6_-*Hv*FdR and NADH, as well as 0.5 µM His_6_-*Hv*FdR and NADPH was undetectable by the assay. NAD(P)H was added at the following concentrations: 50 µM (blue), 100 µM (green), 150 µM (purple), 200 µM (orange), and 250 µM (yellow). D. Microscale thermophoresis (MST) of His_6_-*Hv*FdR as the target and NAD(P)^+^ as the ligand. Analysis was done at 24 °C and at 40% excitation power in buffer containing 20 mM HEPES [pH 7.5] and 2 M NaCl. A starting concentration of 0.5 µM His_6_-*Hv*FdR labeled with 50 nM RED-tris-NTA dye was used for a final concentration of 0.25 µM His_6_-*Hv*FdR and 25 nM Red-Tris NTA dye. According to the manufacturer instructions, fraction bound normalization is best suited for a direct comparison of binding affinities. *K*_d_, dissociation constants. Final concentration of 0.125 µM His_6_-*Hv*FdR and varying amounts of NADP^+^ (**top**) and varying amounts of NAD^+^ (**bottom**). E. H_2_O_2_ levels generated in cell lysate. Promega ROS-Glo H_2_O_2_ Assay was used for quantification (see Methods for details). Values represent normalized H_2_O_2_ levels measured from three biological and three technical replicates. *H. volcanii* H1207 (parent, dark purple), KT05 (Δ*fdr*, green), KT05/pJAM3944 (+His_6_-*Hv*FdR, pink), KT05/EV (empty vector, light blue), and H1207/EV (empty vector, yellow). A student’s *t*-test was used to determine the statistical significance (*p*-value <0.05, *; ns, non-significant) of H_2_O_2_ concentration (nM) in H1207 compared to KT05 (0.1), H1207/EV (0.007*), KT05/EV (0.01*), and KT05/pJAM3944 (0.04*). KT05 vs KT05/pJAM3944 (0.4). H1207/EV compared to KT05/EV (0.05) and KT05/pJAM3944 (0.01*). KT05/EV compared to KT05/pJAM3944 (0.01*).

### *Hv*FdR-mediated transfer of electrons from NAD(P)H to oxygen results in the production of hydrogen peroxide

To evaluate whether His_6_-*Hv*FdR can form reactive oxygen species during the oxidation of NAD(P)H, the production of hydrogen peroxide (H_2_O_2_) was monitored under aerobic conditions using a horseradish peroxidase (HRP)- coupled 2,2′-azino-bis(3-ethylbenzthiazoline-6-sulfonic acid) (ABTS) colorimetric assay. H_2_O_2_ production was not detectable at 1.5 µM His_6_-*Hv*FdR during NADH oxidation or 0.5 µM His_6_-*Hv*FdR for NADPH oxidation, as H_2_O_2_ levels were below the detection limit of the ABTS assay; therefore, these conditions were not included in analyses. However, when the enzyme concentration was increased to 3 µM and 1.5 µM, H_2_O_2_ production was detectable when either NADH or NADPH served as the electron donor, respectively (**Fig. 5C**). These findings establish the optimal enzyme concentration for activity and reveal *Hv*FdR can mediate H_2_O_2_ production utilizing NAD(P)H as the electron donor *in vitro*.

### NADPH is the preferred electron donor to *Hv*FdR

Microscale thermophoresis (MST) was next utilized to quantitatively assess the cofactor binding affinity of His_6_-*Hv*FdR. NAD(P)H-dependent oxidoreductases often exhibit catalytic activity with both NADPH and NADH. However, many enzymes display a distinct preference for one cofactor over the other, and in some cases exclusively interact with a single nicotinamide cofactor [8–10, 17, 27, 30, 60, 73]. As reduced cofactors can generate H_2_O_2_ under assay conditions, the oxidized forms (NADP^+^ and NAD^+^) were used in the MST experiment to evaluate binding interactions with His_6_-*Hv*FdR, independent of catalytic turnover. Compared to NADP^+^, His_6_-*Hv*FdR exhibited a 19.7-fold higher dissociation constant (*K*_d_) for NAD^+^, indicating a lower binding affinity for this cofactor (**Fig. 5D**). These results demonstrate *Hv*FdR has a preferential affinity for NADP(H), supporting its role as the physiological electron donor for this enzyme.

### Impact of *fdr* deletion on intracellular H_2_O_2_ production

*Hv*FdR shares structural homology with NAD(P)H oxidases, exhibits H_2_O_2_-producing activity *in vitro*, and contributes to NADP(H) redox homeostasis, as deletion of *fdr* increases the intracellular NADPH:NADP⁺ ratio. These observations suggested that loss of *Hv*FdR may alter intracellular H_2_O_2_ production. To examine this possibility, *H. volcanii* parent and *Δfdr* mutant strains were grown to log phase, quenched, lysed, and assayed for H_2_O_2_ levels (**Fig. 5E**). Although differences in intracellular H_2_O_2_ levels were observed among strains, these changes were not statistically significant due to variability among biological replicates. Nevertheless, the *Δfdr* mutant exhibited a trend toward reduced H_2_O_2_ levels compared with the parent strain, independent of the presence of the empty vector, suggesting that loss of *Hv*FdR may decrease intracellular peroxide production. Conversely, expression of His_6_-*Hv*FdR in the *Δfdr* mutant increased H_2_O_2_-levels relative to the empty vector control. Together, these findings suggest that *Hv*FdR contributes to intracellular H_2_O_2_ production, either directly through its NAD(P)H oxidase activity under oxidative conditions or indirectly by modulating pathways that influence peroxide metabolism.

### Amino acid substitutions that modulate catalytic efficiency and electron transfer properties

To further evaluate the biochemical properties of *Hv*FdR, amino acid substitutions were generated that were predicted to impact the flavin binding and/or electron transfer efficiency of this enzyme. Conserved residues positioned on opposite faces of the 3D-structural model of *Hv*FdR were selected including: K47 localized on the predicted oxygen/ferredoxin interaction surface and Y323 on the NADP(H)-binding interface. Alanine substitutions were used to disrupt FAD-binding interactions and potential electron donor/acceptor interactions, while aromatic substitutions were generated to assess whether preservation of the benzyl ring-maintained activity.

All single and double variants of His_6_-*Hv*FdR (K47A, Y323A, Y323W, K47A/Y323A and K47A/Y323W) were purified to homogeneity and found to migrate at 60 kDa by SDS-PAGE, similarly to the wild-type enzyme (WT) (**Fig. 6A**). UV–visible spectroscopy (**Fig. 6B**) was used to assess flavin incorporation by normalizing the A_460_ signal characteristic of the bound FAD to protein concentration (mg·mL^-1^). Fold changes in flavin occupancy were then calculated relative to WT, and enzyme concentrations for subsequent assays were adjusted accordingly to flavin occupancy. Relative to WT, the K47A variant exhibited 0.8-fold higher FAD occupancy, while the Y323A and K47A/Y323A variants displayed a 1.5-fold and 1.2-fold reduction in flavin content, respectively (**Supplemental Table S2**). While the single Y323W variant displayed minimal change in flavin incorporation, combining K47A with Y323W shifted the spectral properties to include a broad peak spanning wavelengths 425 to 475 nm. These results suggest that K47A and Y323A have opposite effects on flavin occupancy (increased vs. decreased), whereas Y323W combined with K47A produces an additive effect that alters the spectral properties of the enzyme-bound flavin. Notably, the A_280_ remained unchanged relative to the WT, indicating that the observed spectral shift was specific to the flavin chromophore and was not due to changes in the overall protein absorbance.

**Figure 6.**
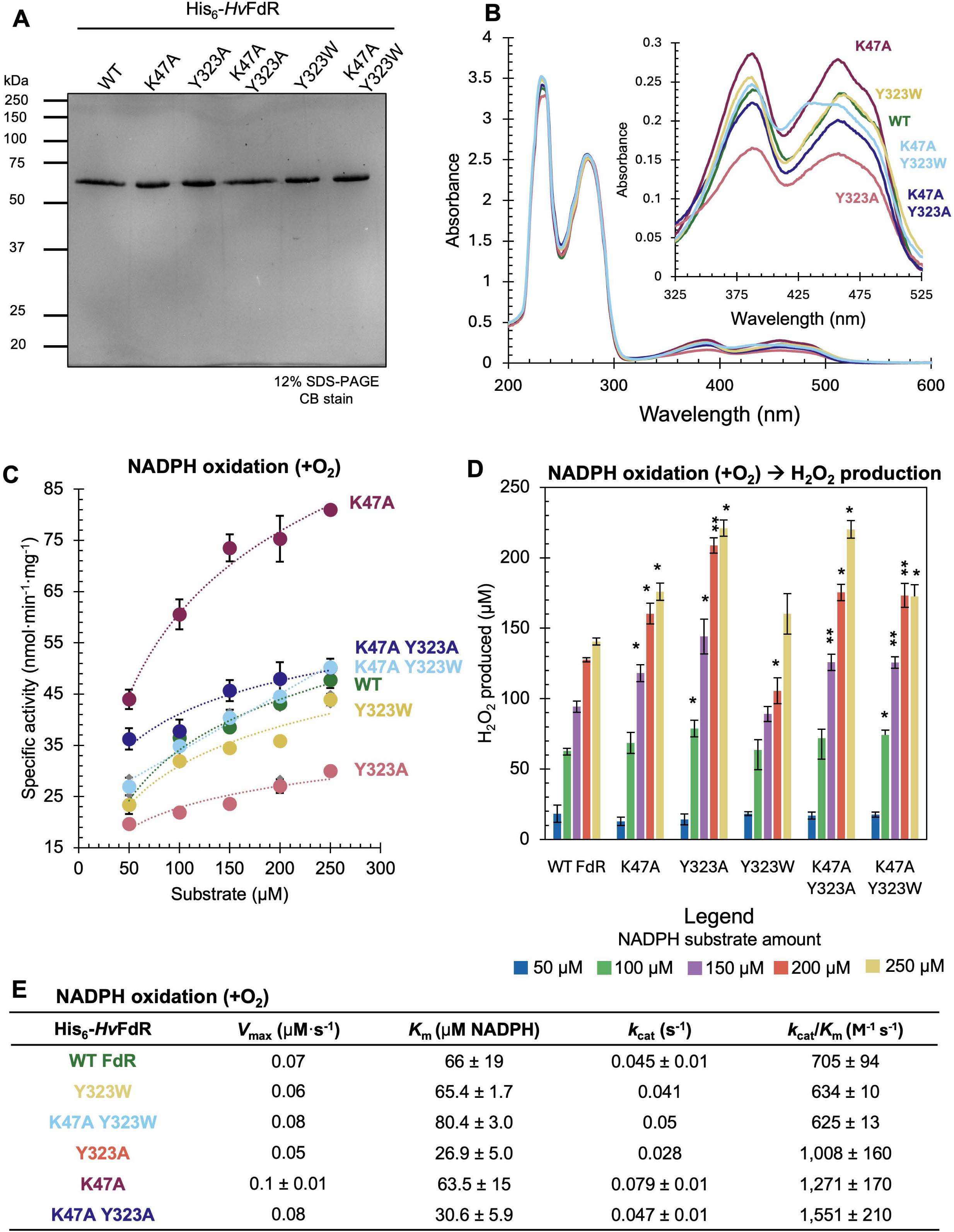
Purification and flavin binding of His_6_-*Hv*FdR proteins with amino acid substitutions. Proteins were purified by Ni-NTA affinity chromatography (HisTrap) and Superdex 75 10/300 GL chromatography from cells grown in ATCC974 rich medium supplemented with 0.2 µg·mL^-1^ novobiocin (see Methods for details). A. His_6_-*Hv*FdR wild type (WT) and variant proteins separated by reducing 12% SDS-PAGE and analyzed by Coomassie blue staining (CB stain). His_6_-*Hv*FdR samples were normalized to 1 µg for TCA precipitation and boiled for 10 min prior to SDS-PAGE. B. UV-visible spectrum (UVVIS), plotted by wavelength versus absorbance, of His_6_-*Hv*FdR and variant proteins (2 mg·mL^-1^) (K47A, Y323A, K47A Y323A, Y323W, and K47A Y323W). Protein absorbance was monitored from 200 to 600 nm aerobically. All His_6_-*Hv*FdR samples were found to exhibit two peaks at 389 nm and 460 nm, typical of other FAD-dependent proteins. Dark blue, wild type FdR; orange, K47A; dark green, Y323A; light blue, K47A Y323A; purple, Y323W; light green, K47A Y323W. C. Specific activity of His_6_-*Hv*FdR with and without amino acid substitutions. Varying amounts of protein (µM) were added to the reaction dependent on flavin occupancy (**Supplemental Table S2**). NADPH oxidation was tested using concentrations of 50-250 µM NADPH in a 150 µL reaction buffer containing 20 mM HEPES [pH 8] and 4 M NaCl at 51 °C. Measurements were monitored at A_340_ over 30 min. Specific activity is defined as nmol NAD(P)H oxidized per min per mg of protein. His_6_-*Hv*FdR WT (green), K47A (burgundy), Y323A (pink), Y323W (yellow), K47A Y323A (dark purple), and K47A Y323W (light blue). Dashed lines represent the best-fit curves generated from the mean specific activity values. D. H_2_O_2_ production was determined following the 30 min oxidation of NADPH. ROS production was determined via ABTS oxidation (see Methods for details). ABTS oxidation was monitored by measuring at A_412_ at 51 °C. NAD(P)H was added at the following concentrations: 50 µM (blue), 100 µM (green), 150 µM (purple), 200 µM (orange), and 250 µM (yellow). E. Steady-state kinetics of His_6_-*Hv*FdR and flavin binding variants utilizing NADPH as the electron donor under aerobic conditions. Kinetic parameters include *V*_max_ (µM·s^-1^), *K*_m_ (µM), *k*_cat_ (s^-1^), and *k*_cat_/*K*_m_ (M^-1^ s^-1^). Values are reported as mean ± standard deviation. Steady-state kinetic parameters were determined from the initial linear portion of each reaction, where less than 20% of the NADPH substrate was consumed.

The His_6_-*Hv*FdR variants were further analyzed for NADPH oxidation and H_2_O_2_ production. Specific activity measurements that monitored NADPH oxidation (**Fig. 6C**) showed that K47A exhibited 1.7-fold higher activity at 250 μM NADPH relative to WT. In contrast, Y323A and Y323W displayed a reduction in activity (1.8- and 1.1-fold, respectively), while both double variants (K47A/Y323A and K47A/Y323W) retained near WT activity. H_2_O_2_ production followed a similar trend across NADPH concentrations (**Fig. 6D**), although catalytic turnover and peroxide generation were not strictly proportional. Notably, Y323W produced the lowest H_2_O_2_ levels, whereas the other variants consistently generated statistically higher amounts of H_2_O_2_ compared to WT.

Further analysis of NADPH oxidation by steady-state kinetic analysis (**Fig. 6E**) demonstrated variant K47A/Y323A retained WT-like turnover but decreased affinity exhibiting the highest catalytic activity. Variants Y323W and K47A/Y323W displayed similar turnover and reduced catalytic activity compared to WT. Despite the single Y323A substitution reducing maximal turnover, this variant displayed improved efficiency due to a lower *K*_m_, supporting enhanced substrate affinity. Collectively, these findings suggest that Lys47 and Tyr323 modulate electron transfer efficiency by altering the local active-site environment, thereby influencing hydride transfer between NADPH and the enzyme-bound flavin.

### Thermal stability of *Hv*FdR and flavin binding variants

The effects of NADP^+^ binding and amino acid substitutions (K47A, Y323A, Y323W, K47A/Y323A and K47A/Y323W) on His_6_-*Hv*FdR thermal stability were assessed by differential scanning fluorimetry (DSF), using the intrinsic fluorescence of the flavin isoalloxazine ring [74]. NADP^+^ was used in place of NADPH to prevent H_2_O_2_ formation. All proteins displayed sigmoidal melting curves characteristic of cooperative thermal denaturation (**Fig. 7AB, left**), while first-derivative analysis revealed distinct transition profiles (**Fig. 7AB, right**). In the absence of NADP^+^, the WT, K47A and Y323A enzymes each displayed a single melting transition, with *T*_m_ values of 51.5 °C, 55.0 °C and 65.0 °C, respectively. In contrast, the remaining variants displayed two transitions suggesting altered FAD binding and/or distinct conformational states of protein subdomains. Addition of NADP^+^ did not affect the *T*_m_ of WT or K47A enzymes but increased thermal stability in all other variants and stimulated two transition states for Y323A (**Fig. 7AB, right**). The strongest stabilization by NADP^+^ was observed for the K47A/Y323A variant, in which the *T*_m_ at 52°C increased to 59.5 °C. Notably, all variants with substitutions at Y323 exhibited a second higher-temperature transition, a feature not observed in the WT or K47A alone. This additional transition may reflect distinct unfolding states or the presence of holo- and apo-enzyme conformations with different thermal stabilities. These results (summarized in **Fig. 7C**) suggest that the amino acid substitutions near the predicted flavin binding site altered the conformational properties and/or cofactor binding of His_6_- *Hv*FdR.

**Figure 7.**
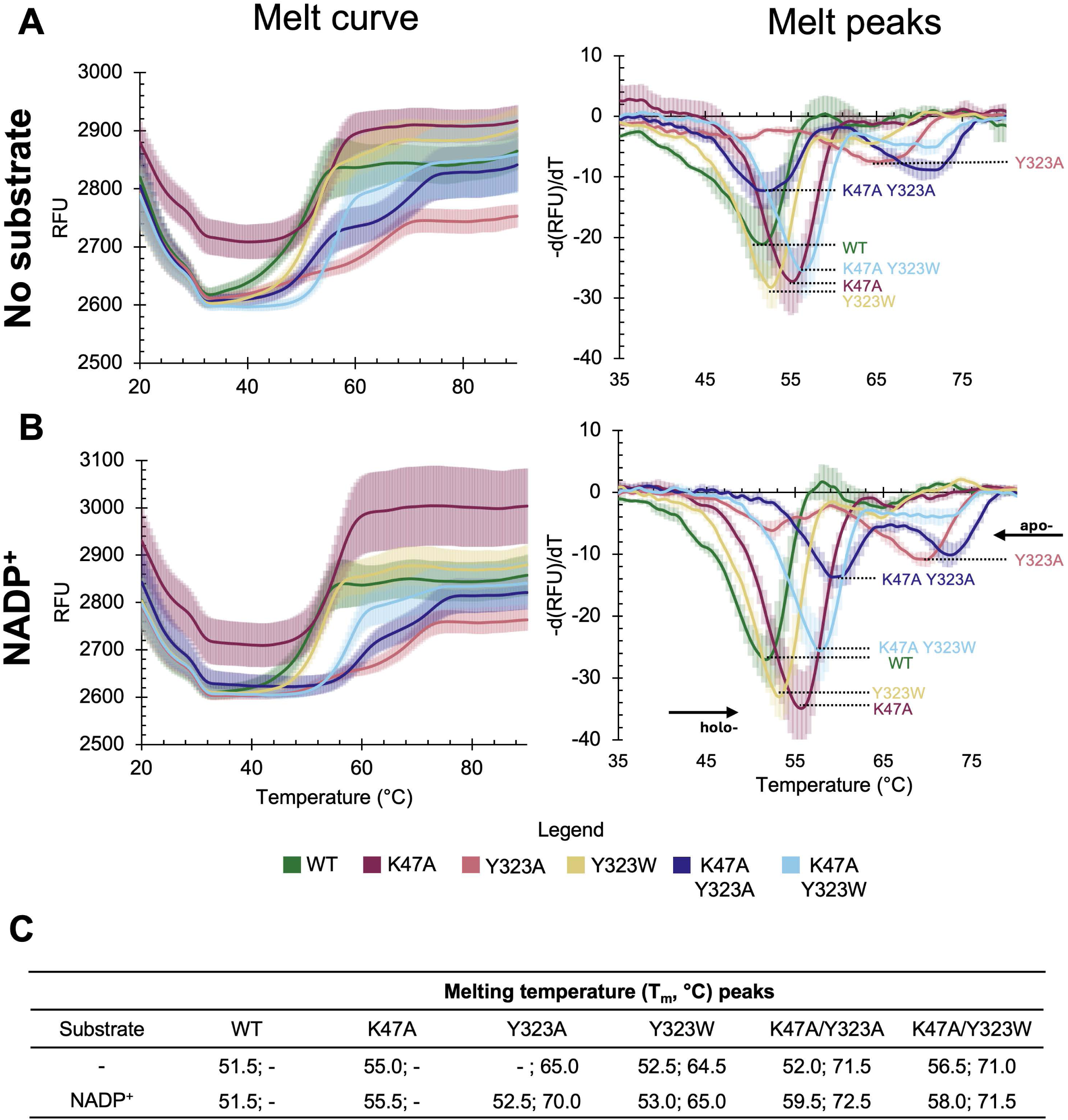
Thermal stability of *Hv*FdR in the presence of NADP^+^. Melting temperature (*T*_m_) peaks were determined for WT His_6_-*Hv*FdR, as well as single and double variants (K47A, Y323A, Y323W), in buffer containing 20 mM HEPES [pH 8] and 4 M NaCl via ThermoFAD methodology. WT (green), K47A (burgundy), Y323A (pink), Y323W (yellow), K47A Y323A (dark purple), and K47A Y323W (light blue). RFU, relative fluorescence units; d(RFU)/dT, rate at which fluorescence changes as temperature increases. A. Melt cure (left) and peaks (right) of enzyme alone. B. Melt curve (left) and peaks (right) of enzyme with 150 µM NADP⁺. C. Melting temperatures (°C) determined by analysis of the melt peaks. As indicated by -, His_6_-*Hv*FdR WT, K47A, and Y323A proteins displayed only a single melt peak under specified conditions. A student’s *t*-test was used to determine the statistical significance (*p*-value <0.05) of melt peak with WT FdR verse: K47A (4.95 × 10^-7^), Y323A (2.91 × 10^-4^), K47A/Y323A (0.02), Y323W (0.001), and K47A Y323W (1.84 × 10^-8^). For the second peak wild type FdR verses: K47A/Y323A (2.76 × 10^-12^), Y323W (9.88 × 10^-11^), and K47A/Y323W (7.36 × 10^-8^). A student’s *t*-test was used to determine the statistical significance (*p*-value <0.05) of melt peak with addition of 150 µM NADP^+^ and wild type FdR verse: K47A (2.61 × 10^-8^), Y323A (0.18), K47A/Y323A (3.50 × 10^-10^), Y323W (2.22 × 10^-4^), and K47A Y323W (1.33 × 10^-10^). For the second peak wild type FdR verses: Y323A (7.49 × 10^-14^), K47A/Y323A (4.29 × 10^-16^), Y323W (6.95 × 10^-14^), and K47A/Y323W (7.49 × 10^-10^).

### Redox potential of *Hv*FdR-bound FAD

The midpoint redox potential (*E*_0_′) of His_6_- *Hv*FdR-bound FAD was determined by anaerobic chemical reduction with sodium dithionite (Na_2_O_4_S_2_, *E*_0_′ = -660 mV, pH 7.0) as an electron donor/titrant and methyl viologen (MV^2+^) (MV^2+^; *E*_0_′ = -446 mV, pH 7.0) as an electron mediator/reference dye. After reduction equilibration, UV–visible spectra (**Fig. 8A**) were recorded to monitor the redox states of MV^•+^ at A_604_ and His_6_-*Hv*FdR-bound FAD at A_460_. Incremental addition of sodium dithionite decreased the A_460_ values, indicating the reduction of His_6_-*Hv*FdR- bound FAD. Fitting the experimental data to the rearranged Nernst equation (see method) yielded a midpoint redox potential of -413 ± 9 mV with 2.3 electrons transfer event (*n*) for His_6_-*Hv*FdR-bound FAD (**Fig. 8B**). Based on these results, *Hv*FdR-bound FAD exhibits a low midpoint redox potential (-413 mV) relative to the standard redox potential of the NADPH/NADP^+^ couple (-320 mV), indicating a strong thermodynamic role to function as an electron donor.

**Figure 8.**
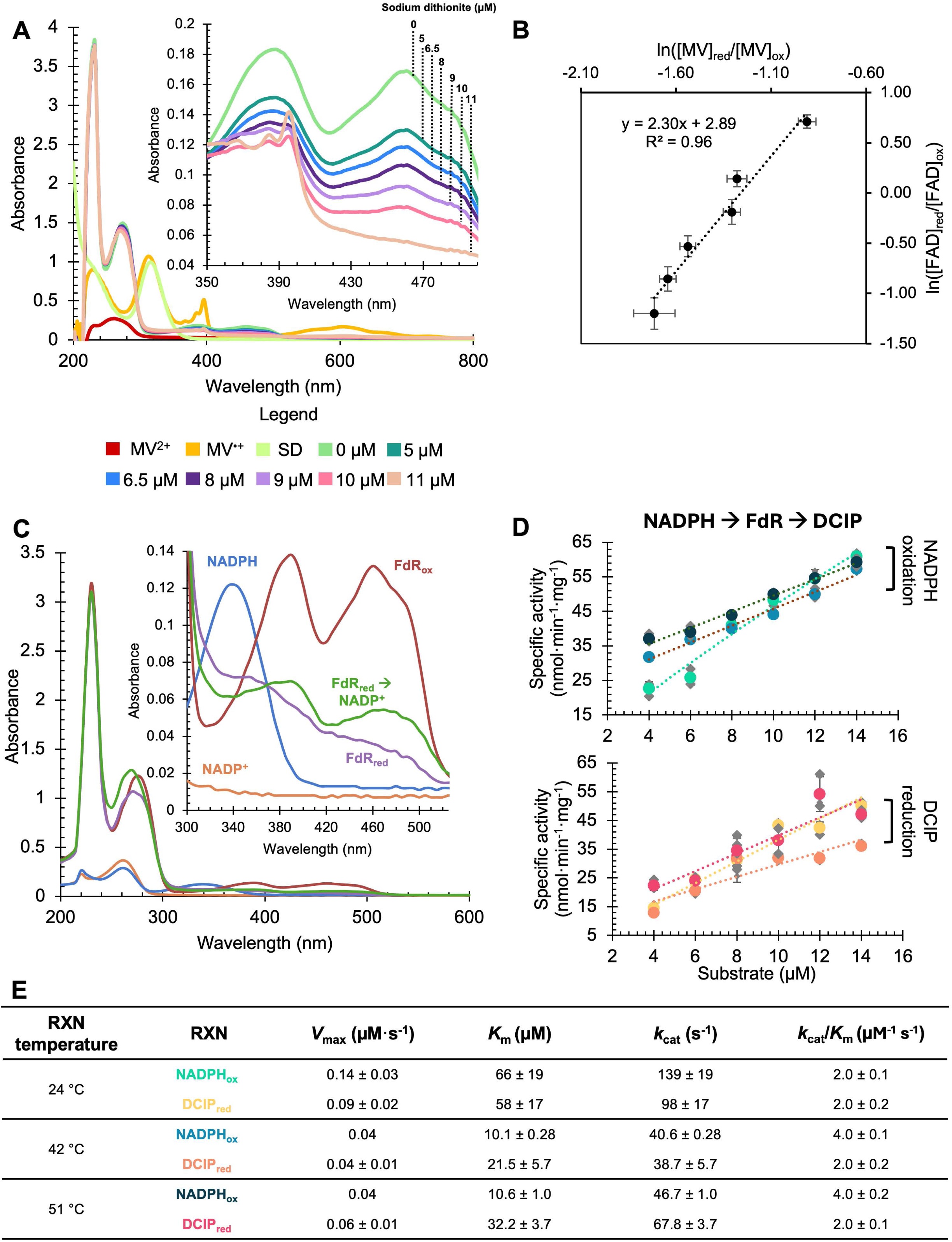
Determination of redox potential (E₀′) and electron transfer kinetics under anaerobic conditions. All reactions were performed under an N_2_:H_2_ (98:2) atmosphere in 20 mM HEPES [pH 7.5] containing 2 M NaCl at 25–27 °C, unless specified. All reagents were rendered oxygen-free prior to use using argon gas (100%). A. Oxidized FdR (20 µM) was reacted anaerobically with increasing concentrations of sodium dithionite (SD) in the presence of 10 µM methyl viologen (MV^2+^) as a redox mediator and reference dye. After 30 min of equilibration, UV–visible spectra were recorded. Values at A_604_ (MV^•^⁺) and A_460_ (FdR-bound FAD) were used to calculate the reduction state of the enzyme. B. The midpoint redox potential (E₀′) of FdR-bound FAD was determined by fitting the titration data to the modified Nernst equation (see method) by plotting ln([MV_red_]/[MV_ox_]) against ln([FAD_red_]/[FAD_ox_]). Control spectra were collected for 10 µM oxidized MV^2+^, 10 µM fully reduced MV^•+^ (generated with 100 µM SD), and 100 µM SD alone. Each data point represents the mean ± SD of three independent experiments. C. UV–visible spectrum of FdR reduced anaerobically with sodium dithionite and incubated with NADP^+^ (20 µM) in 20 mM HEPES [pH 7.5] and 2 M NaCl. Controls included NADP^+^, NADPH, and FdR under oxidizing and reducing conditions. Spectra (200-600 nm) were collected after 30 min to ensure equilibration. D. NADPH oxidation and DCIP reduction catalyzed by FdR. Reactions (100 µL) contained FdR_ox_ (1 nM) and the artificial electron acceptor DCIP (15 µM) in 20 mM HEPES, pH 7.5, and 2 M NaCl. Reactions were initiated by the addition of NADPH (0–14 µM). NADPH oxidation and DCIP reductions were monitored spectrophotometrically at A_340_ and A_600_, respectively. Measurements were recorded at 1 min intervals for 30 min at 28 °C, 42 °C, 51 °C. Specific activity was calculated in nanomoles of substrate oxidized or reduced per minute per mg of enzyme (nmol·min^-1^·mg^-1^). Dashed lines represent the best-fit curves generated from the mean specific activity values. E. Steady-state kinetics of NADPH oxidation (NADPH_ox_) and DCIP reduction (DCIP_red_) under anerobic conditions. Kinetic parameters include *V*_max_ (µM·s^-1^), *K*_m_ (µM), *k*_cat_ (s^-1^), and *k*_cat_/*K*_m_ (µM^-1^ s^-1^). Values are reported as mean ± standard deviation. Steady-state kinetic parameters were determined from the initial linear portion of each reaction, where less than 20% of the NADPH substrate was consumed.

### Anaerobic electron transfer and directionality of electron flow

His_6_-*Hv*FdR was next evaluated for its ability to transfer electrons to NADP⁺ under anaerobic conditions. *Hv*FdR_ox_ was first reduced with sodium dithionite (SD), dialyzed to remove excess sodium dithionite, then the reduced enzyme (*Hv*FdR_red_) was tested for its ability to transfer electrons to NADP⁺ at a 1:1 stoichiometric ratio based on an increase in A_460_, consistent with oxidation of the enzyme-bound flavin cofactor (**Fig. 8C**). An increase in A_460_ was observed, consistent with oxidation of His_6_-*Hv*FdR following incubation with NADP^+^ under anaerobic conditions. Additionally, the ability of *Hv*FdR_red_ to donate electrons to NADP⁺ following dialysis demonstrated that sodium dithionite treatment did not irreversibly denature the enzyme. In contrast, under anaerobic conditions, at equal molar ratios, NADPH did not reduce His_6_-*Hv*FdR (*Hv*FdR_ox_), as evidenced by the absence of a decrease in A_340_ or A_460_, even when present at 1-, 2-, or 3-fold molar excess relative to the enzyme.

Electron transfer was further evaluated under anaerobic conditions using saturating NADPH as the electron donor and dichlorophenolindophenol (DCIP, +217 mV redox potential) as an artificial electron acceptor. Under these conditions, His_6_- *Hv*FdR (*Hv*FdR_ox_) catalyzed rapid NADPH oxidation and efficiently transferred electrons to DCIP, exhibiting diaphorase activity (**Fig. 8D**). Steady-state kinetic analysis revealed high catalytic turnover for both NADPH oxidation and DCIP reduction, with catalytic efficiencies about 10^3^ times higher than under aerobic conditions (**Fig. 8E vs. Fig. 5B**). Notably, the reaction proceeded rapidly, reaching completion within minutes even at nanomolar enzyme concentrations, demonstrating high catalytic activity while confirming that His_6_-*Hv*FdR remains enzymatically active under the conditions tested.

### *Hv*FdR displays ferredoxin reductase activity

The efficient anaerobic electron transfer properties of His_6_-*Hv*FdR suggested that it functions as a ferredoxin reductase. To evaluate this possibility, 3D structural modeling and biochemical assays were performed. AlphaFold modeling predicted that the [2Fe-2S] ferredoxin FerA5 of *H. volcanii* (*Hv*Fdx; HVO_2995) forms a complex with *Hv*FdR (**Fig. 9A**). The 3D-model revealed close spatial organization between the NADPH-binding pocket and FAD cofactor of *Hv*FdR and the [2Fe–2S] cluster of *Hv*Fdx, with predicted electron transfer distances compatible with productive redox coupling (**Fig. 9A**). Biochemical analyses demonstrated that His_6_-*Hv*FdR efficiently transferred electrons from NADPH through its FAD cofactor to *Hv*Fdx, followed by reduction of DCIP as the terminal electron acceptor. In the His_6_-*Hv*FdR activity assay, NADPH oxidation and DCIP reduction activity were elevated by the addition of *Hv*Fdx in a concentration-dependent manner (**Fig. 9B**). Addition of 3.5 μM *Hv*Fdx resulted in 2.5- and 3.3-fold higher for NADPH oxidation and DCIP reduction activities, respectively (**Fig. 9B**), consistent with efficient electron transfer through His_6_-*Hv*FdR to *Hv*Fdx. Kinetic analysis showed that electron transfer through the NADPH → *Hv*FdR → *Hv*Fdx → DCIP (**Fig. 9B**) pathway was more efficient than direct NADPH → *Hv*FdR → DCIP reduction alone (**Fig. 8D**), supporting a preferential interaction between *Hv*FdR and its ferredoxin partner (**Fig. 9C**). Additionally, NADPH was unable to transfer electrons to *Hv*Fdx in the absence of *Hv*FdR, indicating that *Hv*FdR is required to mediate electron transfer from NADPH. Collectively, these findings identify *Hv*FdR as a physiological ferredoxin reductase *in vitro* capable of coupling NADPH oxidation to ferredoxin reduction under anaerobic conditions.

**Figure 9.**
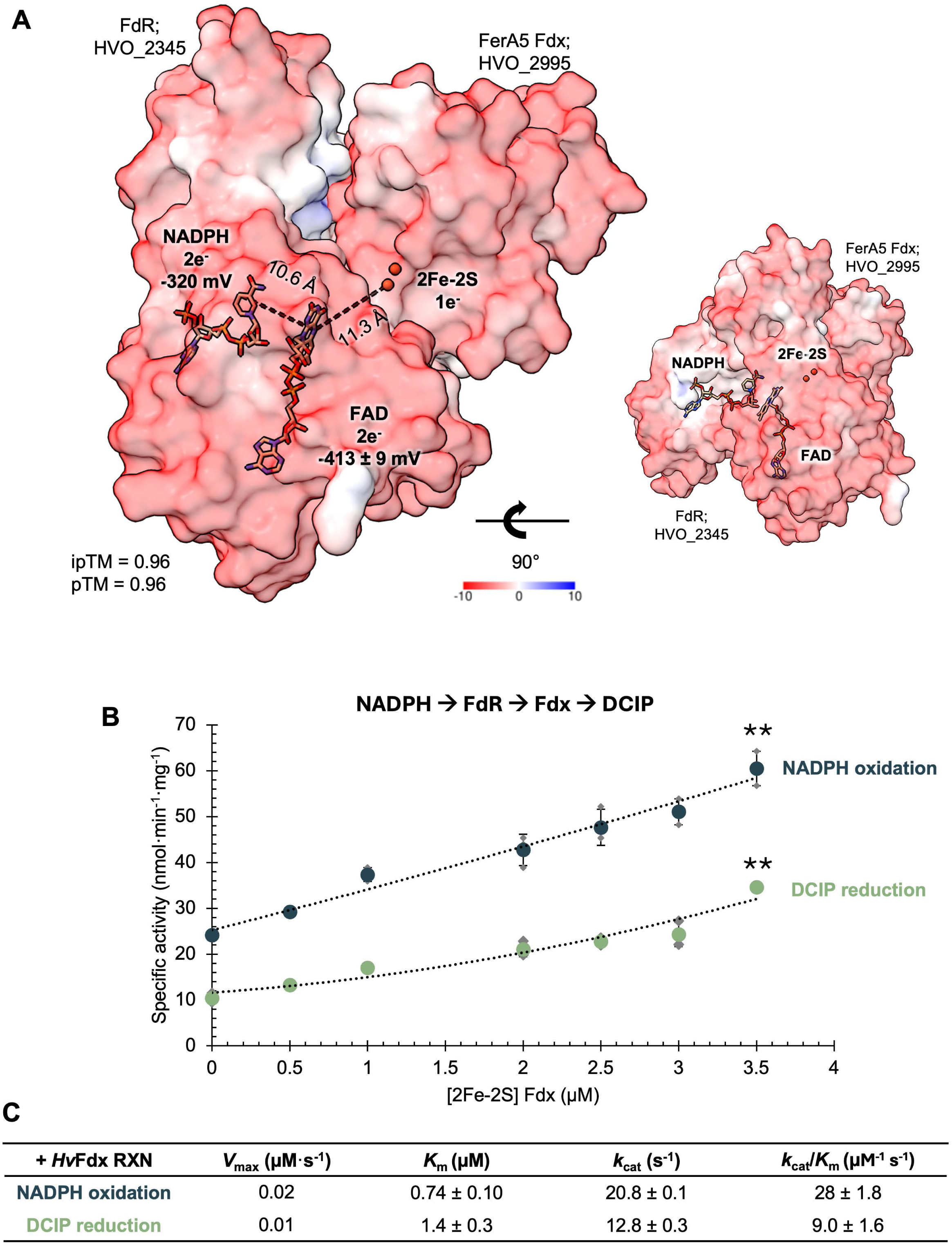
Structural modeling and biochemical characterization of electron transfer between *Hv*FdR and *Hv*Fdx. A. The schematic depicts the proposed ferredoxin reductase two-electron transfer mechanism, where NADPH donates a hydride equivalent (2e^⁻^) to FAD, transferring electrons to downstream to [2Fe-2S] ferredoxin (FerA5, HVO_2995). AlphaFold 3 server model (ipTM = 0.96 and pTM = 0.96) of *Hv*FdR, FAD, NADPH, 2Fe^3+^, and *Hv*Fdx. Electrostatic surface representation of the ferredoxin reductase colored by surface potential (red, negative; blue, positive; scale shown). The distance between C4 carbon of NADPH nicotinamide ring to the N5 nitrogen of the FAD is approximately 10.6 Å, while the N5 FAD to [2Fe-2S] is approximately 11.3 Å. Midpoint redox potentials: NADPH, -320 mV; FdR, -413 ± 9 mV (experimentally characterized). A 90° rotated view (right) highlights the spatial alignment of the cofactors within the protein matrix. B. NADPH oxidation and DCIP reduction catalyzed by FdR-Fdx. Reactions (100 µL) contained FdR_ox_ (1 nM), Fdx_ox_ (0-3.5 µM), and the artificial electron acceptor DCIP (15 µM) in 20 mM HEPES, pH 7.5, and 2 M NaCl. Reactions were initiated by the addition of NADPH (4 µM). NADPH oxidation (dark blue) and DCIP reduction (light green) were monitored spectrophotometrically at A_340_ and A_600_, respectively. Measurements were recorded at 1 min intervals for 30 min at 28 °C. Specific activity was calculated in nanomoles of substrate oxidized or reduced per minute per mg of enzyme (nmol·min^-1^·mg^-1^). The addition of 3.5 μM *Hv*Fdx significantly increased both NADPH oxidation and DCIP reduction relative to reactions lacking *Hv*Fdx (NADPH, *p* = 0.003; DCIP, *p* = 0.0006). Dashed lines represent the best-fit curves generated from the mean specific activity values. C. Steady-state kinetics of NADPH oxidation and DCIP reduction under anerobic conditions. Kinetic parameters include *V*_max_ (µM·s^-1^), *K*_m_ (µM), *k*_cat_ (s^-1^), and *k*_cat_/*K*_m_ (µM^-1^ s^-1^). Values are reported as mean ± standard deviation. Steady- state kinetic parameters were determined from the initial linear portion of each reaction, where less than 20% of the NADPH substrate was consumed or DCIP was reduced.

## Discussion

Structural and biochemical characterization demonstrated that *Hv*FdR is a versatile redox-active flavoprotein. *Hv*FdR purified as a soluble, monomeric protein containing non-covalently bound FAD, a mode of flavin association characteristic of most flavoproteins, in which binding is mediated through hydrogen bonding and electrostatic interactions rather than covalent attachment of the flavin [6, 75]. The partial flavin occupancy observed for purified *Hv*FdR is also consistent with other FAD-dependent oxidoreductases [76–78]. Collectively, these findings indicate that *Hv*FdR conforms to canonical flavoprotein architecture, with non-covalent FAD binding likely supporting conformational flexibility and dynamic redox cycling.

Through chromosomal mutational analysis, insight is provided regarding the physiological role of *Hv*FdR. Deletion of *fdr* impaired cellular growth and altered intracellular NADP(H) balance, supporting a physiological role for *Hv*FdR in maintaining redox homeostasis. The accumulation of NADPH in the Δ*fdr* strain suggests that *Hv*FdR contributes to cellular cofactor balance. As all strains were harvested at the same growth stage, these differences are unlikely to reflect growth-dependent variation. In both cases, expression of His_6_-*Hv*FdR in the Δ*fdr* strain restored growth and NADPH redox balance, indicating that the tag does not impair protein function. The slight growth difference relative to the wild type strain is likely due to the metabolic burden associated with plasmid-based expression rather than impaired *Hv*FdR activity. A similar trend was observed when the genes encoding the multisubunit flavin-based NAD(P)H-dependent ferredoxin NADP^+^ oxidoreductases (NfnI PF1327-1328 and NfnII PF1910-1911) were deleted in *Pyrococcus furiosus*, resulting in impacted growth phenotypes and altered intracellular concentrations of NAD(P)H [79]. Altogether, *Hv*FdR directly or indirectly through interactions with other redox proteins, supports optimal cellular growth and contributes to maintaining intracellular NADP(H) redox homeostasis.

With its potential to act as an oxidase, NAD(P)H oxidase activity was utilized to probe for *Hv*FdR-dependent NAD(P)H oxidation. *Hv*FdR activity was strongly influenced by pH, salt concentration, and temperature. Preference for NADPH oxidation under alkaline conditions may reflect improved stabilization and recognition of the 2′- phosphate group by positively charged residues, such as arginine, whose interactions may be favored at higher pH values [80, 81], whereas NADH utilization under acidic conditions may rely more heavily on hydrogen bonding and electrostatic interactions [82]. Extended incubation of *Hv*FdR at temperatures above the organism’s optimal growth range (≥51 °C) and at low salt concentrations (≤0.05 M) destabilized the enzyme, as observed for other haloarchaeal proteins [83, 84]. Addition of exogenous FAD or FMN to purified *Hv*FdR preparations did not result in detectable NADPH oxidation or improve thermostability of the wild-type enzyme under the conditions tested. High concentrations of free flavin likely interfere with electron transfer by competing for interactions near the active site or by acting as an alternative redox sink, thereby decreasing the observed rate of NADPH oxidation. These experimental results indicate that free flavin cofactors in solution were insufficient to increase *Hv*FdR activity, supporting the importance of the enzyme-bound flavin for function.

With its ability to oxidize NAD(P)H under aerobic conditions, *Hv*FdR was found to generate H_2_O_2_ at elevated enzyme and cofactor concentrations. **Table 2** summarizes our findings regarding the NAD(P)H substrate specificity and oligomeric state of *Hv*FdR in relation to other FAD-dependent oxidoreductases from bacteria and archaea, which display considerable diversity in these biochemical properties. Compared to the *S. solfataricus* NAD(P)H oxidase SsNOX38 (SSO2222, which shares only 22.5% amino acid identity), reduced O_2_ in a stoichiometric manner relative to NADH [62]; however, this did not occur for *Hv*FdR except at enzyme concentrations of 2.5 µM with NADPH. Thus, we suggest the *Hv*FdR NADPH oxidase activity is physiologically relevant under conditions of high intracellular NADPH and oxygen availability, where the reduction of oxygen represents a secondary or condition-dependent activity rather than the primary physiological role of the enzyme. *H. volcanii* encodes numerous oxidative stress defense systems, including thioredoxin-dependent peroxiredoxins (HVO_2699), superoxide dismutase (HVO_2913), peroxidases (HVO_1778), as well as various thioredoxins and rubredoxins [85, 86] indicating that *Hv*FdR likely functions within a broader redox network involved in oxidative stress adaptation.

**Table 2.** Comparison of NAD(P)H oxidase activities across extremophilic microorganisms.

| Organism | Enzyme | UniProt | Pfam<sup>a</sup> | Oligomeric State | Cofactor | Electron donors | $K_m$ (NADH) μM | $K_m$ (NADPH) μM | Ref |
| --- | --- | --- | --- | --- | --- | --- | --- | --- | --- |
| *Thermus thermophilus* HB8 | TTHA0425 | Q60049 | PF00881 | Monomer | FAD | NADH, NADPH | 4.14 | 14.0 | [59] |
| *Thermotoga maritima* | TM_1433 | Q9X1E7 | PF07992 | Heterodimer | FAD | NADH | 42.0 ± 3 | - | [71] |
| *Sulfolobus acidocaldarius* | Saci_2144 | Q4J6Z4 | PF07992 | Monomer | - | - | 21 | - | [60, 61] |
| *Acidianus ambivalens* | NoxA | Q7ZAG8 | PF07992 | Monomer | FAD | NADH | - | - | [27] |
| *Archaeoglobus fulgidus* | NoxA2 | O29852 | PF07992; PF02852 | Monomer | FAD | NADH | 3.1 | - | [58, 118] |
| *Haloferax volcanii* | HvFdR | D4GWJ4 | PF07992; PF18267 | Monomer | FAD | NADH, NADPH | 335 | 72 | This study |
| *Pyrococcus furiosus* | Nox1 | Q8U1K9 | PF07992; PF22353 | Dimer | FAD | NADH | < 4.0 | - | [10] |
| *Sulfolobus solfataricus* | SsNOX38 | Q97WJ5 | PF07992 | Homodimer | FAD | NADH, NADPH | 413 | - | [60, 61] |
| *Thermococcus kodakarensis* | TK1481 | Q5JJB9 | PF07992; PF02852 | Homotetramer | FAD | NADH, NADPH | 47.1 | 42.9 | [30] |

<sup>a</sup>PF07792 and PF02852: class I and class II oxidoreductases, NADH oxidases, peroxidases. PF1862 and PF22353: NAD(P)H:rubredoxin oxidoreductase.

Although many flavin-dependent reductases have been structurally characterized, the mechanistic roles of conserved flavin-site residues in modulating stabilization and electron transfer remain incompletely resolved across diverse systems. In *Hv*FdR, K47 and Y323 are positioned near the FAD-binding pocket, where the K47A substitution increased catalytic activity while increasing flavin binding, suggesting that K47 is not critical for cofactor stabilization. Reduced flavin incorporation observed in the Y323A variant suggests that aromatic interactions near the isoalloxazine ring contribute to FAD stabilization within the binding pocket. Similar residues modulating catalytic efficiency and electron transfer in related systems have been reported. For example, in *Rhodopseudomonas palustris*, Y328 variants in a homodimer type ferredoxin-NADP^+^ oxidoreductase decreased *K*_m_ and increased catalytic efficiency, likely by altering nicotinamide access to the flavin [87], whereas substitutions of K328 in the putidaredoxin reductase weakened binding of the ferredoxin (putidaredoxin) and supported the FAD si-face as the ferredoxin interaction interface [17]. Consistent with these studies, the altered catalytic efficiencies and increased H_2_O_2_ production observed in *Hv*FdR alanine-substitution variants suggest that K47 and Y323 normally impose structural or electrostatic constraints that help regulate electron transfer and limit uncontrolled reduction of oxygen. Such restrained catalytic behavior may be advantageous for facultative anaerobes such as *H. volcanii*, which encounter fluctuating oxygen concentrations and reactive oxygen species during growth [45, 88–91].

Thermal stability analysis further supported the role of flavin occupancy in stabilizing *Hv*FdR structure. Biphasic melting transitions observed in Y323 variants likely reflect heterogeneous populations containing both flavin-bound (larger peak) and flavin-free (smaller peak) protein species or differential un-folding states, consistent with the stabilizing effect of flavin cofactors reported in other flavoproteins [92, 93]. The absence of biphasic transitions in the WT and K47A variants suggests that substitution of Y323 alters the unfolding or conformational landscape of the enzyme. Previous studies have shown that while some flavoproteins exhibit changes in thermal stability between apo- and holo- forms [94, 95], others do not [5], suggesting that the observed decrease in melting temperature may reflect domain-specific unfolding or weakened FAD binding. Interestingly, WT and K47A exhibited the highest flavin occupancy but smaller fluorescence signal (-d(RFU)/dT), indicating that peak magnitude did not correlate directly with flavin content. In addition, thermal shift analysis indicated conformational changes occurring around 51°C, suggesting partial opening or destabilization of the flavin-binding interface of *Hv*FdR. We suggest as the predicted flavin-gating residues of wild type *Hv*FdR become more flexible at elevated temperature, specific activity initially increases, potentially due to enhanced electron transfer accessibility. However, prolonged incubation at elevated temperatures (1 h, ≥51 °C) decreased activity, consistent with progressive thermal denaturation or degradation of the enzyme, resulting in loss of its catalytic organization.

*Hv*FdR was found to have a relatively low midpoint potential, which may favor electron donation from the reduced flavin of *Hv*FdR to NADP^+^ (-320 mV) under anaerobic conditions [96]. The high ionic strength of the haloarchaeal cytosol may interfere with electrostatic interactions required for efficient electron transfer, whereas the relatively negative midpoint potential of *Hv*FdR could offset this effect and strengthen the thermodynamic driving force for redox cycling. *Hv*FdR efficiently mediates electron transfer from saturating concentrations of NADPH to artificial acceptors, such as DCIP, under anaerobic conditions, demonstrating diaphorase activity at a higher catalytic efficiency than under oxidative conditions (*V*_max_, 2-fold and catalytic efficiency, 10^3^-fold). At saturating NADPH concentrations, the reaction equilibrium may shift to favor electron flow toward *Hv*FdR reduction. The ability of *Hv*FdR to mediate reverse electron transfer and reduce electron acceptors, such as ferredoxin, further supports a bi-functional and -directional role in electron transfer.

Previous studies showed that a ferredoxin-NADP^+^ reductase from spinach exhibits NADPH oxidase and diaphorase activities but cannot reduce cytochrome *c* [97]. Similarly, we found *Hv*FdR unable to transfer electrons to spinach cytochrome *c*, likely influenced by the high-salt required for optimal enzymatic assay and negatively charged protein surface of *Hv*FdR. Some NADH oxidases also function as broader oxidoreductases, transferring electrons to alternative acceptors [59]. For example, the *E. coli* Nox reduces quinones [98], while NADH oxidases from *Desulfurolobus ambivalens* and *A. acidianus* function as NADH:ferredoxin oxidoreductases under anaerobic conditions [27, 28]. These enzymes have additionally been linked to sulfate respiration, H_2_O_2_ production, and host interaction processes [99–101].

Collectively, these findings support a model in which *Hv*FdR functions as a flexible redox hub that modulates electron transfer according to environmental and metabolic conditions (**Fig. 10**). In this model, K47 located at the electron acceptor interface, likely influences accept accessibility and flavin intermediate stabilization, whereas Y323, positioned near the NADPH-binding site, modulates hydride transfer to FAD. These results suggest that K47 and Y323 may act as evolutionary gatekeeping residues that restrict uncontrolled electron flow and limit O_2_ reduction. By tuning flavin accessibility and cofactor positioning, these residues likely favor directed anaerobic electron transfer to physiological partner proteins while minimizing electron leakage and ROS formation. All together, these structural features appear to regulate cofactor stabilization, electron transfer efficiency, and reactive oxygen species formation under oxidative conditions, thereby balancing catalytic activity with oxidative stress adaptation in *H. volcanii*.

**Figure 10.**
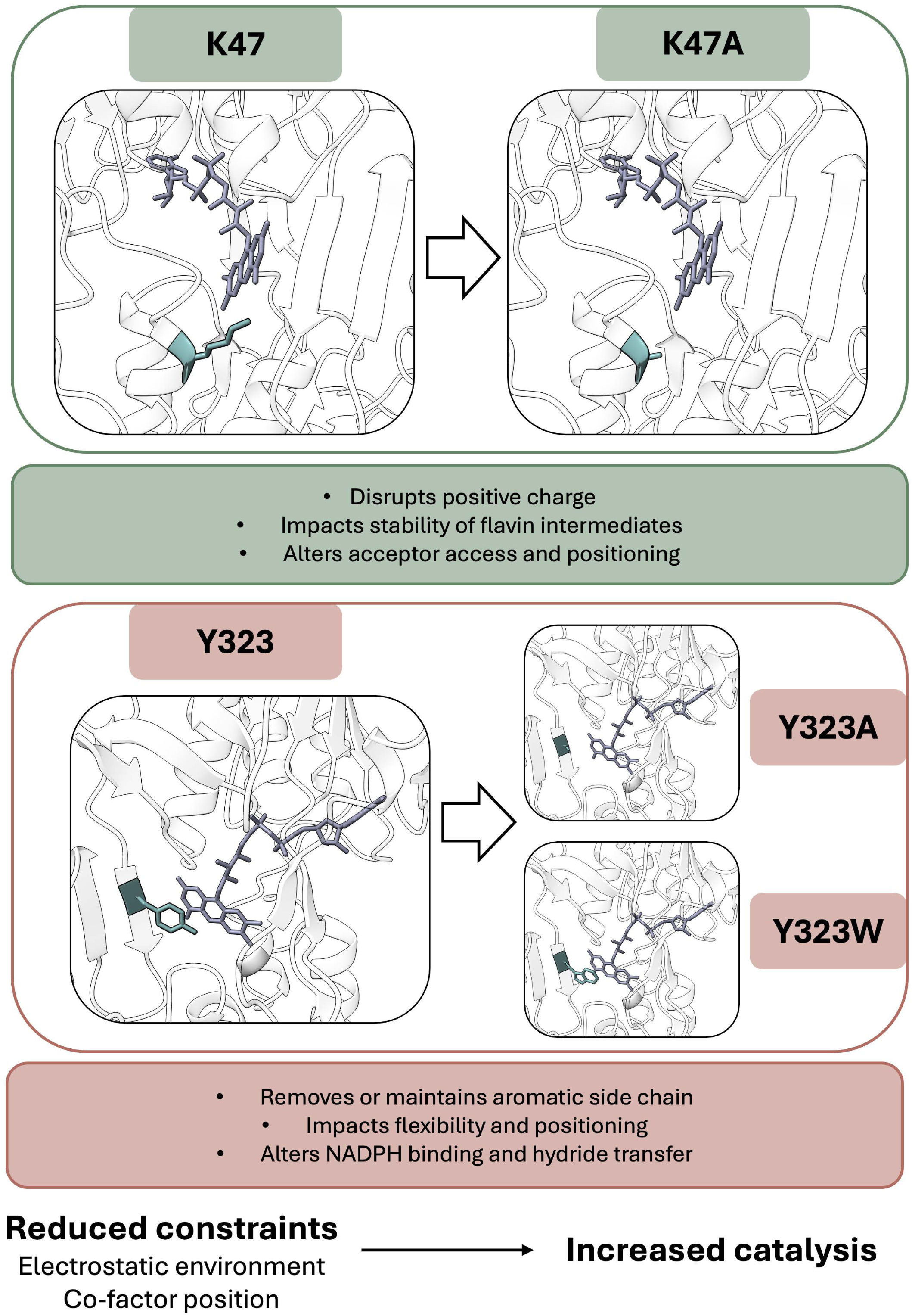
Structural modeling of *Hv*FdR active-site residue variants involved in electron transfer and cofactor interactions. Structural models support a role for Lys47 and Tyr323 in modulating flavin accessibility and stability, cofactor positioning, and electron transfer dynamics within *Hv*FdR. **Top:** The K47A substitution is located adjacent to the FAD cofactor, where replacement of Lys47 with alanine removes the positively charged ε-amino group and its hydrogen-bonding capability. This substitution is suggested to disrupt electrostatic interactions that contribute to stabilization of flavin redox intermediates. Removal of the lysine side chain may also increase local conformational flexibility and improve positioning of the electron acceptor for catalysis. **Bottom**: Tyr323 is positioned between the FAD cofactor and the NADPH-binding site, where it is proposed to contribute to substrate positioning and hydride transfer. Substitution of Tyr323 with alanine removes both the aromatic side chain and hydroxyl group, eliminating π-stacking interactions and hydrogen-bonding potential that may stabilize and orient the nicotinamide ring for efficient hydride transfer. This substitution is suggested to increase local flexibility and perturb the geometry of the electron transfer pathway. In contrast, substitution with tryptophan retains an aromatic residue but introduces a larger indole side chain lacking the tyrosine hydroxyl group. The increased steric bulk and altered geometry may modify interactions with NADPH and the flavin cofactor, potentially affecting nicotinamide positioning and hydride transfer efficiency while preserving aromatic character.

## Materials and Methods

### Materials

Biochemicals were purchased from Fisher Scientific (Atlanta, GA, USA), Bio-Rad (Hercules, CA, USA), and Sigma-Aldrich (St. Louis, MO, USA). Phusion High-Fidelity DNA Polymerase (cat. no. M0530), restriction enzymes (NdeI cat. no. R0111S, BlpI cat. no. R0585S, BamHI-HF cat. no. R3136S, HindIII-HF cat. no. R3104S, and DpnI cat. no. R0176), Antarctic phosphatase (cat. no. M0289S), 2x Quick Ligase (cat. no. M2200), and KLD enzyme mix (cat. no. M0554S) were purchased from New England Biolabs (NEB, Ipswich, MA, USA). GeneRuler 1 kb plus DNA ladder (cat. no. SM1331) was purchased from Thermo Fisher Scientific (Waltham, MA, USA). Eurofins Genomics (Louisville, KY, USA) was used for oligonucleotide synthesis and DNA Sanger sequencing services.

### Strains and media

Strains, plasmids, and primers used in this study are listed in **Supplemental Tables S3-4**. *Escherichia coli* strains were grown at 37 °C in Luria-Bertani (LB) medium that was supplemented with ampicillin (100 μg·mL^-1^) for plasmid selection. *H. volcanii* strains were grown at 42 °C in ATCC 974 medium composed of 125 g NaCl, 50 g MgCl_2_·6H_2_O, 5 g K_2_SO_4_, 0.2 g CaCl_2_·6H_2_O, 5 g tryptone and 5 g yeast extract per liter, adjusted to pH 6.8 using 1 N KOH. ATCC 974 medium was supplemented with novobiocin (Nov, 0.2 μg·mL^-1^) for plasmid selection or 5-fluoroorotic acid (5-FOA, 50 μg·mL^-1^ dissolved in DMSO) for chromosomal deletion. *H. volcanii* strains were also grown in Hv-Ca⁺ medium supplemented with uracil (50 μg·mL^-1^) according to the *Halohandbook* [102]. For solid plates, 1.5% (w/v) agar was included. Strains were stored at -80 °C in 20% (v/v) glycerol stocks. The *H. volcanii* strains were prepared for the -80 °C storage by diluting stationary phase cultures at a 1:4 ratio with a cryopreservation solution consisting of 80 mL 100% glycerol, 20 mL of 30% salt water, and 0.2 mL of 0.5 M CaCl_2_.

### PCR and plasmid isolation

Genomic DNA (gDNA), extracted from *H. volcanii* H26 by DNA spooling method [102], was used as the template for PCR-based amplification of target genes. PCR was performed in GC buffer using the high-fidelity Phusion DNA polymerase supplemented with 3% (v/v) DMSO, following manufacturer’s instructions (NEB). To initiate PCR by ‘hot start’, the reaction mixtures were transferred directly from ice to the denaturation temperature, and thermal cycling was performed using a T100 ThermoCycler (Bio-Rad Laboratories, Hercules, CA, USA). PCR products and other DNA fragments were separated by 0.8% (w/v) agarose gel electrophoresis (90 V and 30 min). DNA fragments were isolated using Monarch PCR & DNA Cleanup Kit (cat. no. T1030) and DNA Gel Extraction Kit (cat. no. T1020) (NEB, Ipswich, MA, USA). During plasmid construction, plasmids were transformed into *E. coli* Top10, with transformants selected on LB medium supplemented with ampicillin (100 μg·mL^-1^). Plasmid DNA was isolated using the Invitrogen PureLink Quick Plasmid Miniprep Kit (cat. no. K210011) (Waltham, MA, USA). Primers and restriction enzymes utilized were validated *in silico* using NEB Cloner and SnapGene software (v8.0), to ensure proper orientation and to check for potential frameshift mutations. The fidelity of all cloned plasmids was confirmed via Sanger DNA sequencing (Eurofins Genomics, Louisville, KY, USA).

### Generation of the *H. volcanii Δfdr* mutant strain

*H. volcanii* mutant strain KT05 (H1207 *Δfdr*, where *fdr* corresponds to *hvo_2345*) was generated using a *pyrE2*-based pop-in pop-out homologous recombination strategy as previously described [103] with modifications. In this strategy, a pre-deletion plasmid (pJAM4465) was generated by ligation of a PCR-amplified DNA fragment comprising ∼500 bp regions flanking the 5′ (upstream) and 3′ (downstream) ends of the target gene (*fdr*) into the BamHI to HindIII sites of plasmid vector pTA131. The pre-deletion plasmid served as the template for inverse PCR, using primers that removed the target gene while preserving the neighboring genes. The inverse PCR product was treated with DpnI to remove the methylated DNA template, gel-extracted, and ligated using the KLD Enzyme Mix (New England Biolabs, Ipswich, MA) according to the manufacturer’s instructions to generate the deletion plasmid pJAM4467.

*H. volcanii* H1207 served as the parent strain for the generation of the KT05 Δ*fdr* mutant strain. H1207 was transformed with the Δ*fdr* deletion plasmid (pJAM4467) and plated on Hv-Ca⁺ medium (uracil-free medium) to select for integration of the deletion plasmid (pop-in event). Plasmid integration was verified via colony PCR. Transformants were then cultured in 5 mL ATCC974 medium supplemented with 50 µg·mL^-1^ 5-fluoroorotic acid (5-FOA) (prepared from a 50 mg·mL^-1^ stock dissolved in DMSO) in 13 × 100 mm culture tubes. Cultures were incubated at 42 °C in the dark with orbital shaking at 200 rpm for 3–4 days to promote plasmid excision (pop-out event). Serial dilutions (10^-2^ to 10^-6^) of these cultures were plated on ATCC974 solid medium (1.5% w/v agar) supplemented with 50 µg·mL^-1^ 5-FOA. Colonies were screened by PCR to identify successful deletion mutants. Confirmed deletion strains were further streaked for isolation on 5-FOA-containing ATCC974 plates four consecutive times to ensure genetic homogeneity. Final verification of the deletion event was performed via PCR using check primers located approximately 700 bp upstream and downstream of the targeted gene locus.

### Generation of the *H. volcanii* His-FdR expression strain

The *fdr* gene was isolated by PCR using *H. volcanii* gDNA as template and a primer pair that included NdeI and BlpI sites (FdR - pJAM503 NdeI and FdR - pJAM503 BlpI, **Supplemental Table S4**). The PCR product and pJAM503 plasmid backbone (which encodes an N-terminal His-tag) were digested with NdeI and BlpI. The plasmid backbone was additionally treated with Antarctic phosphatase to prevent self-ligation. DNA fragments were purified from agarose gel slices with DNA gel extraction kit or PCR/DNA cleanup kit. Ligation of digested insert and vector was carried out at room temperature using a 2x Quick Ligase. The resulting His_6_-*Hv*FdR expression plasmid (pJAM3944) was first transformed into *Escherichia coli* Top10 with selection of transformants on LB medium supplemented with ampicillin (100 μg·mL^-1^). The plasmid was isolated and confirmed by DNA sequencing prior to transformation into *H. volcanii* KT05 [104, 105] with selection of transformants on ATCC974 supplemented with novobiocin (0.2 μg·mL^-1^).

### Generation of the *H. volcanii* Fdx-His expression strain

Generation of Fdx-His expression strain was performed as previously described (Weber *et al*., submitted). The *fdx* gene was amplified from *H. volcanii* genomic DNA by PCR using primers containing NdeI and NotI restriction sites. The PCR product and pET24b vector were digested with NdeI and NotI, ligated, and transformed into E. coli DH5α, generating pJAM4459. To generate a *H. volcanii* expression construct while retaining the native C-terminal His_6_ tag, *fdx* was PCR amplified from pJAM4459 using primers containing NdeI and NotI sites. The PCR product and pTA963 vector were digested with NdeI and NotI to remove the N-terminal His6 tag from pTA963, ligated, and transformed into E. coli DH5α, generating pJAM4455. Plasmids were isolated from LB medium supplemented with ampicillin (100 µg·mL^−1^), verified by DNA sequencing, and transformed into *H. volcanii* H1207 with selection on Hv-Ca^+^.

### SDM construction

Site-directed mutagenesis (SDM) was performed using the reduce- recycle PCR (rrPCR) method as previously described [106]. The His_6_-*Hv*FdR expression plasmid pJAM3944 served as the template. Amino acid substitutions included *Hv*FdR K47A, Y323A, and Y323W. After verification in *E. coli* Top10, the resulting plasmids expressing the His_6_-*Hv*FdR variants were transformed into *H. volcanii* KT05 as described above.

### Protein modeling

Three-dimensional structural models of *Hv*FdR were generated using two independent computational prediction platforms: AlphaFold Server 3 [107] and Phyre2 [108]. The AlphaFold-predicted structure exhibited high confidence, with a predicted a Template Modeling (pTM) score of 0.94. To assess the impact of cofactor binding on structural predictions, models were generated with various ligand combinations. The *Hv*FdR model complexed with FAD showed a pTM score of 0.94 and an ipTM score of 0.98, which slightly increased to pTM 0.95 and ipTM 0.96 upon inclusion of both FAD and NADPH. Phyre2, employed for homology-based structural modeling, produced an *Hv*FdR model with 100% confidence. All structural visualizations and analyses were conducted using ChimeraX (version 1.9). Electrostatic surface properties were assessed using Coulombic surface coloring in ChimeraX [109], where negatively charged regions are depicted in red, positively charged regions in blue, and neutral regions in white. Structural alignments and root-mean-square deviation (RMSD) calculations were performed using the MatchMaker tool in ChimeraX.

### Expression and purification of His_6_-*Hv*FdR and *Hv*Fdx-His_6_

i. **Expression of His_6_-*Hv*FdR and variants.** *H. volcanii* KT05 strains expressing His_6_-*Hv*FdR wild type and K47A, Y323A, and Y323W variants were inoculated from 20% (vol/vol) glycerol stocks (−80°C) onto ATCC974 solid medium (1.5% (w/v) agar) supplemented with novobiocin (0.2 μg·mL^-1^) (+Nov), and incubated for 4-6 days at 42 °C. Isolated colonies were inoculated into ATCC974 liquid medium (+Nov) and cultivated with aeration (orbital shaking, 200 rpm) at 42 °C. For large-scale cultures, 100 mL starter cultures were grown for 2 days in ATCC974 liquid medium (+Nov).
ii. **Expression of *Hv*Fdx-His_6_.** Expression and purification of *Hv*Fdx-His_6_ were performed as previously described. *H. volcanii* H1207 expressing *Hv*Fdx-His_6_ (pJAM4455) strains were inoculated from 20% (vol/vol) glycerol stocks (−80°C) onto Hv-Ca^+^ solid medium (1.5% (w/v) agar). Isolated colonies were inoculated into Hv-Ca^+^ and cultivated with aeration (orbital shaking, 200 rpm) at 42 °C. For large-scale cultures, 100 mL starter cultures were grown for 2 days in Hv-Ca^+^.
iii. **Large scale cultures**. Aliquots (5–10 mL) of initial 100 mL cultures were used to inoculate 500 mL of respective fresh medium in 2.8 L Fernbach flasks. Cell density was monitored by measuring OD_600_, where 1.0 OD_600_ unit is equivalent to approximately 1×10^9^ colony-forming units (CFU)/mL [110]. Cultures were grown to stationary phase (OD_600_ of 1.2–1.6), and cells were harvested by centrifugation at 2,862 × *g* for 50 min at room temperature (Sorvall Evolution RC centrifuge, Fiberlite F9-4×1000y rotor). Cell pellets were stored at –80 °C until further use.
iv. **Cell lysis.** Cell pellets were resuspended at a ratio of 1 g (wet weight) per 5 mL lysis buffer composed of 20 mM HEPES [pH 7.5], 2 M NaCl, 20 mM imidazole, 5 mM 2-mercaptoethanol, 5 mM MgCl_2_, 4 mM CaCl_2_, DNase I (4 μg·mL), and cOmplete Mini EDTA-free Protease Inhibitor Cocktail (1 tablet per 10 mL lysis buffer) (cat. no. 1183617000, Roche). Cells were disrupted using a French press (Glen-Mills, NJ, USA) for a total of five passages (1,500–2,000 psi (kPa), minimum high-ratio of 140 for a piston 1” diameter). Cell lysate was clarified by centrifugation at 10,000 × *g* for 50 min at 4 °C (ThermoFisher Sorvall Legend XTR, Fiberlite F14-6×250 LE). The resulting supernatant was filtered through 0.45 μm bottle filters (PES membrane; cat. no. 25-232, GenClone). Clarified lysates were used immediately for downstream protein purification.
v. **Ni-NTA affinity chromatography and dialysis.** A Bio-Rad Biologic DuoFlow system was used for fast protein liquid chromatography (FPLC). The instrument was primed with filtered nanopure H_2_O at a flow rate of 1 mL·min^-1^for 10 min prior to column installation. A 1 mL HisTrap column (cat. no. 17524701, Cytiva) was washed with five column volumes of NPH_2_O and equilibrated with 10 column volumes of binding buffer (20 mM HEPES, pH 7.5, 2 M NaCl, 20 mM imidazole) at 1 mL·min^-1^. Clarified lysate was loaded at 0.5 mL·min^-1^. The column was then washed with 5 column volumes of binding buffer, followed by elution of bound His_6_-tagged proteins using 20 mM HEPES, pH 7.5, 2 M NaCl, and 500 mM imidazole at 1 mL·min^-1^. The column was subsequently washed with nanopure H_2_O and stored in 20% ethanol for reuse. The HisTrap column was stripped and recharged with 0.1 M NiSO_4_ according to manufacturer’s instructions. Elution fractions were dialyzed against 20 mM HEPES, pH 7.5, and 2 M NaCl overnight at 4 °C. Dialysis was performed using SnakeSkin Dialysis Tubing (cat. no. 68035, 3.5 kDa MWCO, ThermoFisher). The elution fractions were dialyzed using a buffer-to-sample volume ratio of 2 L of buffer per 1 mL of elution fraction.
vi. **Size exclusion chromatography (SEC).** His-*Hv*FdR and variant proteins were further purified by size exclusion chromatography using the Bio-Rad BioLogic DuoFlow chromatography system. Prior to loading, samples were filtered through a 0.22 µm PES syringe filter (Whatman Uniflo, Cytiva) to remove particulate contaminants. Superdex 75 10/300 GL column (GE Healthcare) was equilibrated with two column volumes (24 mL) of filtered nanopure H_2_O, followed by two column volumes of 20 mM HEPES [pH 7.5] 2 M NaCl, and freshly prepared 1 mM dithiothreitol (DTT). Approximately 500 µL of protein sample was loaded onto the column and eluted at a flow rate of 0.3 mL·min^-1^. Protein elution was monitored by UV absorbance at 280 nm (A_280_), and fractions were collected for analysis. Molecular weights were determined using the Gel Filtration Standards kit (cat. no. 1511901, Bio-Rad) including the following: bovine thyroglobulin (670 kDa), bovine Ɣ-globulin (158 kDa), chicken ovalbumin (44 kDa), horse myoglobin (17 kDa), and vitamin B12 (1.325 kDa). Molecular mass estimations were derived from the linear regression (R^2^ > 0.99) of the logarithmic values of molecular mass against the gel phase distribution coefficient (*K*_av_). *K*_av_ was calculated using the equation: *K*_av_=(*V*_R_−*V*_o_)/(*V*_c_−*V*_o_) where *V*_R_ represents the retention (elution) volume of the protein, *V*_o_ is the void volume of the column, and *V*_c_ is the geometric bed volume. Collected fractions were stored at 4 °C and, if necessary, concentrated using Vivaspin 500 centrifugal concentrators (cat. no. VS0111, Sartorius) according to the manufacturer’s instructions.
vii. **UV-Visible spectroscopy (UVVIS).** Purified His_6_-*Hv*FdR and variant proteins (2 mg·mL^-1^) in 400 μL of 20 mM HEPES [pH 7.5], 2 M NaCl buffer were loaded into a quartz cuvette (Spectrophotometer Cell Micro 16.50-Q-10/8.5 mm), and analyzed using Agilent BioTek Epoch 2 with the cuvette attachment from 200-600 nm. Characteristic maximum absorbance peaks of oxidized flavin cofactors were identified by maxima at A_370_ and A_460_ [111].
viii. **Protein concentration**. Concentration was determined by measuring UVVIS at A_280_ and calculating the protein extinction coefficient of His_6_-*Hv*FdR. The molar extinction coefficient (ε) was calculated based on the amino acid composition of the protein, specifically the number of tryptophan, tyrosine, and cysteine residues (eq. 1). For concentrations reported in mg·mL^-1^, the molecular weight of the protein was incorporated using where A_280_ is the absorbance at 280 nm, MW is the molecular weight of the protein, and ε_molar_ is the molar extinction coefficient (eq. 2).

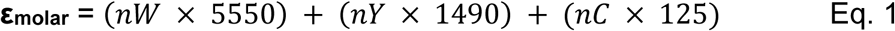

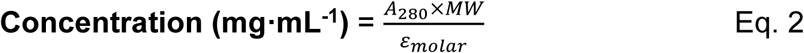

### TCA precipitation

His_6_-*Hv*FdR and variant proteins (1 μg) were diluted in nanopure H_2_O to a final volume of 100 µL in a screw cap 1.5 mL tube (cat. no. 3467TS, ThermoFisher). Ice-cold 80% trichloroacetic acid (TCA) was added to a final concentration of 10% (14.3 μL per 100 μL sample). The mixture was thoroughly mixed by gentle pipetting and incubated on ice overnight at 4 °C. Following incubation, samples were centrifuged at 16,000 × *g* for 10 min at 4 °C. The supernatant was carefully removed without disturbing the pellet. To wash the pellet, 1 mL of ice-cold 100% acetone was added, and the tube was gently inverted to mix. The sample was centrifuged again at 16,000 × *g* for 10 min at 4 °C. The supernatant was removed carefully to avoid disturbing the pellet, which was then air-dried at room temperature for 30 min. For resuspension, 1 µL of 1 M Tris-HCl [pH 8] was added per 10 µL of 1× Laemmli sample buffer (LSB) supplemented with 10% (v/v) 2-mercaptoethanol. A total of 11 µL of 1× LSB mixture was used to resuspend the pellet, with mixing performed by pipetting. The sample was then boiled for 10 min, briefly centrifuged for quick spin, and subsequently analyzed by SDS-PAGE.

### SDS-PAGE

Protein purity was determined by reducing 12% sodium dodecyl sulfate– polyacrylamide gel electrophoresis (SDS-PAGE) performed at 200 V for 50-60 min in Tris-glycine-SDS buffer [pH 8.3] composed of 25 mM Tris, 192 mM glycine, and 0.1% SDS. Precision Plus Protein Kaleidoscope molecular mass markers (Bio-Rad) were used as the molecular weight standard. SDS-PAGE gels were stained with Coomassie Brilliant Blue G-250 (0.2 g in 40% (v/v) ethanol and 5% (v/v) acetic acid) for 1 h and destained overnight in deionized water at room temperature. Gels were imaged using the iBright FL 1000 imaging (ThermoFisher Scientific) under the visible protein detection setting.

### HPLC analysis of flavin content

Flavin content was determined as previously reported [112]. To extract the cofactor from freshly purified His_6_-*Hv*FdR, 100 µL of a solution with 6.26 µM protein was combined with 500 µL of a methanol:dichloromethane (9:10, v/v) mixture in a 1.5-mL microcentrifuge tube. The mixture was vortexed at high speed for 60 sec, followed by the addition of 240 µL of 0.1 M ammonium acetate [pH 6] bringing the total volume to 840 µL. After a second round of vortexing for 60 sec, the sample was centrifuged at 16,000 x g for 5 min at 4°C to facilitate phase separation. The upper aqueous phase, which contained the extracted cofactor, was collected into a new 1.5 mL tube (typically yielding 400–500 µL solution). This extraction procedure was repeated twice to obtain a 700–800 µL aqueous phase in total. The pooled solution was filtered through a 0.45 µm pore-size membrane filter (Pall Acrodisc syringe filter; Pall Corporation, Port Washington, NY) and transferred to an HPLC vial. Cofactor content of the sample was analyzed with a 4.6 × 250 mm Vydac C18 reverse-phase analytical column (Separation Group, Hesperia, CA) on a Shimadzu Prominence HPLC, equipped with LC-20AD dual pumps, SIL-20A autosampler, SPD-M20A diode array detector, and CBM 20A controller system (Shimadzu Scientific Instruments, Columbia, MD). Separation was performed using a binary solvent system consisting of solvent A (2% acetonitrile in 27.5 mM sodium acetate buffer [pH 4.7]) and solvent B (100% acetonitrile), delivered at a flow rate of 0.6 mL·min^-1^. After sample application under 100% solvent A and an isocratic flow of 100% solvent A for 2 min that followed, linear gradients to 2% solvent B from 2 to 6 min, to 10% B from 6 to 15 min, and finally to 100% B from 15 to 18 min, were applied in that sequence. The column was maintained at 100% solvent B from 18 to 21 min before returning to 0% B for re-equilibration from 21 to 24 min. Elution was monitored at 450 nm, and UV-visible spectra of the compounds in the identified peaks were collected automatically using a diode array detector. Identification of the flavin eluting from the column was based on comparison of respective chromatographic retention time and UV–visible absorbance spectrum with those of commercially available flavin mononucleotide (FMN) and flavin adenine dinucleotide (FAD) (Thermo Fisher Scientific). Quantification of FAD in the sample was performed by using a calibration curve generated with known quantities of FAD.

### Complementation assay

H1207, (H1207 Δ*fdr*), KT05/pJAM3944 (His_6_-*Hv*FdR), KT05/pJAM202c (empty vector or EV), and H1207/EV were inoculated from 20 % (v/v) glycerol stocks (-80 °C) onto ATCC974 rich medium supplemented with 1.5 % (w/v) agar. Strains carrying plasmids were additionally supplemented with novobiocin (0.2 μg·mL^-1^) for selection. Plates were incubated at 42 °C for 5 days. Single colonies were cultured with 5 mL ATCC974 medium in rotating culture tubes (13 × 100 mm). Cells were grown to log phase (OD_600_ 0.5-0.6) and sub-cultured to OD_600_ 0.02 (13 x 100 mm, tubes) with 5 mL ATCC974 medium and grown to log phase. Cells were sub-cultured again to OD_600_ 0.02 with 1 mL respective medium in 1.5 mL Eppendorf tubes. In a 96- well CellPro cell culture plate (Alkali Scientific, FL), 150 µL of subculture was aliquoted into three replicate wells. Each strain included three biological and three technical replicates. Using the BioTek Epoch 2 microplate reader and Gen5 software (Agilent, Santa Clara, CA), cell growth was measured as follows: OD_600_ was measured every 15 min for 99 h, with aeration (double orbital continuous shaking), and temperature setpoint 42 °C. No inoculum controls were included to assess potential background signals from the medium alone.

### Measuring NADPH and NADP^+^ levels and H_2_O_2_ production in cell lysate

i. **Cell growth.** *H. volcanii* H1207, KT05, KT05/pJAM3944, KT05/EV, and H1207/EV were plated from 20% (v/v) glycerol stocks (−80 °C) onto ATCC974 rich medium supplemented with 1.5% (w/v) agar. Strains carrying plasmids were additionally supplemented with novobiocin (0.2 μg·mL^-1^) for selection. Single colonies were inoculated into 5 mL ATCC974 liquid medium in rotating culture tubes (13 × 100 mm). Cells were grown in rotating culture tubes to mid-log phase (OD_600_ of 0.6) and subsequently sub-cultured to an initial OD_600_ of 0.02 (13 × 100 mm, tubes) in fresh ATCC974 medium. Growth was at 42 °C.
ii. **Cell quenching.** All cells were quenched at mid-log phase (OD_600_ of 0.5-0.6) using ice cold 18% salt water. The 18% salt water was placed at -80 °C for 20 min until a slushy consistency was achieved. When the liquid started to solidify, it was shaken to create a slurry, which was stored at -20 °C and utilized within 2 h. In a 15 mL falcon tube, 1 mL of cell culture and 1.5 mL of ice slurry were added together for a final volume of 2.5 mL on ice and inverted twice. The mixture was centrifuged (4,000 × *g*, 10 min, 1 °C), the supernatant was removed, and the pellets were stored at −80 °C until analysis.
iii. **Cell lysis.** Frozen cell pellets were resuspended in 300 μL of ice cold (slurry consistency) 1× PBS [pH 7.4] by gentle pipetting, followed by addition of 300 μL of 0.2 N NaOH containing 1% (w/v) dodecyltrimethylammonium bromide (DTAB) with gentle pipetting to mix. Lysates were incubated on ice for 20 min and briefly centrifuged (4,000 × *g*, 10 min, 4 °C) to remove cellular debris. Clarified lysate was carefully transferred to a clean 1.5 mL tube for analysis. Protein concentration (mg·mL^-1^) was determined by NanoDrop (Thermoscientific NanoDrop One) in triplicate and averaged.
iv. **Measuring NADPH and NADP^+^ levels in cell lysate.** Cellular NADP(H) levels were quantified using the NADP/NADPH-Glo Assay (Promega; cat. no. G9081) with modifications. Reaction volumes were reduced to 1/4 of the manufacturer’s recommended volume, as 12.5 μL of cell lysate or NADP^+^ standard was dispensed into each well of a 384-well plate (Corning; cat. no. CLS3576) kept on ice. The protocol was followed according to the manufacturer’s protocol using 6.25 μL 0.4 M HCl and 6.25 μL 0.5 M Trizma for NADP^+^ detection, and 12.5 μL HCl/Trizma mixture for NADPH detection. To start the reaction, 25 μL of NADP/NADPH-Glo detection reagent was added to each well. Plates were analyzed using an Agilent BioTek Synergy H1 plate reader with the following settings: orbital shaking for 30 s, incubation for 20 min at RT, followed by luminescence measurements collected every 2.5 min over a 1 h period. NADP⁺ standards were prepared as recommended by diluting a 2 mM stock solution to 10 μM stock solution, followed by serial dilution to final reaction concentrations of 800, 400, 200, 100, 50, 25, 12.5 and 0 nM NADP⁺. Standards were processed identically to experimental samples. Two independent experiments were performed using three biological replicates and technical replicates from the values at ∼ 58 and 60 min. NADP(H) values were normalized to total protein (µg) added to the reaction. Ratios were calculated from each technical replicate and averaged. Controls with buffer only and cell lysate only were also analyzed.
v. **Measuring H_2_O_2_ production in cell lysate.** H_2_O_2_ levels produced by cell lysate were quantified using the ROS-Glo H_2_O_2_ Assay (Promega; cat. no. G8820) with modifications. Reaction volumes were reduced to from the manufacturer’s recommended volume, yielding a total cell lysate volume of 25 μL per reaction. For luminescence measurements, 25 μL of cell lysate was dispensed into each well of a 384-well plate (Corning; cat. no. CLS3576), followed by addition of 6.25 μL of H_2_O_2_ Substrate according to the manufacturer’s protocol. Immediately after, a 1:1 ratio of ROS-Glo Detection Solution (31.25 μL) prepared according to the manufactures protocol was added to the plate. Plates were analyzed using an Agilent BioTek Synergy H1 plate reader with the following settings: orbital shaking for 30 s, incubation for 10 min at RT, followed by luminescence measurements collected every 2.5 min over a 1 h period. H_2_O_2_ standards were prepared as recommended by diluting a 100 mM stock solution to 1 μM, then serially diluting to final concentrations of 800, 400, 200, 100, 50, 25, 12.5 and 0 nM H_2_O_2_. Standards were processed identically to experimental samples. Experiments were performed using three biological replicates, each measured with three technical replicates. Two independent experiments were performed using replicates from the values at ∼ 58 and 60 min. H_2_O_2_ values were normalized to total protein (µg) in the cell lysate. Controls included buffer only and no addition of H_2_O_2_. The substrate provided by the manufacturer’s kit was also analyzed.

### Aerobic NAD(P)H oxidase enzymatic assay development

Aerobic enzymatic assays were performed in clear flat-bottom 96-well plates in a final reaction volume of 150 µL unless otherwise specified. Reactions were prepared in the indicated assay buffer conditions being tested and contained purified His_6_-*Hv*FdR with 150 µM NAD(P)H unless otherwise specified. All reactions were initiated by the addition of saturating NAD(P)H. Control reactions containing only assay buffer, NAD(P)H in assay buffer, or purified His_6_-*Hv*FdR in assay buffer were included under each tested condition to verify that absorbance changes resulted from enzymatic activity rather than spontaneous cofactor oxidation or buffer effects. NAD(P)H oxidation was monitored spectrophotometrically at A_340_ every minute for 30 min using a BioTek Epoch2 plate reader. Activity was quantified using the extinction coefficient of NADPH at A_340_ (ε_340_ = 6200 M^-1^·cm^-1^).

i. **Optimization of buffer conditions for *Hv*FdR activity**. Reactions containing 1.5 µM FdR and 150 µM NAD(P)H were tested under varying buffer, pH, salt, temperature and storage conditions (**Supplemental Fig. S7**). Prior to assays, proteins were buffer exchanged into the indicated conditions using Zeba Spin desalting columns (cat. no. 89877; Thermo). For pH optimization, reactions were performed in 20 mM MES [pH 6], HEPES [pH 7, 7.5, and 8], Tris-HCl [pH 8 and 9], CAPSO [pH 9 and 10], or CAPS [pH 11] supplemented with 2 M NaCl at 42°C. Following identification of the optimal pH, salt dependence was evaluated using NaCl (0, 0.05, 0.5, 1, 2, 3, and 4 M) or KCl (0, 0.05, 0.5, 1, 2, and 3 M). Salt-dependent stability was assessed by incubating samples at 4 °C for 24, 48, or 72 h prior to activity measurements. Temperature optimization was performed at 24, 33, 42, 51, 60, and 65 °C under optimal buffer and salt conditions. Thermal stability was further evaluated by incubating purified FdR for 1 h at the indicated temperatures using a BioRad T100 Thermocycler in hold mode prior to activity measurements.
ii. **Determination of optimal enzyme concentrations for oxidase reaction.** Varying concentrations of *Hv*FdR were incubated with 0–250 µM NAD(P)H. NADPH-dependent reactions contained 0.5, 1.5, or 2.5 µM FdR in 20 mM HEPES [pH 8] and 4 M NaCl, whereas NADH-dependent reactions contained 1.5, 3.0, or 6.0 µM FdR in 20 mM MES [pH 6] and 3 M NaCl. Reactions were monitored at A_340_ every minute for 30 min at 51 °C. H_2_O_2_ production was quantified by ABTS oxidation as described below.

### Kinetic analysis

Specific activity was calculated as nmol substrate oxidized per minute per mg of enzyme (nmol·min^-1^·mg^-1^), and kinetic parameters were determined by Michaelis–Menten and Lineweaver–Burk analyses. Steady-state kinetic parameters were determined from the initial linear portion of each reaction, where less than 20% of the NAD(P)H substrate was consumed. The Michaelis–Menten curve represents the nonlinear least-squares fit of the experimental data to the Michaelis–Menten equation, *v* = *V*_max_[*S*]/(*K*_m_ + [*S*]). Nonlinear regression was performed using the Solver add-in in Microsoft Excel [113].

### NAD(P)H oxidase hydrogen peroxide (H_2_O_2_) quantification

H_2_O_2_ generated from FdR-dependent NAD(P)H oxidation was quantified via ABTS oxidation, catalyzed by horseradish peroxidase (HRP), measured at A_412_ (38). In this reaction, reduced ABTS (colorless) is oxidized to a green-colored radical cation (ABTS^•^⁺) through a one-electron transfer mediated by H_2_O_2_ (H_2_O_2_+ ABTS + HRP → H_2_O + ABTS^•^⁺). Once NAD(P)H oxidation ceased, reactions were diluted sixfold (25 µL reaction + 125 µL diluent, final volume 150 µL) using 20 mM HEPES [pH 8], and 1.6 M NaCl to yield a final buffer composition of 20 mM HEPES [pH 8], and 2 M NaCl for NADPH-dependent reactions, or 20 mM HEPES [pH 6], and 1.8 M NaCl to yield a final buffer composition of 20 mM MES [pH 6] and 2 M NaCl for NADH-dependent reactions. For H_2_O_2_- dependent ABTS oxidation, 3 µL of 10 mM ABTS (final concentration: 0.2 mM) in H_2_O was added in triplicate to each diluted reaction. To initiate the reaction, 3.8 µL horseradish peroxidase (1 µg·µL^-1^) in H_2_O was added immediately. The reaction was carried out in a 96-well plate at 51 °C with linear shaking (fast speed, 10 seconds), before A_412_ was measured. H_2_O_2_ concentrations were derived from a standard curve.

A standard curve for H_2_O_2_ quantification was generated using a 30% H_2_O_2_ stock solution (9.8 M; cat. no. H1009, Sigma-Aldrich), diluted to final concentrations ranging from 0-80 µM in 20 mM HEPES [pH 8], and 2 M NaCl, or 0-60 µM in 20 mM MES [pH 6], and 2 M NaCl. In a 96-well plate 150 µL of each H_2_O_2_ standard solution was combined with 3 µL of 10 mM ABTS in triplicate. Reactions were initiated by the addition of 3.8 µL of horseradish peroxidase (1 µg·µL^-1^). The plate was incubated at 51 °C with linear shaking (fast speed, 10 seconds), before ABTS oxidation was measured at A_412_.

### NADPH oxidase assay of putative flavin binding variants

His_6_-*Hv*FdR and putative flavin-binding variants (K47A, Y323A, Y323W, K47A/Y323A, and K47A/Y323W) were purified by Ni-NTA affinity chromatography and SEC using the methods described above and analyzed for NADPH oxidase activity under aerobic conditions. Flavin incorporation was quantified by normalizing A_460_, corresponding to the FAD isoalloxazine peak, to the total protein concentration (mg·mL^-1^) determined by UV– visible spectroscopy. This normalization accounts for protein concentration differences and allows comparison of relative flavin occupancy across variants. For each sample, a normalized flavin (FAD) value was calculated as: A_460_/protein concentration (mg·mL^-1^). This generates a unit of absorbance per mg protein, reflecting relative FAD content per protein concentration. Fold changes of flavin incorporation were then calculated relative to WT using: Unit_Variant_ / Unit_WT_. A fold value of >1 indicates increased flavin occupancy relative to WT, whereas a value of <1 indicates reduced incorporation. As catalytic turnover depends on flavin occupancy, enzyme concentrations used in kinetic assays were adjusted based on these fold-change values compared to WT. Specifically, if a variant exhibited reduced flavin occupancy compared to WT, a proportionally higher amount of protein was added to reactions to maintain an equivalent concentration of FAD-bound active sites. Variants with increased flavin incorporation required proportionally less protein to achieve comparable active enzyme concentrations. This normalization ensured that observed differences in catalytic activity reflected intrinsic kinetic effects of the mutations rather than variations in flavin loading.

Buffer exchange was performed using Zeba Spin desalting columns into the optimal conditions, 20 mM HEPES [pH 8], and 4 M NaCl. Reactions were prepared with the normalization (µM) of His_6_-*Hv*FdR and variant proteins in 20 mM HEPES [pH 8] and 4 M NaCl using varying concentrations of saturating NADPH (0–250 µM in H_2_O). NADPH was added to initiate the reaction. NADPH oxidation was monitored at A_340_ every minute for 30 min at 51 °C. H_2_O_2_ production was quantified via ABTS oxidation as outlined above.

### ThermoFAD

Buffer exchange of His_6_-*Hv*FdR and variant proteins was performed using Zeba Spin desalting columns into 20 mM HEPES [pH 8.0] buffer containing 4 M NaCl. The purified proteins were adjusted to a final concentration of 0.5 mg·mL^-1^ in 20 μL. To assess ligand-dependent stabilization, proteins were supplemented with 150 μM NADP^+^. Samples were loaded into a 96-well plate (iCycler IQ PCR plates, cat. no. 2239441, Bio-Rad), with buffer-only samples included as negative controls. Thermal melting point assays were performed using a C1000 thermal cycler equipped with a CFX96 real-time detection (Bio-Rad Laboratories). Samples were heated from 20 °C to 90 °C at an incremental rate of 0.5 °C per 10 sec, with fluorescence recorded once per step. Intrinsic flavin fluorescence was monitored using an excitation range of 470–500 nm and a SYBR Green emission filter set, which overlaps with the emission range of the flavin isoalloxazine [74, 114].

### Microscale thermophoresis (MST)

Binding affinity measurements for FdR were performed using the His-Tag Labeling Kit RED-tris-NTA 2nd Generation (Cat. no. MO- L018, NanoTemper Technologies) and premium capillaries (cat. no. MO-K025) according to the manufacturer’s protocol. Following the manufactures instructions to assess the affinity of the fluorescent dye for the target protein, 4 µM WT FdR in 20 mM HEPES [pH 7.5] and 2 M NaCl was subjected to a 1:1 serial dilution in the same buffer, yielding a final concentration range down to 0.12 nM. A fixed concentration of 25 nM RED-tris-NTA dye (from a 50 nM stock) was added to each dilution. The protein–dye mixtures were incubated at room temperature for 30 minutes before loading into capillaries. Measurements were conducted using 40% LED/excitation power and medium MST power. The resulting *K*_d_ (dissociation constant) was used to determine the optimal protein-to-dye ratio for ligand-binding assays.

To evaluate the binding affinity of WT FdR for target ligands, serial dilutions of each ligand were prepared in 20 mM HEPES [pH 7.5] and 2 M NaCl. A starting concentration of 0.5 µM FdR labeled with 50 nM RED-tris-NTA dye (final concentration of 0.25 µM FdR and 25 nM Red-tris NTA dye). To note, a final concentration once added to the ligand is 0.125 µM FdR. Dye was removed by centrifugation (10 min, 14 x g, 4C). Following the manufacturer’s instructions, for NADP⁺, a starting concentration of 2,500 µM was serially diluted 1:1 to a final concentration of 0.15 µM. Capillaries were then loaded and analyzed at 40% LED/excitation and medium MST settings. The 6^th^, 7^th^, and 8^th^ runs were used for analysis. The same procedure was followed for NAD⁺, with a starting concentration of 10,000 µM serially diluted 1:1 to a final concentration of 1.22 µM. Capillaries were then loaded and analyzed at 80% LED/excitation and medium MST settings.

### Anaerobic electron transfer assays

All reactions were performed under strict anaerobic conditions (98-99% N_2_ and 1-2% H_2_) at 25-27 °C. All buffers and enzymes were gas exchanged in 100% argon gas before entrance into the Coy anaerobic chamber. UV-visible absorbance spectrum (200–800 nm) (UVVIS) were collected with a quartz cuvette (1 cm path length) (Spectrophotometer Cell Micro 16.50-Q-10/8.5 mm) using Agilent BioTek Epoch 2 with the cuvette attachment, unless specified.

i. **Midpoint potential.** The value of the midpoint redox potential (*E*_0_′) of *Hv*FdR- bound FAD was determined by anaerobic reductive titration using sodium dithionite (SD) as the reductant and methyl viologen (MV^2+^) as a redox reference dye. First a series of reaction mixtures (400 µL) containing purified His_6_-*Hv*FdR (20 μM), 10 μM MV^2+^, 20 mM HEPES buffer [pH 7.5] and 2 M NaCl was place in 1.5 mL microfuge tubes. Then to these tubes freshly prepared sodium dithionite (SD) solutions of varying concentrations but fixed volumes were added to achieve a final concentration range of 0 - 11 μM and to reduce the FAD cofactor and the reference dye at increasing degrees. After the reaction mixtures were incubated for 30 min at 27 °C, the respective UVVIS were collected and processed to obtain the absorbance values at A_460_, which is characteristic of oxidized flavin chromophore, and at A_604_ for the MV^•+^ radical; the oxidized form (MV^2+^) shows negligible absorbance at A_604_. The UVVIS of the control solutions containing 10 μM MV^2+^ or MV^•+^, and 100 μM SD were recorded in triplicate to verify the spectral features of each component and to confirm the integrity of the SD dilution. The extinction coefficient of the oxidized form of His_6_-*Hv*FdR at A_460_ was determined to be 0.0084 μM^-1^·cm^-1^; the extinction coefficient of reduced methyl viologen radical at A_604_ is 0.012 μM^-1^·cm^-1^ [115, 116]. Using these extinction coefficient values, the concentrations of oxidized FAD in FdR and reduced methyl viologen with various SD concentrations were calculated using the following equations:

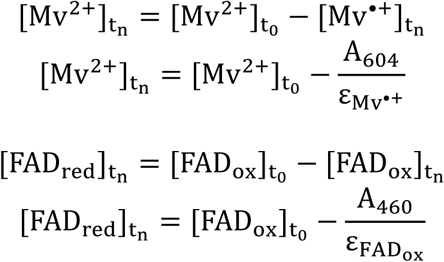 where *t_n_* represents the concentration of the compound at *n* μM SD, and *t_0_* represents the concentration of the compound at 0 μM SD. Then, these values were fitted to the Nernst equation to obtain the midpoint redox potential of His_6_- *Hv*FdR-bound FAD, considering that methyl viologen, the redox indicator dye (D), was in equilibrium with the flavin (F) [117].

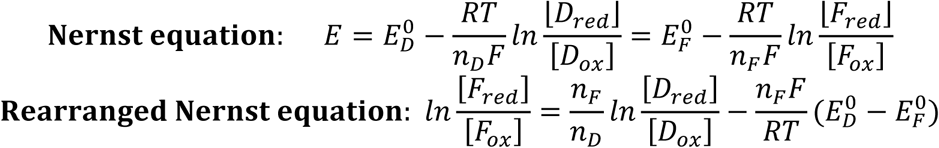 [From the plot of the rearranged Nernst equation: With the value of the slope, 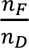, and n_D_ of 1 for MV^2+^/MV^•+^ pair, the electron transfer event for FAD (*n_F_*) was determined. The value of the intercept, 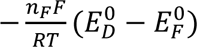, was used to calculate the midpoint redox potential value for FdR-bound FAD; *E*_0_′ MV^2+^/MV^•+^, -446 mV vs SHE; gas constant R, 8.314 J K^−1^·mol^−1^; temperature T, 298 K; Faraday’s constant F, 96,485 c·mol^-1^]
ii. **Molar extinction coefficient of *Hv*FdR.** The molar extinction coefficient (ε) of the flavin-bound His_6_-*Hv*FdR was determined at A_460_ by plotting absorbance versus protein concentration (0–40 μM) and applying a linear regression to obtain the line of best fit. Measurements were performed in 20 mM HEPES [pH 7.5] buffer containing 2 M NaCl, and monitored by UVVIS. The ε^460^ of the purified His_6_-*Hv*FdR was determined to be 6,050 M^−1^·cm^−1^.
iii. **Direction of electron flow to *Hv*FdR.** To assess the physiological direction of electron transfer, His_6_-*Hv*FdR (20 μM) was reduced with 10 μM sodium dithionite for 30 min, in 20 mM HEPES [pH 7.5] buffer containing 2 M NaCl. Excess reductant was removed using a Zeba desalting column following manufacturer’s instructions. Following enzyme reduction, NADP⁺ (20 μM) was added, and the reaction mixture (FdR_red_ → NADP^+^) was incubated for an additional 30 min. After incubation, the redox state of the reaction was monitored by UVVIS. Control samples included FdR_ox_, FdR_red_, NADP^+^, and NADPH only in 20 mM HEPES [pH 7.5] containing 2 M NaCl. A reverse reaction of NADPH → FdR_ox_ did not occur when 3:1, 2:1, 1:1, 1:2, and 1:3 FdR:NADPH ratios were incubated with DCIP (100 μM). Thus, establishing that increased concentrations of NADPH are needed to push electron transfer to FdR under the reaction conditions.
iv. **Diaphorase activity of *Hv*FdR.** To determine anaerobic enzyme kinetics of His_6_- *Hv*FdR, NADPH oxidation coupled to the reduction of the artificial electron acceptor 2,6-dichlorophenolindophenol (DCIP) was monitored. To a total reaction volume of 100 μL in a 96-well plate, oxidized His_6_-*Hv*FdR (1 nM) and DCIP (15 μM) were mixed in reaction buffer: 20 mM HEPES [pH 7.5] containing 2 M NaCl. To start the reaction, saturating NADPH (0 – 14 µM) was added. NADP(H) oxidation was monitored at A_340_ and DCIP reduction was monitored at A_600_, every minute for 30 min at 28 °C. The reaction was also analyzed at 42 and 51 °C. Other electron acceptors, hexacyanoferrate (III) (ferricyanide) and Nitro Blue Tetrazolium chloride (NBT), accepted electrons from FdR_red_, but DCIP was chosen for further experimentation due to its ease of readability. Non-optimal buffer conditions were also chosen to ensure accurate reading of oxidation-reduction of all species. Specific activity was calculated in nmol substrate oxidized or reduced per min per mg of enzyme (nmol·min^-1^·mg^-1^).
v. **Ferredoxin-dependent activity of *Hv*FdR.** To a total reaction volume of 100 μL in a 96-well plate, oxidized FdR (1 nM) and FerA5 Fdx (0-3.5 μM), were added together and incubated for 2 min at room temperature in a reaction buffer of 20 mM HEPES and 2 M NaCl. The artificial electron acceptor DCIP (15 μM) was added to the protein mix. To start the reaction, saturating NADPH (4 μM) was added. NADP(H) oxidation was monitored at A_340_ and A_600_ for DCIP reduction every minute for 30 minutes at 28 °C. Specific activity was calculated in nmol substrate oxidized or reduced per min per mg of enzyme (nmol·min^-1^·mg^-1^).

### Statistical analysis

All experiments were conducted in at least two independent trials and included three biological replicates with three technical replicates per condition. Data are presented as mean ± standard deviation (SD). Statistical significance was determined using Student’s *t*-test, with thresholds defined as *p* < 0.05, *p* < 0.005, or not significant (n.s.). All analyses were performed using Microsoft Excel.

## Supporting information

This article contains supporting information with citations: 30, 45, 47, 48, 50, 51, 118–121

## Funding and additional information

Funds awarded to JMF to advance archaeal biocatalysts were through the U.S. Department of Energy, Office of Basic Energy Sciences, Division of Chemical Sciences, Geosciences and Biosciences, Physical Biosciences Program (DE-FG02-05ER15650) and to advance cellular mechanisms were through the National Institutes of Health (R35 MIRA 1R35GM161171-01). USDA National Institute of Food and Agriculture (Hatch Project FLA-MCS-006312) funds to JMF also supported this project. Funds awarded to KRW through NASA Florida Space Grant (FSGC 80NSSC20M0093) to determine functional significance of post-translation modifications in haloarchaea.

## Competing interests

The Author(s) declare that there is no conflict of interest.

## Notes

### Competing Interest Statement

The authors have declared no competing interest.

