## Supplemental Table S1 for "Mechanistic insights into redox activity and catalytic determinants of the haloarchaeal flavin-dependent oxidoreductase *Hv*FdR"

**Supplemental Table S1.** Comparison of *H. volcanii Hv*FdR to NAD(P)H oxidase and ONFR enzymes by 3D-structural homology modeling.

| Organism | Protein | PDBe; UniProt ID | Pfam | AA | Seq. Id. | Prob. | E-value | RMSD (atom pairs) – Phyre2[a] | RMSD (atom pairs) – Alpha Fold[b] |
|---|---|---|---|---|---|---|---|---|---|
| *Haloferax volcanii* | *Hv*FdR | - D4GWJ4 | PF07992 PF18267 | 413 | - | - | - | 1.126 Å (232); 3.307 Å (408) | |
| **NAD(P)H oxidases:** | | | | | | | | | |
| *Lactobacillus sanfranciscensis* | Nox | 2cdu; Q9F1X5 | PF07992 PF02852 | 452 | 21.5 | 1.0 | 8.42e-34 | 0.980 Å (266); 3.150 Å (390) | 1.072 Å (222); 4.996 Å (389) |
| *Archaeoglobus fulgidus DSM 4304* | Nox-A3 | 6pfz; O29847 | PF07992 PF02852 PF00581 | 551 | 25.9 | 1.0 | 3.63e-34 | 0.973 Å (220); 5.117 Å (402) | 1.038 Å (250); 4.609 Å (397) |
| *Enterococcus faecalis* | Npr | 1npx; P37062 | PF07992 PF02852 | 447 | 25.5 | 1.0 | 5.59e-9 | 1.115 Å (188); 5.083 Å (390) | 1.131 Å (238); 5.538 Å (397) |
| *Streptococcus pyogenes* | NOXase | 2bc0; Q5XC60 | PF07992 PF02852 | 456 | 18.1 | 1.0 | 1.08e-29 | 1.023 Å (193); 4.567 Å (400) | 1.028 Å (214); 5.204 Å (402) |
| **ONFRs:** | | | | | | | | | |
| *Pseudomonas sp.* (strain KKS102) | BphA4 | 2gqw; Q52437 | PF07992 PF14759 | 408 | 22.7 | 1.00 | 5.61e-35 | 0.980 Å (266); 3.150 Å (390) | 1.072 Å (222); 4.996 Å (389) |
| *Novosphingobium aromaticivorans* | ArR | 3lxd; Q2GBV9 | PF07992 PF14759 | 415 | 25.8 | 1.00 | 1.04e-35 | 0.183 Å (378); 1.190 Å (404) | 1.110 Å (224); 3.376 Å (396) |
| *Pseudomonas putida* | Pdr | 1q1r; P16640 | PF07992 PF14759 | 422 | 22.7 | 1.00 | 1.12e-36 | 1.085 Å (297); 2.632 Å (404) | 0.998 Å (252); 3.175 Å (402) |
| *Rhodopseudomonas palustris* | PuR | 3fg2; Q6N3B2 | PF07992 PF14759 | 405 | 20.8 | 1.00 | 4.15e-35 | 0.954 Å (287); 2.690 Å (399) | 1.055 Å (257); 3.544 Å (400) |

*Hv*FdR Phyre2[a] and Alpha Fold[b] generated models compared to each other and to other crystalized homologs as indicated. Root Mean Square Deviation (RMSD) values based on pruned and total atom pairs calculated using ChimeraX. Sequence identity (Seq. Id.), probability (Prob.), and E-value metrics determined by comparison of *Hv*FdR AlphaFold model to FdR homologs using Foldseek. -, not applicable; PDBe, PDB entry; UniProt ID, accession number; AA, amino acid length. Phyre2 (confidence in the model: 100.0%). PF07992, pyridine nucleotide-disulfide oxidoreductase; PF18267, rubredoxin $NAD^+$ reductase C-terminal domain; PF02852, pyridine nucleotide-disulfide oxidoreductase, dimerization domain; PF00581, rhodanese-like domain; PF14759, reductase C-terminal domain.
