## Supplemental Table S2 for "Mechanistic insights into redox activity and catalytic determinants of the haloarchaeal flavin-dependent oxidoreductase *Hv*FdR"

**Supplemental Table S2.** UV-visible spectra of $His_6$-*Hv*FdR and variant proteins.

| $His_6$-*HvFdR* | $A_{280}$ | mg/mL | Normalization $A_{460}$/(mg/mL) | Fold-change WT/Variant | µM for reaction |
|---|---|---|---|---|---|
| WT | 2.476 | 2.005 | 0.235/2.005 = 0.1171 | - | 1.50 |
| K47A | 2.447 | 1.961 | 0.275/1.961 = 0.1402 | 0.835 | 1.25 |
| Y323A | 2.480 | 2.008 | 0.156/2.008 = 0.0776 | 1.508 | 2.26 |
| Y323W | 2.445 | 1.980 | 0.232/1.980 = 0.1171 | 1.000 | 1.50 |
| K47A Y323A | 2.466 | 1.997 | 0.199/1.997 = 0.0996 | 1.175 | 1.76 |
| K47A Y323W | 2.464 | 1.995 | 0.218/1.995 = 0.1092 | 1.074 | 1.61 |

Absorbance spectra were collected to determine relative flavin binding and fold changes in flavin occupancy for each $His_6$-*Hv*FdR protein. Quantitative analysis of spectral features was used to calculate the micromolar amount of flavin required for subsequent enzymatic assays.
