## Supplemental Table S3 for "Mechanistic insights into redox activity and catalytic determinants of the haloarchaeal flavin-dependent oxidoreductase *Hv*FdR"

**Supplemental Table S3.** List of strains and plasmids used in this study.

| Name | Description | Ref. |
|---|---|---|
| **Strain** | | |
| Top10 | $F^-$ *mcrA* Δ*(mrr-hsdRMS-mcrBC) Φ80lacZΔM*15 Δ*lacX74 recA1 araD139* Δ*(ara leu)* 7697 *galU galK rpsL* ($Str^r$) *endA1 nupG λ-* | Invitrogen |
| GM2163 | F⁻ *ara-14 leuB6 fhuA31 lacY1 tsx78 glnV44 galK2 galT22 mcrA dcm-6 hisG4 rfbD1 rpsL136 dam13::*Tn*9 xylA5 mtl-1 thi-1 mcrB1 hsdR2* | New England Biolabs |
| DS70 | wild-type isolate DS2 cured of plasmid pHV2 | [118] |
| H26 | DS70 *ΔpyrE2* | [119] |
| H1207 | H26 Δ*pyrE2* $pitA_{Nph}$ Δ*mrr* | [120] |
| KT05 | H1207 Δ*fdr* | This study |
| **Plasmids** | | |
| pET24b | $Kan^r$; pBR322-derived plasmid containing C-terminus $P_{T7}$-6xHis tag | Novagen |
| pTA131 | $Amp^r$; pBluescript II containing $P_{fdx}$-*pyrE2* | [119] |
| pJAM503 | $Amp^r$; $Nov^r$; *H. volcanii*-*E. coli* shuttle vector with coding sequence for N-terminal $His_6$ tag (fused to N-terminus of HVO_0850) | [121] |
| pTA963 | $Amp^r$; p.*tnaA* with N-terminal $His_6$-tag *pyrE2*$^+$*hdrB*$^+$ | [120] |
| pJAM4459 | $Kan^r$; pET24b carrying Fdx-$His_6$ | This study |
| pJAM4455 | $Amp^r$; pTA963 with N-terminal $His_6$ replaced with Fdx-$His_6$ from pJAM4459 | Weber *et al.* submitted |
| pJAM202c | $Amp^r$; $Nov^r$; *H. volcanii*-*E. coli* shuttle $P2_{rrn}$ empty vector | [121] |
| pJAM3944 | $Amp^r$; $Nov^r$; *H. volcanii*-*E. coli* shuttle $P2_{rrn}$ $His_6$-FdR expression | This study |
| pJAM4465 | $Amp^r$; pTA131-based *fdr* pre-deletion plasmid | This study |
| pJAM4467 | $Amp^r$; pTA131-based Δ*fdr* deletion plasmid | This study |
| pJAM4530 | $Amp^r$; $Nov^r$; *H. volcanii*-*E. coli* shuttle $P2_{rrn}$ $His_6$-FdR K47A expression | This study |
| pJAM4531 | $Amp^r$; $Nov^r$; *H. volcanii*-*E. coli* shuttle $P2_{rrn}$ $His_6$-FdR Y323A expression | This study |
| pJAM4532 | $Amp^r$; $Nov^r$; *H. volcanii*-*E. coli* shuttle $P2_{rrn}$ $His_6$-FdR Y323W expression | This study |
| pJAM4533 | $Amp^r$; $Nov^r$; *H. volcanii*-*E. coli* shuttle $P2_{rrn}$ $His_6$-FdR K47A Y323A expression | This study |
| pJAM4534 | $Amp^r$; $Nov^r$; *H. volcanii*-*E. coli* shuttle $P2_{rrn}$ $His_6$-FdR K47A Y323W expression | This study |
