## Supplemental Table S4 for "Mechanistic insights into redox activity and catalytic determinants of the haloarchaeal flavin-dependent oxidoreductase *Hv*FdR"

**Supplemental Table S4.** List of primers used for gene deletions, expression plasmids, and site-directed mutagenesis (SDMs).

| Primer | Sequence (5' to 3') | Source |
|---|---|---|
| Fdx-His$_6$ - pET24b NdeI | TTGCATATGCCCACGGTAACCTACCTC | This study |
| Fdx-His$_6$ - pET24b XhoI | TTTCTCGAGGATGACGCGGTTCTGC | This study |
| Fdx-His$_6$ - pTA963 NdeI | TTGCATATGCCCACGGTAACCTACCTC | This study |
| Fdx-His$_6$ - pTA963 NtoI | TAAGCGGCCGCTGGCAGCAGCCAACTC | This study |
| FdR knock-in BamHI | TAAGGATCCGGAAAACGACTTCGGTCCCGG | This study |
| FdR knock-in HindIII | TTTAAGCTTTCCTCTCCTCGCGGGGCAG | This study |
| FdR inverse knock-out F | GGACCCGCTCCCGCTGTTTTC | This study |
| FdR inverse knock-out R | CGACGTTTGTATTCGGGGTTAATGTGG | This study |
| FdR 700 bp Δ check F | CGACGTTTGTATTCGGGGTTAATGTGG | This study |
| FdR 700 bp Δ check R | GGACCCGCTCCCGCTGTTTTC | This study |
| FdR - pJAM503 NdeI | GGTCATATGAGCCAATCGTACGTGATCGTCGGC | This study |
| FdR - pJAM503 BlpI | TTAGCTCAGCTTACTGTTCGGCCGCCG | This study |
| FdR SDM Anchor R1 | CGTAGCGCCACCACGCCTTGCCGCG | This study |
| FdR K47A F2 | CAACCGCATTCTCATC**gcg**GAATTCGCCAAGGGC | This study |
| FdR SDM Anchor F2 | CACGAGGCGCTCCGCGAGCGCAACG | This study |
| FdR Y323A R3 | GTGGGTGATGGA**ggc**CGACGAGACCCAGC | This study |
| FdR Y323W R3 | GTGGGTGATGGA**cca**CGACGAGACCCAGC | This study |

Primer sequence: underlined nucleotides represent restriction modification sites and lowercase nucleotides represent site-directed mutations.
