## Supplemental Figures S1-S7 for "Mechanistic insights into redox activity and catalytic determinants of the haloarchaeal flavin-dependent oxidoreductase *Hv*FdR"

**A**

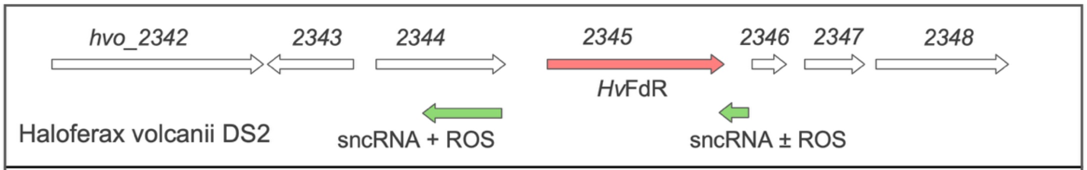


| Gene locus tag | Pfam | Description |
|---|---|---|
| HVO_2342 | PF01554 | MATE-family multidrug and toxin efflux transporter |
| HVO_2343 | PF19646 | Uncharacterized, DUF6149 |
| HVO_2344 | PF24038<br>PF24042 | HTH domain protein, DUF7347 and DUF7351 |
| HVO_2345 | PF07992<br>PF18267 | Flavin-dependent oxidoreductase (*Hv*FdR) |
| HVO_2346 | PF25934 | Uncharacterized, recommended DUF7979 |
| HVO_2347 | PF23439 | Uncharacterized, DUF7124 |
| HVO_2348 | PF02649 | GTP cyclohydrolase, MptA homolog |

**B**

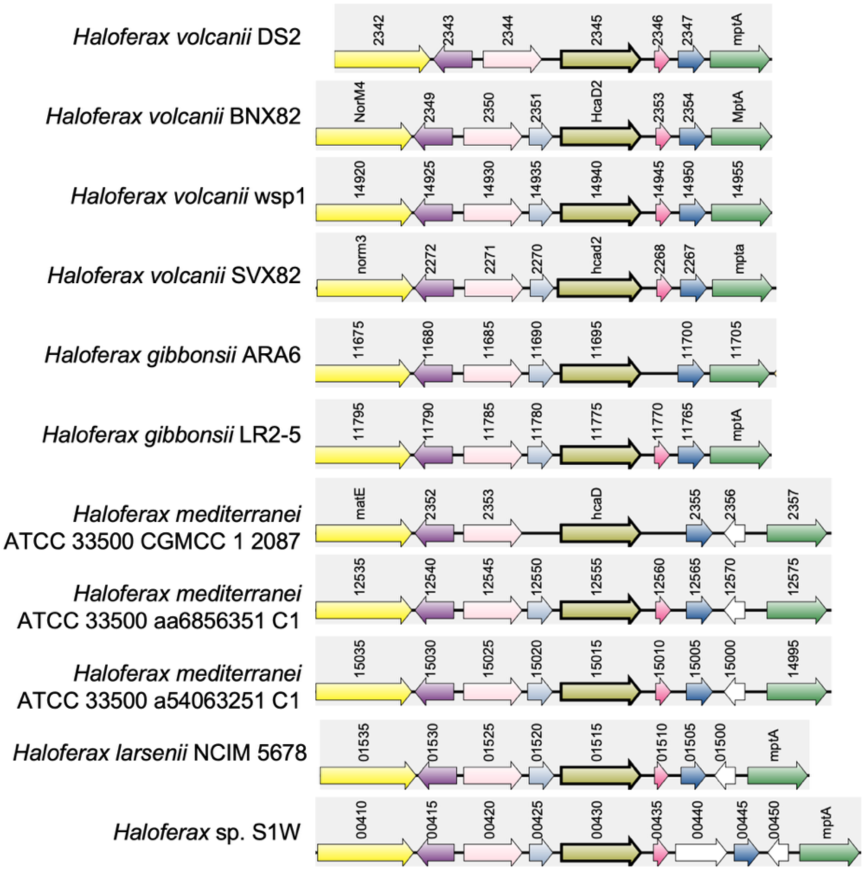


**Supplemental Figure S1.** Genomic neighborhood of *Haloferax volcanii fdr* (HVO_2345, *Hv*FdR) (**A**) compared to its gene homologs among 11 *Haloferax* species (**B**). Arrows represent either open reading frames (ORFs) with gene locus tag numbers or small non-coding RNAs (sncRNAs) identified in the presence (+) or presence and absence (±) of ROS [45], as indicated. DUF, domain of unknown function. HTH, helix-turn-helix. Pfam, database of protein family annotations.

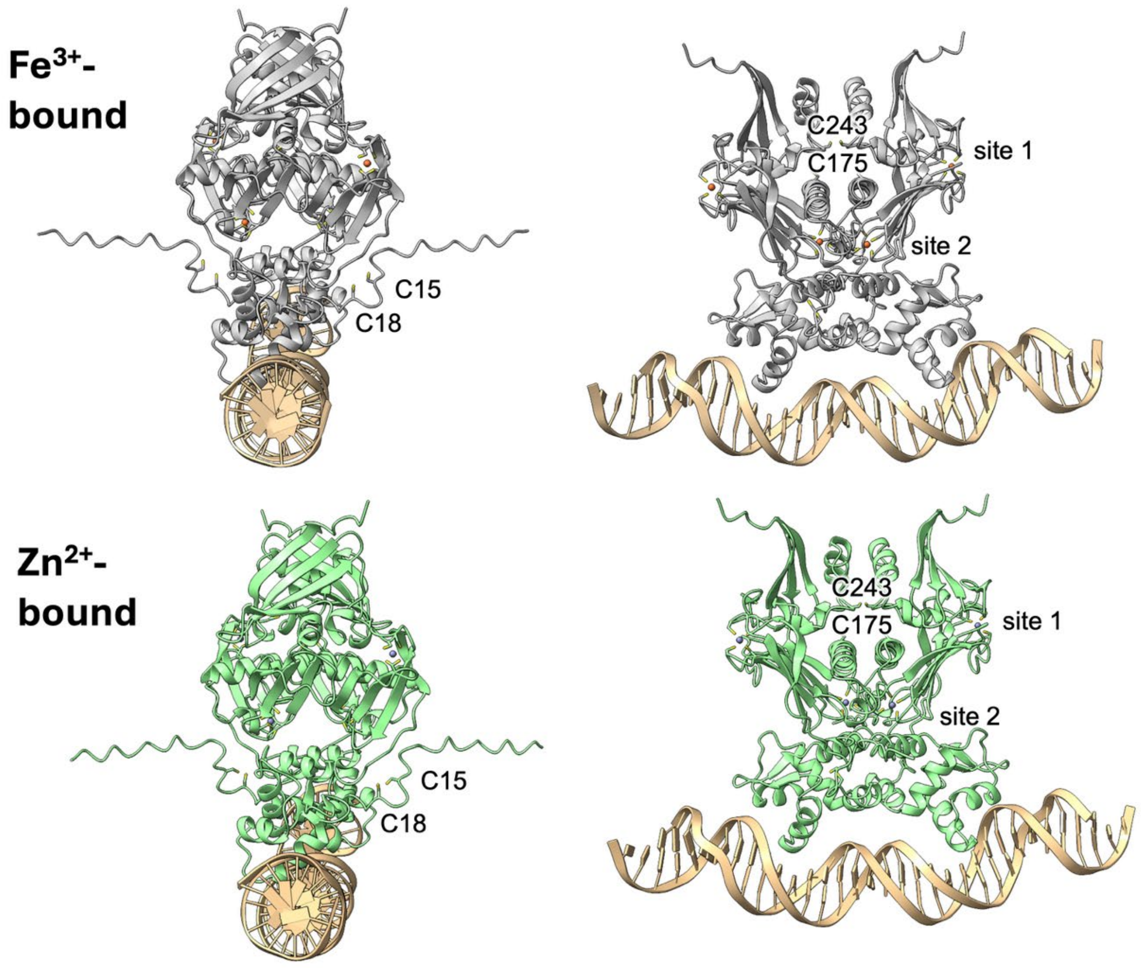


**Supplemental Figure S2.** 3D-model of HTH-domain protein HVO_2344. HVO_2344 was modeled with high confidence as a homodimer bound to double stranded DNA (tan) and 4 molecules of $Fe^{3+}$ (upper, brown spheres) or $Zn^{2+}$ (lower, blue spheres) using AlphaFold 3.0 (probability scores of ipTM = 0.83 pTM = 0.85 for both models). The 3D-models were related with a RMSD of 0.385 Å between 297 pruned atom pairs and 0.673 Å across all 302 atom pairs. dsDNA fragment 5'-AGATAGCAATCCACATTAACCCCGAATACAAA-3' (tan ribbon) was from the *fdr* promoter region. Cysteine residues predicted to coordinate metal ions at site 1 (C117, 120, 137 and 140) and site 2 (C182, 185, 210 and 213) as well as the thiol sensing cysteines (C15 and C18; C175 and 243) are indicated on the $Fe^{3+}$-bound model.

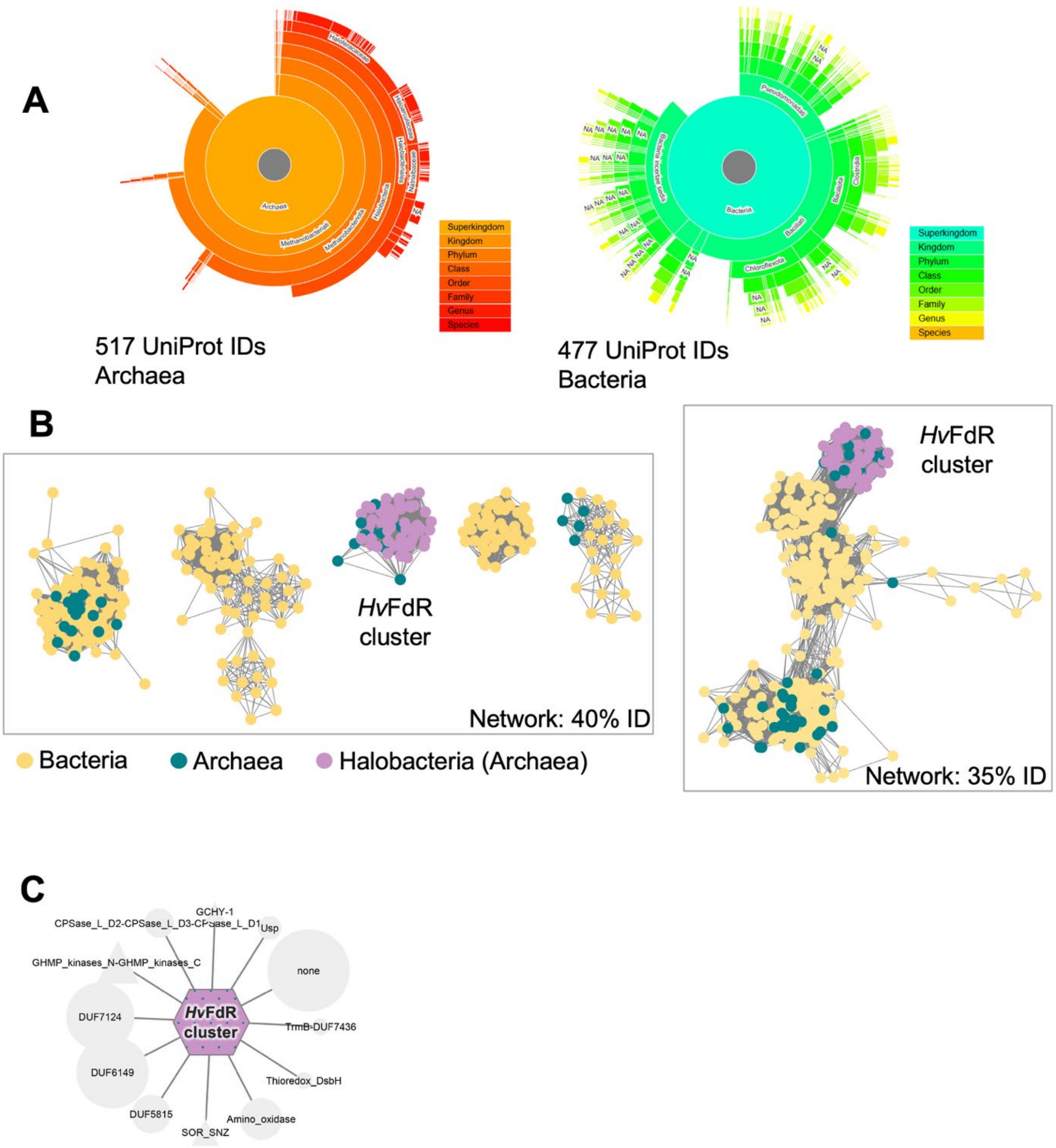


**Supplemental Figure S3.** Sequence similarity network (SNN) and genome neighborhood network (GNN) of *Hv*FdR and its homologs. Networks were generated using EFI-EST [47].

A. Phylogenetic distribution of the top 1000 protein homologs identified using *Hv*FdR (HVO_2345) protein sequence as the input for Blast and EFT-EST default parameters.
B. SNNs generated at 40% and 35% amino acid identity (ID). Alignment scores were at 81 and 65 to obtain the networks at 40 and 35% identity, respectively. Each node represents a group of protein homologs at 70% identity.
C. Pfam and DUF classification of ORFs within the GNN associated with the *Hv*FdR cluster. The cluster comprised 483 sequences that had neighboring genes. Node size in each Pfam group is proportional to the frequency of co-occurrence events. Top hits with *H. volcanii* example were: DUF6149 (HVO_2343), DUF7124 (HVO_2347), DUF5815 (HVO_2359) and PF01593 amino oxidase family (HVO_2340). None, unclassified proteins within EFT-EST including recommended DUF7979 (HVO_2346) and HTH protein (HVO_2344).

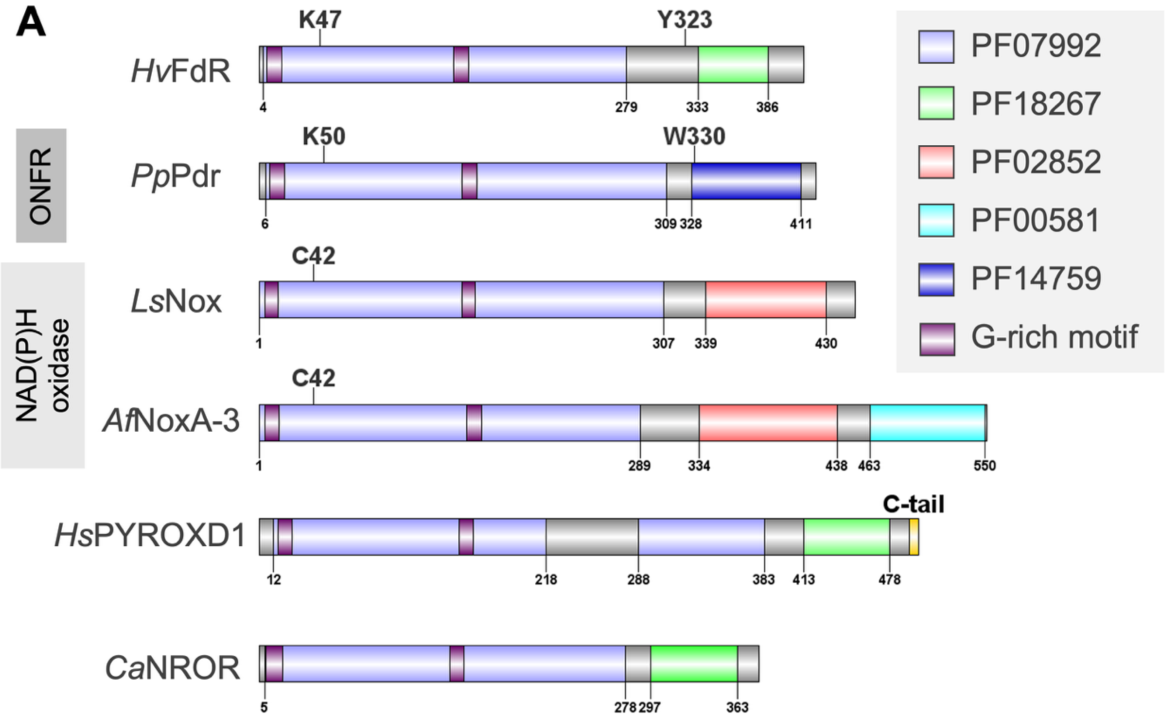


**B**

**Percent Identity Matrix**

| | | | | | | |
|---|---|---|---|---|---|---|
| 1: *Hs*PYROXD1 | 100.00 | 20.15 | 18.84 | 21.33 | 22.16 | 18.72 |
| 2: *Ls*Nox | 20.15 | 100.00 | 27.78 | 23.22 | 23.85 | 22.47 |
| 3: *Af*Nox-A3 | 18.84 | 27.78 | 100.00 | 26.22 | 26.65 | 27.00 |
| 4: *Ca*NROR | 21.33 | 23.22 | 26.22 | 100.00 | 26.50 | 21.95 |
| 5: *Hv*FdR | 22.16 | 23.85 | 26.65 | 26.50 | 100.00 | 24.57 |
| 6: *Pp*Pdr | 18.72 | 22.47 | 27.00 | 21.95 | 24.57 | 100.00 |

**Supplemental Figure S4.** Domain architecture (**A**) and percent identity matrix (**B**) of *H. volcanii Hv*FdR compared to flavin-dependent NAD(P)H oxidase, ONFR and related enzymes. Protein (Uniprot accession): *Hv*FdR (D4GWJ4), *Pseudomonas putida Pp*Pdr (P16640), *Lactobacillus sanfranciscensis Ls*Nox (Q9F1X5), *Archaeoglobus fulgidus Af*Nox-A3 (O29847), *Homo sapiens Hs*PYROXD1 (Q8WU10); *Clostridium acetobutylicum* NROR (Q9AL95). Highlighted Pfam domains include: PF07992, pyridine nucleotide-disulfide oxidoreductase domain; PF18267, rubredoxin $NAD^+$ reductase C-terminal domain; PF02852, pyridine nucleotide-disulfide oxidoreductase, dimerization domain; PF00581, rhodanese-like domain; PF14759, reductase C-terminal domain. Glycine rich motifs predicted to bind NAD(P)H and FAD (V/I-X-G-X-G-X-X-G-X-X-X-G/A) and the C-terminal tail of *Hs*PYROD1 that adopts an alpha-helical conformation upon interaction with tRNA ligase catalytic subunit RTCB [48] are also noted. Percent identity matrix, based on amino acid sequence comparison, generated using Clustal2.1.

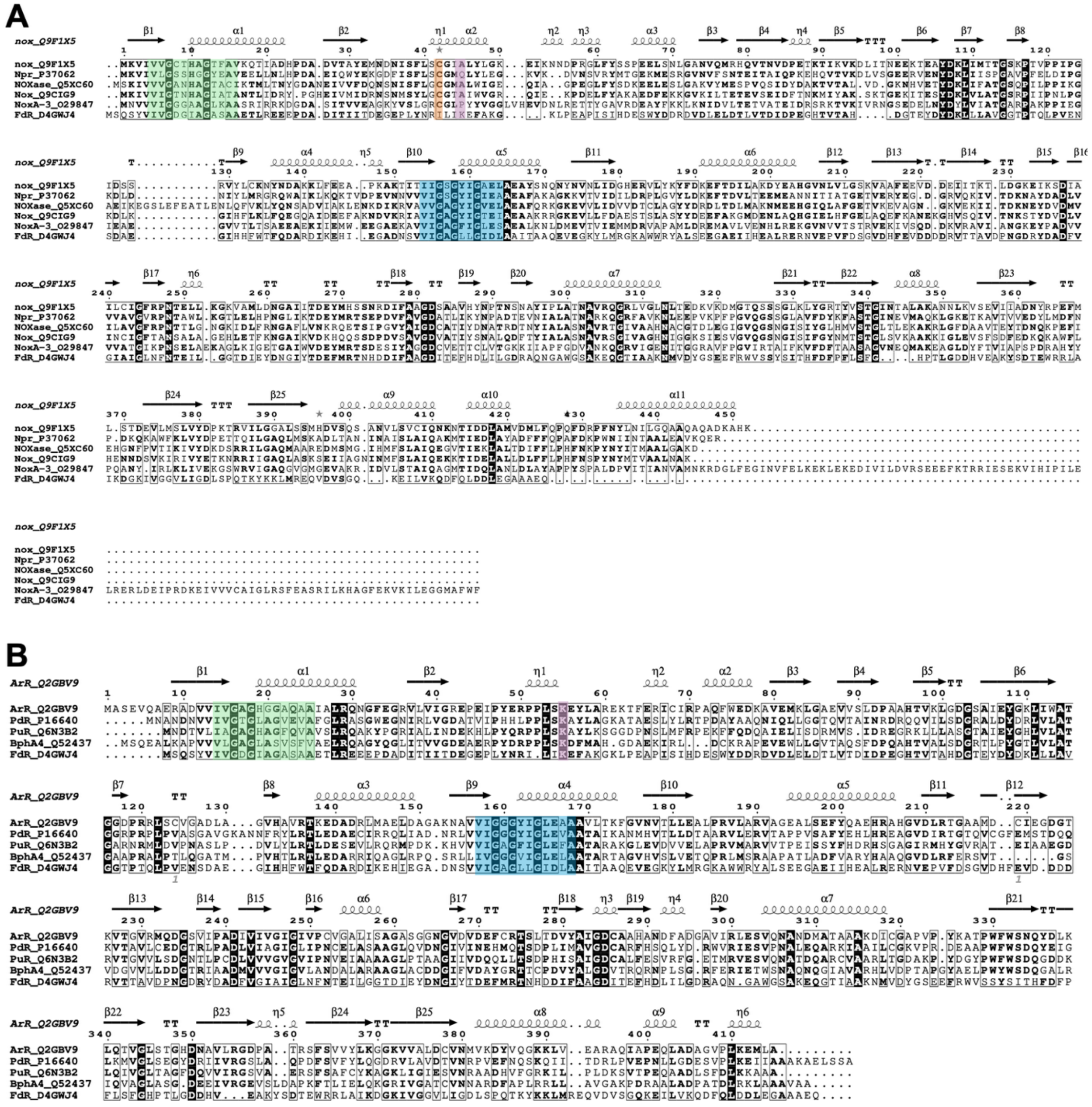


**Supplemental Figure S5.** Multiple amino acid sequence alignment of *Hv*FdR with characterized flavin-dependent NAD(P)H oxidases (**A**) and ONFRs (**B**).

A. *Hv*FdR (FdR_D4GWJ4) aligned with NAD(P)H oxidases *Lactobacillus sanfranciscensis* (Nox, Q9F1X5), *Enterococcus faecalis* (Npr, P37062), *Archaeoglobus fulgidus* DSM 4304 (Nox-A3, O29847), *Streptococcus pyogenes* (NOXase, Q5XC60), and *Lactococcus lactis* (NoxE, Q9CIG9). Redox active cysteine that aids in the four-electron oxidation of NADPH and reduction of $O_2$ to form $H_2O$ under aerobic conditions highlighted in orange. *Hv*FdR lacks cysteine residues [30, 50, 51]. *Hv*FdR K47 predicted to bind FAD is not conserved among NAD(P)H oxidases (purple). Glycine rich motifs predicted to bind NAD(P)H (green) and FAD (blue) (V/I-X-G-X-G-X-X-G-X-X-X-G/A) indicated. Multialign was used to generate the sequence alignment and imported to ESPript 3.0. *Lactobacillus sanfranciscensis* PDBe ID 2cdu was used as the input file for the secondary structure depiction. Arrows, β-strands; Coils, α-helices; η / loops, $3_{10}$ helices or loop regions; TT, tight turns connecting these elements.

B. *Hv*FdR (FdR_D4GWJ4) aligned with ONFR enzymes *Novosphingobium aromaticivorans* (ArR, Q2GBV9), *Pseudomonas* sp. KKS102 (BphA4, Q52437), *Pseudomonas putida* (Pdr, P16640), and *Rhodopseudomonas palustris* (PuR, Q6N3B2). Details as above with the following exceptions. Lysine residue (purple) predicted to be key for FAD binding based on structural analysis of ONFRs is conserved in *Hv*FdR (K47). *Novosphingobium aromaticivorans* PDBe ID 3lxd was used as the input file for the secondary structure depiction.

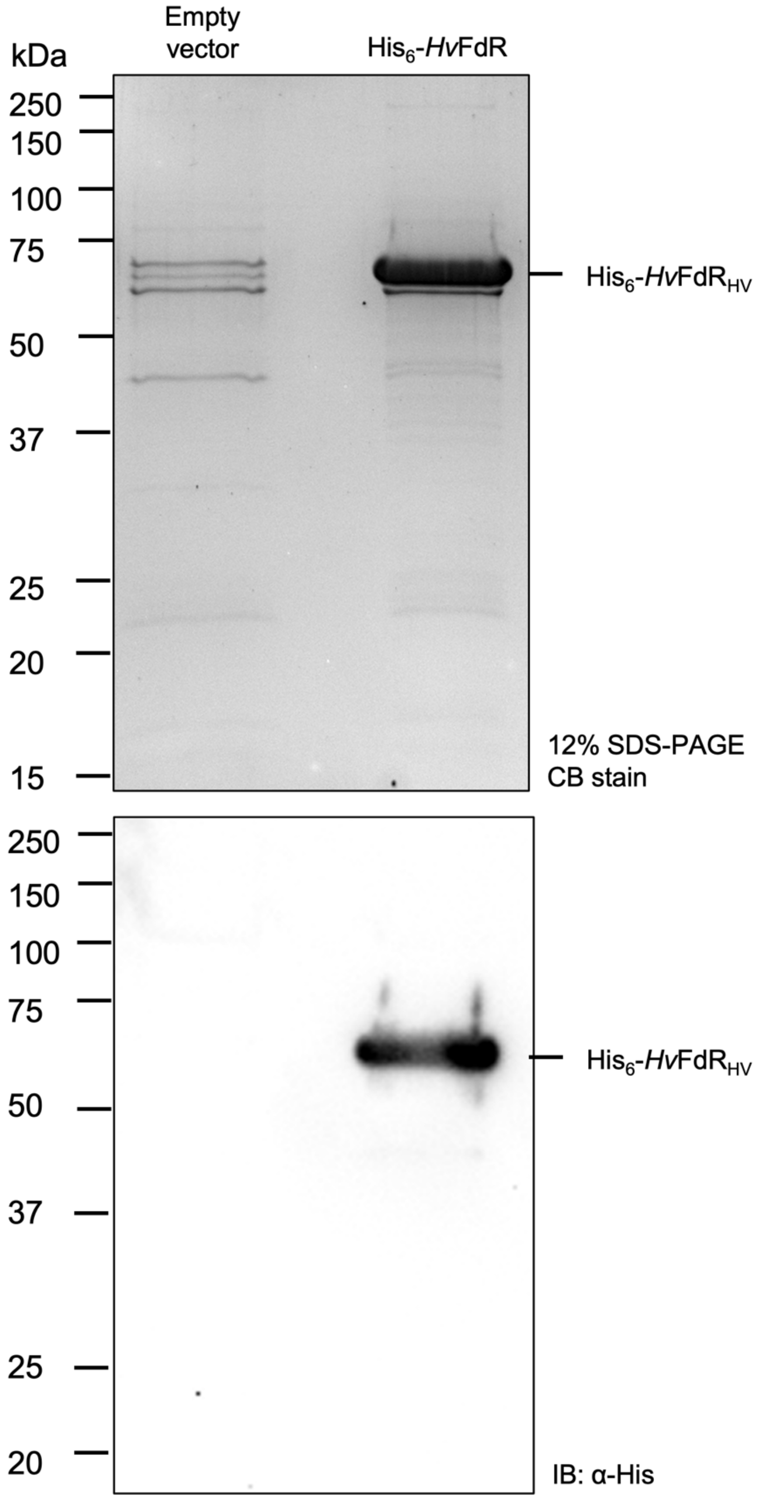


**Supplemental Figure S6.** Ni-NTA purification of His$_6$-*Hv*FdR and empty vector. *Haloferax volcanii Hv*FdR was expressed with an N-terminal His tag (His$_6$) under the control of the P2$_{rrn}$ promoter in KT05 (H1207 Δ*fdr*). Cells harboring the *Hv*FdR expression plasmid or the empty vector control were grown to stationary phase in ATCC974 rich medium supplemented with novobiocin. His$_6$-*Hv*FdR was purified by FPLC using a $Ni^{2+}$-affinity (HisTrap) column equilibrated in 20 mM HEPES (pH 7.5) containing 2 M NaCl and eluted with 500 mM imidazole. Elution fractions were analyzed by 12% SDS-PAGE coomassie blue (CB) stain (**top**) and anti-His immunoblotting (**bottom**). A total of 2.5 µg protein was loaded per lane for both analysis. The theoretical mass of His$_6$-*Hv*FdR is 47.9 kDa.

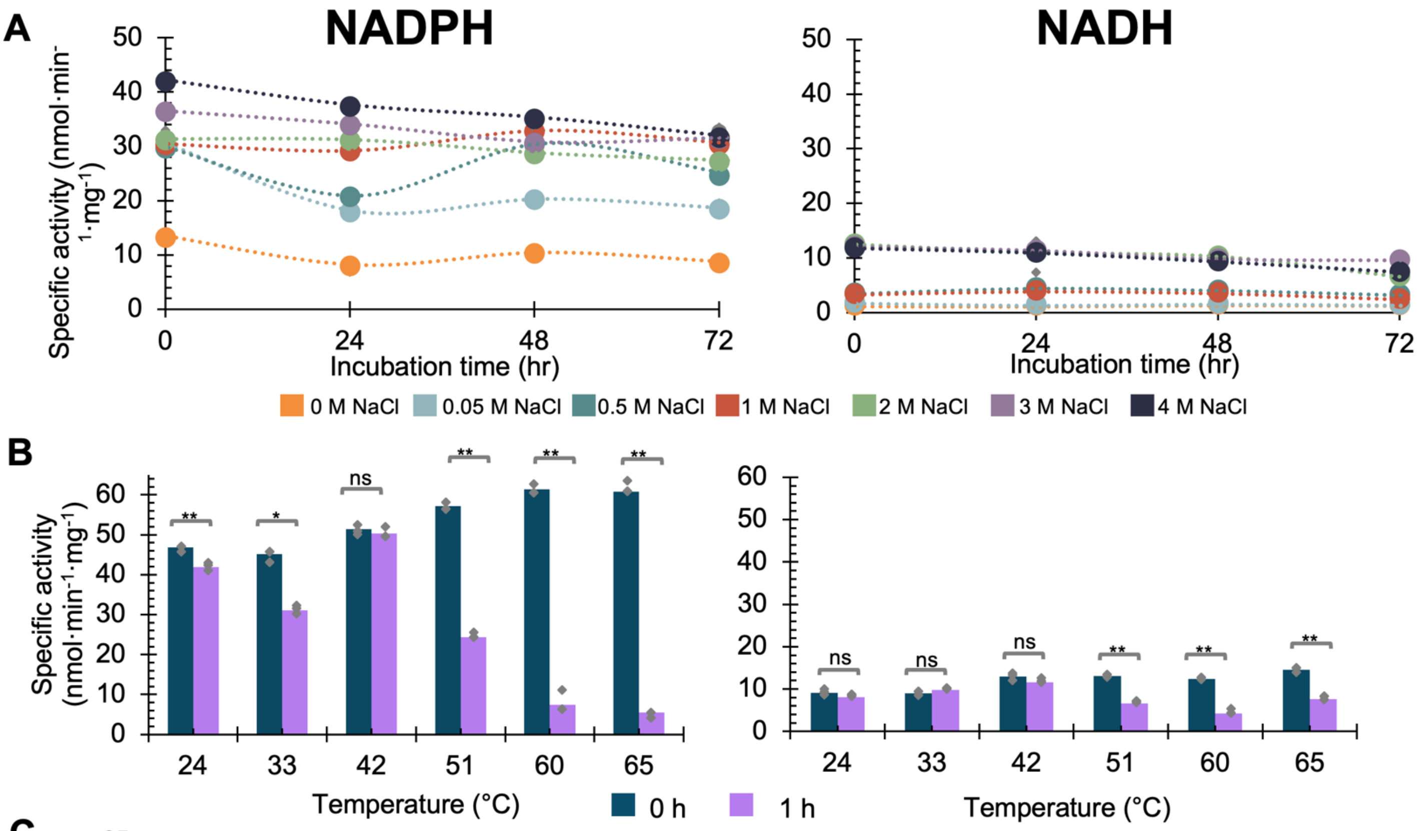


C

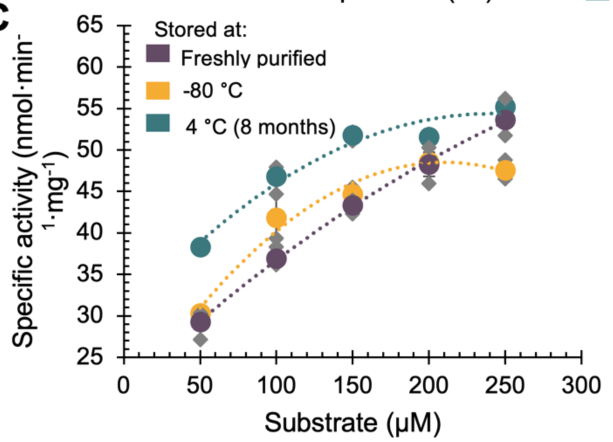


| NADH oxidase kinetic constants | His$_6$-*Hv*FdR (1.5 μM) stored at | | |
|---|---|---|---|
| | **Freshly purified** | **-80 °C** | **4 °C, 8 months** |
| $V_{max}$ (μM·s$^{-1}$) | 0.08 | 0.07 | 0.07 |
| $K_m$ (μM) | 72.5 ± 5.2 | 46.8 ± 6.7 | 31.7 ± 5.5 |
| $k_{cat}$ (s$^{-1}$) | 0.053 | 0.047 | 0.049 |
| $k_{cat}/K_m$ (M$^{-1}$ s$^{-1}$) | 736 ± 47 | 1,007 ± 122 | 1,584 ± 214 |

**Supplemental Figure S7.** Salt, temperature, and storage stability of *Hv*FdR. NADPH oxidation was monitored at absorbance 340 nm over 30 min. Specific activity (nmols·min$^{-1}$·mg$^{-1}$) is defined as nmol of NAD(P)H oxidized per min per mg *Hv*FdR.

A. Salt tolerance was tested by incubating FdR in 20 mM HEPES [pH 8.0] containing varying concentrations of NaCl (0-4 M) for 24, 48, and 72 hours at 4 °C. 0 M NaCl, orange; 0.05 M NaCl, light blue; 0.5 M NaCl, teal; 1 M NaCl, red; 2 M NaCl, green; 3 M NaCl, purple; 4 M NaCl, dark blue. A student's t-test was used to determine the statistical significance of NADPH as the electron donor between hour 0 and 72 hours: 0 M NaCl (0.001), 0.05 M NaCl (0.0001), 0.5 M NaCl (0.00005), 1 M NaCl (0.04), 2 M NaCl (0.01), 3 M NaCl (0.004), 4 M NaCl (0.0005). Circles represent the average of the three replicates and closed grey diamonds represent the individual replicates. In a total reaction volume of 150 μL, 1.5 μM of FdR and 150 μM NADPH (**left**) and 150 μM NADH (**right**) as the electron donor were added under aerobic conditions. A student's t-test was used to determine the statistical significance of NADH as the electron donor between hour 0 and 72 hours: 0 M NaCl (0.9), 0.05 M NaCl (0.09), 0.5 M NaCl (0.2), 1 M NaCl (0.01), 2 M NaCl (0.0001), 3 M NaCl (0.0001), 4 M NaCl (0.001).
B. Temperature tolerance was tested by incubating FdR for 1 h at the respective temperature and NADPH oxidation was tested at the respective temperature. For assays using NADPH (left), FdR was incubated in 20 mM HEPES [pH 8.0] with 4 M NaCl, whereas for NADH-dependent assays (right),

FdR was incubated in 20 mM MES [pH 6.0] with 3 M NaCl. 0 hr (no incubation), dark blue; 1 hr (incubation), purple. A student's t-test was used to determine the statistical significance (p-value <0.005, **; <0.05, *; ns, non-significant) of 0 hr and 1 hr when NADPH is the electron donor: 24 °C (0.0004), 33 °C (0.0076), 42 °C (0.2), 51 °C (0.0004), 60 °C (0.00003), 65 °C (0.000002). A student's t-test was used to determine the statistical significance (p-value <0.005, **; <0.05, *; ns, non-significant) of 0 hr and 1 hr when NADH is the electron donor: 24 °C (0.09), 33 °C (0.1), 42 °C (0.21), 51 °C (0.00004), 60 °C (0.0002), 65 °C (0.00009).

C. Storage conditions of FdR were tested for specific activity. Freshly purified FdR (taken from **Fig. 5**) was compared against FdR stored at −80 °C for 2 days in 20 mM HEPES, pH 7.5, 2 M NaCl, and 10% (v/v) glycerol. For long term-storage, FdR was stored in plastic tubes at 4°C for 8 months in 20 mM HEPES, pH 7.5, and 2 M NaCl under aerobic conditions. Freshly purified, purple; Stored at −80 °C, orange; Stored at 4 °C (8 months), teal (**Left**). Freshly purified *Hv*FdR (1.5 μM) used in this data analysis was the same preparation shown in Figure 5. (**Right**) Steady-state kinetic parameters of varying storage conditions of wild-type FdR with NADPH as the electron donor (substrate) under aerobic conditions. $V_{max}$ represents the maximal reaction velocity (μM·$s^{-1}$) achieved at substrate saturation. $K_m$ denotes the Michaelis constant (μM), reflecting the substrate concentration at half-maximal velocity. $k_{cat}$ ($s^{-1}$) is the catalytic turnover number, calculated as $V_{max}$ normalized to enzyme concentration. $k_{cat}/K_m$ ($M^{-1}$ $s^{-1}$) represents catalytic efficiency. Values are reported as mean ± standard deviation.

**Salt tolerance of *Hv*FdR.** To assess salt tolerance, freshly prepared $His_6$-*Hv*FdR was incubated at 4 °C for 0, 24, 48 and 72 h in buffers supplemented with 0 to 4 M NaCl. NAD(P)H oxidase activity was subsequently measured at 42 °C in the corresponding salt concentrations (**Supplemental Fig. S7A**). During the initial 24-h incubation period, $His_6$-*Hv*FdR was found to have reduced activity when stored in low salt buffer (≤ 0.05 M NaCl), irrespective of the electron donor. At the higher salt concentrations, $His_6$-*Hv*FdR remained stable in its enzymatic activity following the 24-h incubation period. After storage for 48 or 72 h in the high salt buffers, $His_6$-*Hv*FdR displayed a 1.2-fold and 1.3-fold reduction in NADPH oxidase activity at 4 M NaCl (**Supplemental Fig. S7A, left and right**). Thus, $His_6$-*Hv*FdR appeared relatively stable and active when stored in a high salt compared to low salt buffer.

**Thermal response of *Hv*FdR.** To assess the effect of temperature on *Hv*FdR activity following prolonged heat exposure, $His_6$-*Hv*FdR was incubated for 1 h at temperatures ranging from 24 to 65 °C, after which enzyme activity was measured at the corresponding incubation temperature (**Supplemental Fig. S7B**). At the lower temperatures (24, 33 and 42 °C), $His_6$-*Hv*FdR was found to retain 68-97% activity with NADPH and 88-108% activity with NADH. The higher initial activity observed at 42 °C compared with 24 °C and 33 °C at 0 h likely reflects an increase in the catalytic rate at elevated assay temperatures or rapid thermal equilibration during measurement rather than enhanced thermal stability. By contrast, a 2 to 11-fold reduction in activity was observed for both electron donors when the enzyme was incubated for 1 h at 51, 60 or 65 °C. The 1 h incubation at 60 °C or above resulted in visible protein precipitation. These findings indicate that long-term exposure to elevated temperatures compromises the stability and catalytic activity of the $His_6$-*Hv*FdR enzyme.

**Storage of *Hv*FdR.** To evaluate the impact of freeze-thaw and long-term storage, $His_6$-*Hv*FdR was evaluated for its NADPH oxidase activity after (i) directly frozen by storage at −80 °C for 2 days and (ii) storage at 4°C for 8 months under aerobic conditions (**Supplemental Fig. S7C**). A reduction in activity was observed after storage at −80 °C compared to freshly purified $His_6$-*Hv*FdR, where specific activity reduced 1.1-fold when assayed at 250 μM NADPH. Activity remined similar at 250 μM NADPH between freshly purified and $His_6$-*Hv*FdR stored at 4 °C. While freshly purified $His_6$-*Hv*FdR exhibited the highest turnover rate, it also displayed a lower affinity for NADPH, resulting in reduced catalytic efficiency. In contrast, $His_6$-*Hv*FdR stored at 4 °C and -80 °C decreased in turnover and increased in substrate affinity, resulting in the highest overall efficiency. Collectively, these results suggest $His_6$-*Hv*FdR may alter structural integrity or conformational states during the long-term storage and freeze-thaw conditions examined.
